# Genome-wide mapping of Cas9-induced sister chromatid exchange across single and over 200 genomic targets

**DOI:** 10.64898/2026.05.13.724813

**Authors:** Peter Chovanec, Shiyang He, Yi Yin

## Abstract

CRISPR/Cas9-induced DNA double-strand breaks (DSBs) are widely used for genome engineering, yet their capacity to provoke sister chromatid exchange (SCE) and associated genome instability remains incompletely understood, in part because exchanges between identical sister chromatids leave no sequence change and are invisible to conventional whole-genome sequencing. Using sci-L3-Strand-seq, a scalable single-cell template-strand sequencing platform for genome-wide SCE mapping, we quantified strand-switch events following targeted Cas9 cleavage. Cas9 targeting of a unique genomic locus led to strong local enrichment of SCE at the break site, reaching up to 41% in the same cell cycle and 17% in the subsequent division, indicating that DSB repair frequently engages non-local inter-sister repair. Extending the assay to 237 repetitive Alu targets using one Alu-specific sgRNA revealed mild and marginally statistically significant enrichment of on-target SCE across wild-type Cas9 and nickase variants (D10A and H840A). However, restricting analysis to a subset of cells with elevated SCE burden (>8 SCEs per cell) uncovered significant enrichment at programmed cut sites, suggesting that recombination at repetitive loci is conditionally engaged in highly recombinogenic cells. Reciprocal daughter-cell pair analysis further revealed large-scale structural alterations on chromosomes with induced SCEs and at SCE junctions, indicating that Cas9-induced SCEs do not universally reflect error-free homologous recombination. Disruption of DNA repair genes at the cut site, including LIG3, LIG4, XRCC1, and XRCC4, did not measurably alter SCE frequency per cell, consistent with delayed functional loss following editing and, in the case of essential genes such as LIG3, selection against disruptive alleles. Together, these findings demonstrate that Cas9-induced DSBs are potent local triggers of SCE at unique loci, while at repetitive regions SCE is more context-dependent and can be associated with structural alterations, highlighting the influence of lesion type and genomic context on recombination outcomes during genome editing.

## Introduction

CRISPR/Cas9 genome editing relies on the induction of site-specific DNA double-strand breaks (DSBs) to trigger endogenous repair pathways (Cong et al., 2013; Ran et al., 2013). The two principal repair outcomes, non-homologous end joining (NHEJ) and homologous recombination (HR), have been extensively characterized at the sequence level in wild-type cells and with DNA repair gene perturbations (Chen et al., 2019; Hussmann et al., 2021; Shen et al., 2018): NHEJ generates small insertions and deletions at the break site, while HR can incorporate an exogenous donor template for precise editing (Richardson et al., 2016). However, this framework captures only the local and mutational fraction of the repair landscape. For non-local mutational outcomes, the structural consequences of Cas9 cutting extend well beyond the small indels detected by standard amplicon sequencing. Large deletions spanning kilobases have been documented at on-target sites (Kosicki et al., 2018), and Cas9-induced DSBs can trigger chromothripsis through micronucleus formation and chromosome bridge breakage (Leibowitz et al., 2021). Chromosome loss and segmental aneuploidies have been observed following Cas9 editing in human embryos, attributed to the isochromatid nature of the break (Zuccaro et al., 2020). At the cytogenetic level, chromosomal instability and rearrangements in edited cancer cell lines are frequently missed by PCR-based genotyping (Regan et al., 2025). These observations establish that Cas9 DSBs can produce a spectrum of structural alterations, yet the relationship between SCE and these larger-scale events has not been examined in the same experimental framework.

In this study, we assay what is likely the most frequent non-local outcome (Fig.S1A) induced by Cas9: sister chromatid exchange (SCE). Here, we operationally define SCE as a reciprocal strand switch between sister chromatids, without implying a single mechanistic pathway. SCE can therefore arise from inter-sister homologous recombination (HR), double non-homologous end joining (“dNHEJ”) across two isochromatid DSBs, or other non-local repair processes. This distinction is particularly relevant for Cas9, which typically cleaves both sister chromatids at the same locus and thereby generates the precise isochromatid substrate on which dNHEJ can act. If only one sister is cleaved, the result is a two-ended, post-replicative DSB with an intact sister available as a repair template, an ideal substrate for HR-mediated SCE. More commonly, however, Cas9 can generate isochromatid DSBs through two routes: by cleaving both post-replicative sisters independently, or by cutting the unreplicated template ahead of the fork, such that the break is replicated into DSBs on both daughters. In either case, ultimately four broken ends are produced, although not necessarily simultaneously, and neither sister is intact (Zuccaro et al., 2020). The repair outcome then depends on how the four ends are resolved: cis joining (each sister re-ligated to its own ends) restores the original configuration, while trans joining between sisters (the proximal end of one sister joined to the distal end of the other) produces a strand switch operationally indistinguishable from HR-mediated SCE. Whether Cas9-induced SCE at isochromatid breaks arises through HR, through trans end joining (dNHEJ), or through a mixture of both has not been determined. In all cases, exchanges between identical sisters are genetically silent and invisible to conventional whole-genome sequencing, and have therefore been largely overlooked as an outcome of Cas9 editing. Whether a single targeted DSB efficiently triggers local SCE, how SCE frequency scales when hundreds of sites are cut simultaneously, and whether Cas9-induced SCEs are uniformly error-free remain open questions.

Template-strand sequencing methods, including Strand-seq and its scalable derivatives OP-Strand-seq and sci-L3-Strand-seq, overcome the fundamental limitation that SCE between identical sisters leaves no sequence change (Chovanec et al., 2026b; Falconer et al., 2012; Hanlon et al., 2022; van Wietmarschen et al., 2018; van Wietmarschen and Lansdorp, 2016). By incorporating BrdU into newly synthesized DNA during one cell cycle, selectively degrading the BrdU-labeled nascent strands, and sequencing only parental template strands, these methods map the Watson- and Crick-oriented parental strands inherited by each daughter cell. SCEs are detected as switches in parental strand orientation along a chromosome, providing a direct genome-wide readout of non-local repair events in single cells independent of any sequence alteration (Fig.S1A). Furthermore, the high throughput of sci-L3-Strand-seq enables recovery of reciprocal daughter-cell pairs (RDCPs), in which both products of a single mitotic division are captured, allowing direct assessment of whether SCE events are reciprocal and copy-neutral or accompanied by structural alterations.

Here, we use sci-L3-Strand-seq to map Cas9-induced SCE at single-cell resolution across two experimental regimes: a single unique-locus cut and simultaneous cleavage at 237 repetitive targets. We show that a single Cas9 DSB at a unique site is a potent local trigger of SCE. At repetitive loci, on-target SCE enrichment is modest in bulk but becomes significant in a subset of cells with elevated recombination burden. RDCP analysis reveals that Cas9-induced SCEs are not uniformly benign: structural alterations at exchange junctions indicate that repair of two-ended, isochromatid Cas9 DSBs frequently deviates from error-free HR, in contrast to the clean reciprocal outcomes observed for spontaneous, replication-born SCEs in a companion study (Chovanec et al., 2026a). Wild-type Cas9 and nickase variants (D10A, H840A) are compared, and the consequences of disrupting DNA repair genes at the cut site are assessed. Together, these results provide the first genome-wide, single-cell map of Cas9-induced SCE and reveal that the isochromatid geometry of Cas9 breaks creates a distinct recombination landscape with both error-free and mutagenic outcomes.

## Results

### Cas9 duplex cleavage, but not nicking, can induce durable cell-cycle arrest at single and repetitive targets

To define how Cas9-induced lesions impact cell-cycle progression and SCE, we used recombinant Cas9 ribonucleoprotein (RNP) complexes (IDT), a widely adopted delivery format for genome editing that enables efficient generation of monoclonal knockouts as well as high-efficiency (>90%) polyclonal knockouts for bulk perturbation experiments. We compared four single-site sgRNA, each targeting a unique genomic locus, with a multi-target sgRNA recognizing 237 repetitive sites. Introduction of duplex-cleaving Cas9 at either a single site or across repetitive targets led to a pronounced G1/S arrest within 24 hours of delivery, indicating that even a single DSB is sufficient to trigger a robust checkpoint response, irrespective of whether the targeted locus resides in an essential gene (LIG3) or a non-essential gene (LIG4). Notably, this arrest persisted after Cas9 removal only in cells targeted with the repetitive sgRNA, consistent with sustained or cumulative DNA damage at multiple genomic sites (**Fig.1A-B, Fig.S1A**).

**Figure 1.**
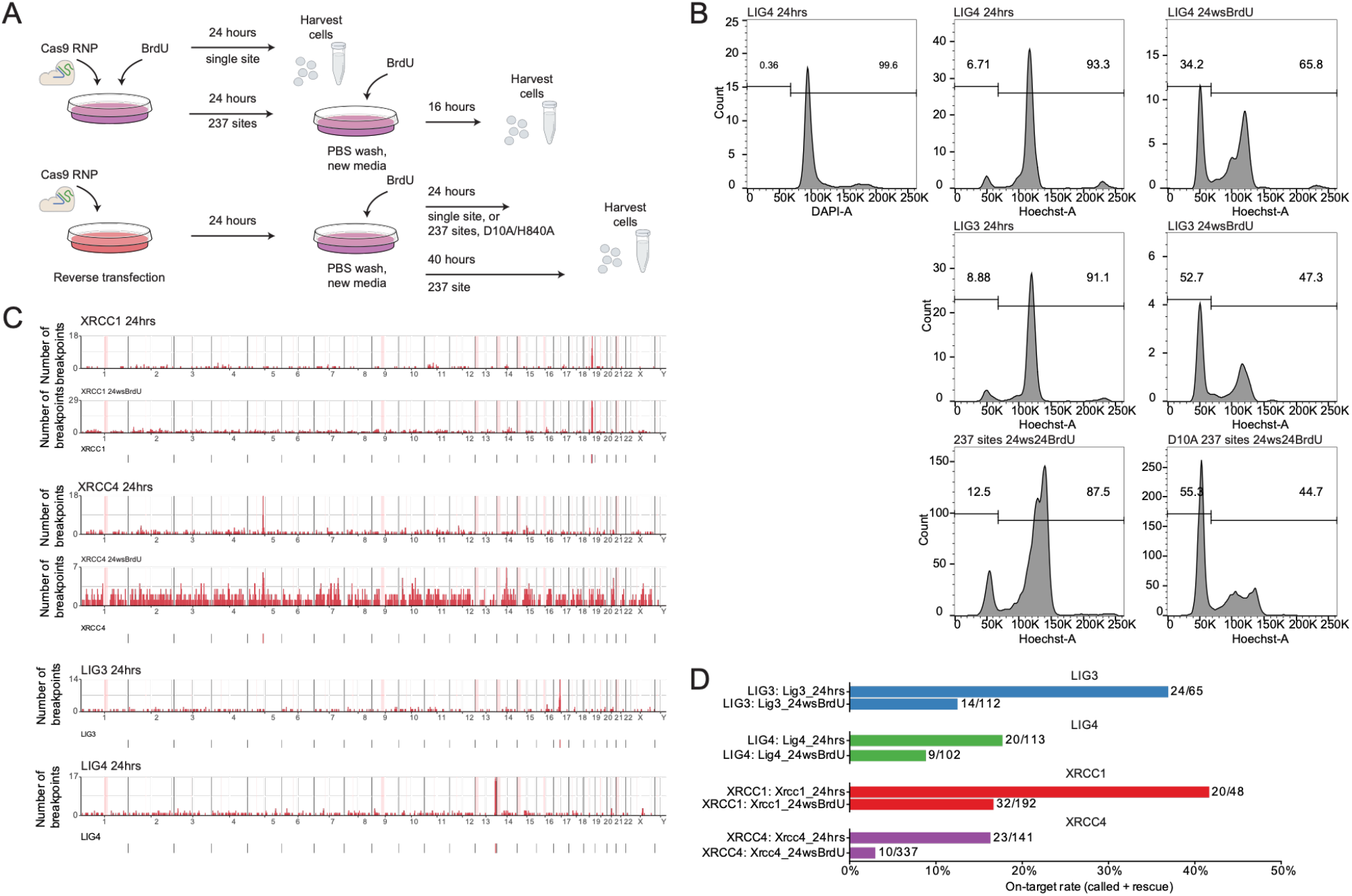
Experimental design, cell cycle perturbation and genome-wide pileup of both spontaneous and Cas9-induced on-target SCE events. **A**. Experimental design. BrdU was incorporated to enable SCE mapping by sci-L3-Strand-seq, either during the same 24-hour window as Cas9 RNP treatment or during a 24-hour period following Cas9 RNP washout. Wild-type Cas9 was used unless otherwise indicated for nickase variants. **B**. FACS analysis of cell-cycle progression following Cas9 RNP treatment. “24hrs” denotes concurrent Cas9 RNP treatment and BrdU labeling, whereas “24 ws BrdU” indicates BrdU labeling after Cas9 RNP washout. Each of the four single-targeting sgRNAs targets a distinct gene, as indicated. For the “LIG4 24hrs” condition, the small fraction of cells that successfully cycled into the subsequent G1 (6.71%) showed approximately half the Hoechst signal (centered at ∼50K) compared with the DAPI signal (centered at ∼100K), consistent with completion of cell division. **C**. Genome-wide pileup plots for both spontaneous and Cas9-induced on-target SCE events. The targeting site is indicated in the panel with the gene name and a vertical red bar. For sgRNAs targeting XRCC1 and XRCC4, SCEs from both the first division during RNP treatment and the post-washout division are shown. For LIG3 and LIG4, SCEs from the first division are shown. Additional pileups, including comparisons between breakpointR-based and rescued SCE calls, are provided in **Fig.S1B**. **D**. Quantification of on-target SCE events for the four single-site sgRNAs in the first division (24 hrs) and in the subsequent division following RNP washout (24 ws).

Cas9 nickases, in which one of the two nuclease domains is inactivated (D10A or H840A), were developed to reduce off-target mutagenesis by converting DSBs into a single-strand nicks (Mali et al., 2013; Ran et al., 2013). Although paired nickases with offset guide RNAs can achieve on-target editing with improved specificity, individual nicks are not inherently inert, as they can be converted into DSBs by replication fork run-off or coincident nicking of the opposite strand. Given that only duplex cleavage at repetitive sequences induced a persistent cell cycle arrest after washout, we tested whether nickase-induced lesions at the same sites had a similar effect. As shown in **Fig.1B**, neither D10A nor H840A nickase variants measurably perturbed cell-cycle progression following washout. These findings indicate that duplex DNA cleavage, particularly when distributed across repetitive loci, imposes a durable cell-cycle barrier, whereas single-strand nicking is largely tolerated under these conditions.

### A single Cas9 DSB induces potent local SCE

We next examined the genomic distribution of Cas9-induced and background spontaneous SCEs using genome-wide pileup analysis (**Fig.1C, Fig.S1B**), in which we plot the total number of SCEs detected across all single cells within each 1 Mb window. Across all four sgRNAs targeting individual genomic loci, SCEs were strongly enriched at the programmed cut sites, demonstrating robust local induction of strand exchange. This enrichment was most pronounced in the first cell division following RNP delivery and was reduced 24 hours after washout. To account for low-coverage cells in which discrete breakpoints could not be confidently called, we performed a complementary analysis by quantifying strand-specific read imbalance within 10 Mb flanking the cut site, which similarly revealed clear evidence of SCE at the targeted loci. We define “on-target” events as the union of SCEs directly called by breakpointR and events recovered by this strand-state rescue approach. Across the single-site conditions, 30–92% of on-target events were directly called by breakpointR, typically >70%, with the exception of LIG4, whose proximity to the telomere makes breakpointR calling more difficult. The remaining on-target events were recovered by strand-state rescue, primarily in cells with lower per-cell sequencing depth. On-target enrichment remained robust when rescued calls were excluded (**Fig.S1B**).

As summarized in **Fig.1D**, the combined on-target SCE rate reached up to 41% in the first division and 17% after washout. Overall, the on-target rate declined by approximately half following washout, with the largest reduction of 74%, indicating that Cas9-induced SCE is most efficiently captured immediately after cleavage and diminishes in subsequent cell cycles. Notably, while Cas9 RNP delivery achieves high bulk editing efficiencies (∼90%), this likely reflects multiple rounds of cleavage and repair at the same locus; consequently, the observed SCE frequencies should not be interpreted as a proxy for the final error-free HR or for knockout efficiency. Rather, the SCE rate reported here reflects the fraction of Cas9-induced breaks repaired through intersister exchange rather than cis repair on the same chromatid.

### Cas9 targeting of DNA repair gene loci did not immediately alter overall SCE frequency per cell

Given the delayed functional loss following Cas9-mediated gene disruption, we do not expect spontaneous SCE to increase as a result of sgRNA-directed cutting within DNA repair genes over the timeframe of these experiments. Even biallelic frameshift mutations do not immediately eliminate the pre-existing protein pool, which must be diluted through turnover and cell division before a functional deficit can emerge. Consistent with this expectation, genome-wide SCE counts per cell were indistinguishable across all four gene-targeting conditions (LIG3, LIG4, XRCC1, XRCC4) at the 24-hour simultaneous labeling timepoint with all showing a median of 3 SCEs per cell, comparable to the wild-type with Cas9 cuts (Chovanec et al., 2026b) and Lig4 washout controls (pairwise Wilcoxon p > 0.09 for all 24-hour comparisons among single-site conditions, **Fig.S1C** and **Tab.S1**). This contrasts with stable XRCC1 knockout, which increases spontaneous SCE approximately five-fold in our companion study (Chovanec *et al*., 2026a). One additional complication is that, although Cas9 RNP achieves >90% knockout efficiency in bulk assays and is therefore used as a substitute for siRNA, sorting BrdU-labeled cells in the subsequent G1 enriches for cells that escaped frameshift editing, particularly for essential genes; thus, 100% knockout in 90% of cells is not equivalent to 90% knockdown in every cell, representing a unique challenge for single-cell assays.

The LIG3 locus provides an informative example. As an essential gene, complete loss of LIG3 is expected to impair cell viability. In the 24-hour simultaneous labeling condition, >91% of recovered cells were in their first division following Cas9 delivery, and we observed no reduction in nuclei recovery, indicating minimal immediate selection. In contrast, in the washout condition, where labeling occurred in the second cycle, nuclei yield was markedly reduced, although the fraction of surviving cells progressing into the subsequent G1 was high. This pattern is consistent with selective loss of cells carrying biallelic loss-of-function alleles in the second cell cycle. The recovered cells, including those with on-target SCE at the LIG3 locus, likely retained functional LIG3, either through error-free repair (including SCE itself, which preserves sequence) or in-frame edits.

More broadly, although Cas9 RNP delivery achieves high bulk editing efficiencies by amplicon sequencing, this does not equate to functional equivalence with siRNA-mediated knockdown in single-cell assays. A population in which ∼90% of cells carry biallelic knockout is not equivalent to one in which all cells experience partial depletion. For essential genes in particular, single-cell readouts are inherently biased toward the surviving, functionally intact fraction, potentially obscuring the phenotype of interest.

### On-target SCE enrichment at repetitive loci is mild but concentrated in high-SCE cells

We then asked whether SCE is enriched at the 237 repetitive target sites following Cas9 cleavage. To test this, we compared the number of SCE breakpoints falling within regions of interest (ROIs) flanking programmed cut sites against the number expected from size-matched random genomic intervals, using a Fisher’s exact test (**Tab. 1**). Across all cells, duplex Cas9 cleavage at 237 repetitive targets produced modest on-target SCE enrichment. The strongest signal came from the post-washout BrdU-labeled condition (“237cuts_24ws_BrdU”), which yielded a 1.20-fold enrichment (p = 4.89 x 10E-55, n = 2,547 cells), with an average of 6.13 on-target SCE breakpoints per cell. Adding Cas9 RNP and BrdU labeling simultaneously (“237cuts_24hrs”) showed a similar fold enrichment of 1.10 (p = 0.02, n = 214), with 5.72 on-target SCEs per cell. By contrast, the earlier WT237 experiment showed no significant enrichment (1.11-fold, p = 0.19, n = 74), likely reflecting the smaller sample size. The D10A nickase produced a marginally significant enrichment (1.09-fold, p = 0.01, n = 330), while the H840A nickase showed no enrichment (1.02-fold, p = 0.37, n = 757). These results indicate that duplex cleavage at repetitive loci induces detectable but mild on-target SCE enrichment in the bulk population, while single-strand nicking produces little or no measurable effect.

**Tab 1.**
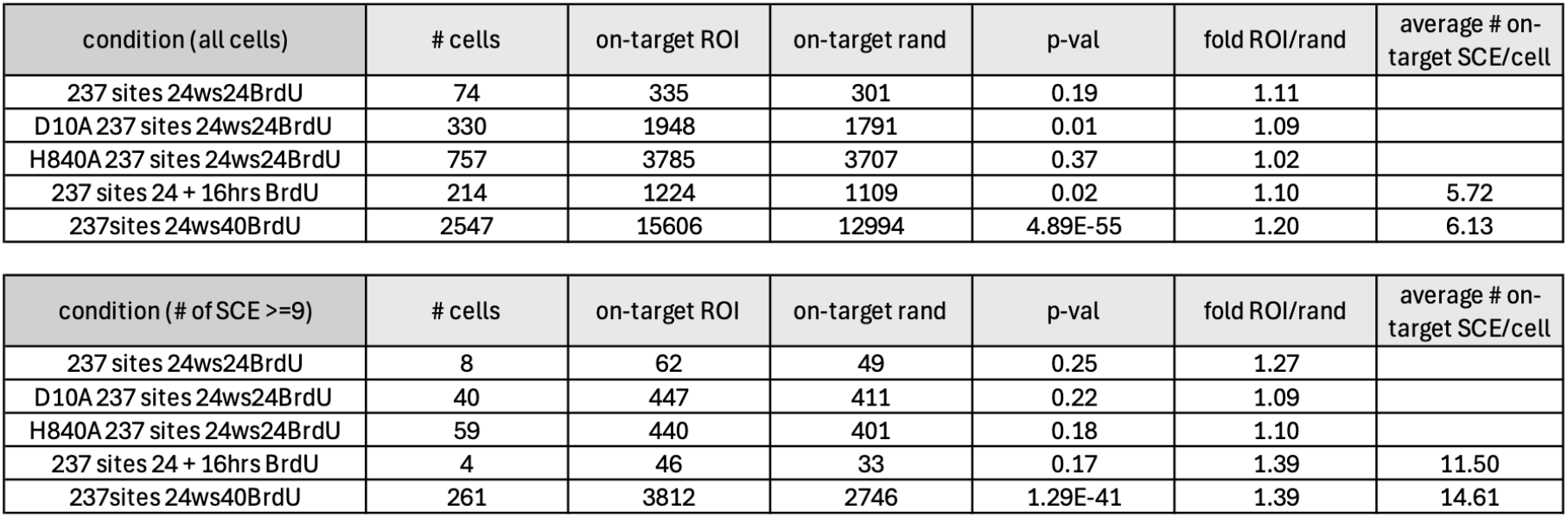
Enrichment analysis for the 237-site targeting experiment, comparing the number of SCEs falling within sgRNA-targeted sites (ROI: regions of interest) with randomly sampled sites (“rand”). “24ws24BrdU” refers to conditions in which cells were treated with Cas9 RNP (or the indicated nickase variants, D10A or H840A) for 24 hours, followed by washout and BrdU labeling for the subsequent 24 hours (or 40 hours, in the case of 24ws40BrdU to allow more cells to progress into the next G1). “24 + 16hrs BrdU” refers to the condition in which Cas9 RNP and BrdU were added simultaneously; after 24 hours of Cas9 RNP treatment, BrdU labeling was continued for an additional 16 hours (a total of 40 hours of BrdU).

A stronger pattern emerged when analysis was restricted to cells with significantly elevated SCE count (>=9 SCE per cell). This threshold represents a 2-3 fold increase above the median of 3-4 SCE in non-cut wild-type cells (Chovanec et al., 2026b), and approximately two-fold above the median for the respective Cas9-treated populations, in which the median SCE per cell does not increase beyond 4 despite the introduction of up to 237 DSBs. In the post-washout BrdU condition, this high-SCE subset (n = 261 cells) showed 1.39-fold enrichment (p = 1.29 x 10E-41), with an average of 14.61 on-target SCEs per cell, compared to 6.13 in the full population. The simultaneous 24-hour condition, though limited to 4 cells, showed a comparable 1.39-fold enrichment with 11.50 on-target SCEs per cell. That the bulk median SCE remains unchanged despite 237 concurrent DSBs, while a minority of cells shows both elevated total SCE and concentrated on-target enrichment, suggests that most cells resolve Cas9-induced breaks through non-exchange, *cis* repair, and that inter-sister exchange is conditionally engaged in a SCE-permissive subpopulation. The WT237 (n = 8) and nickase conditions showed no significant enrichment in this subset, consistent with either insufficient statistical power or a genuine absence of on-target SCE in high-SCE cells following nicking. Relaxing the threshold to >=8 SCE per cell yielded qualitatively similar trends, particularly for the 24-hour condition (1.33-fold, p = 0.06, n = 10) and the post-washout condition (1.35-fold, p = 7.92 x 10E-45, n = 395).

Together, these data indicate that Cas9-induced SCE at repetitive targets is not uniformly distributed across the cell population. A subset of cells with elevated SCE burden shows both more on-target breaks (∼2-fold) and a higher fold enrichment of SCE at cut sites relative to random loci (1.39 vs 1.10-1.20 in the bulk), indicating that these cells not only sustain more Cas9 cleavage events but also resolve a greater fraction of them through inter-sister exchange. The high-SCE subset did not show evidence of lower sequencing quality, based on either sequencing coverage (p=0.13) or background SCE levels (p=0.17). However, we cannot distinguish whether the elevated on-target SCE reflects a cell-cycle state more permissive to both cutting and recombination, or stochastic variation in RNP uptake coupled with a recombination-prone chromatin environment.

### RDCP analysis reveals large-scale SVs on chromosomes with induced SCE, as well as structural alterations at Cas9-induced SCE junctions

Reciprocal daughter cell pair (RDCP) analysis enables direct recovery of all four chromatids following replication, providing a stringent readout of SCE outcomes and associated structural changes. Leveraging the throughput of sci-L3-Strand-seq, we previously showed that spontaneous SCEs in XRCC1^−^/^−^ cells are uniformly reciprocal and copy-neutral. This framework allows us to test whether Cas9 RNP-induced lesions similarly resolve without gross rearrangement. Given the limited sampling achievable with single targeting sites, we focused RDCP analysis on cells targeted with the 237-site sgRNA. We identified one RDCP under simultaneous labeling conditions, containing two reciprocal SCE events (**Fig.2A**). Following washout, we recovered 11 RDCPs, of which four exhibited associated structural variants (SVs) (**Fig.2B**). Across these four pairs, 14 of 26 events were accompanied by SVs, often arising across two consecutive cell divisions, whereas 12 events were reciprocal and copy-neutral. The remaining seven RDCPs comprised a total of 24 exclusively reciprocal, copy-neutral SCEs (**Fig.S2**). Reconstruction of repair outcomes is detailed in **Tab.S2**. Our interpretation is further validated by haplotype-aware analysis in **Fig.S3**. Overall, 14 of 50 SCE events were coupled with SVs, in contrast to 87 of 87 spontaneous SCE events in XRCC1^−^/^−^ cells, which were uniformly reciprocal and copy-neutral in our companion study (Chovanec et al., 2026a). Additionally, we observed structural alterations at Cas9-induced SCE junctions consistent with the resolution of under-replicated regions (URR) or replication terminal zones as SCE (**Fig.2B**, Pair 4, **Fig.S3D**). The mechanistic implications of these observations are explored further in the Discussion. We summarize RDCP counts, copy-neutral SCE counts, and SV-associated events across all conditions in **Tab.S3**.

**Figure 2.**
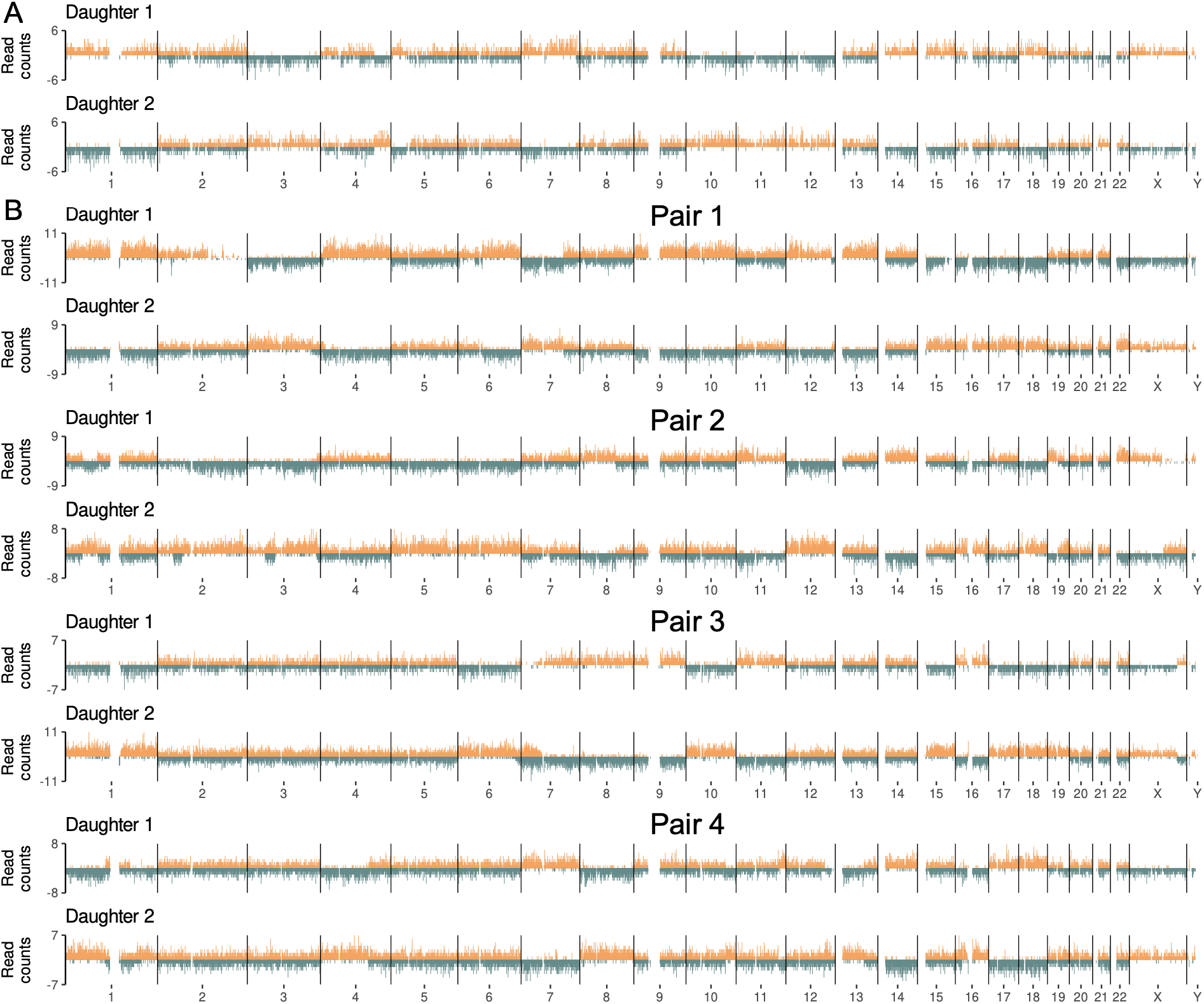
RDCP analysis with sgRNA targeting 237 genomic sites. In sci-L3-Strand-seq plots, reads mapping to the W and C template strands are plotted above and below zero on the y-axis, respectively. SCEs are identified as switches between W and C strand states along a chromosome. **A**. A single RDCP identified from 214 single cells following Cas9 RNP treatment targeting 237 sites with concurrent BrdU labeling. **B**. Four RDCPs exhibiting large-scale SVs recovered from cells labeled with BrdU after Cas9 RNP washout, corresponding to the second division.

## Discussion

This study provides the first genome-wide, single-cell map of SCE induced by CRISPR/Cas9. Three principal findings emerge. First, a single Cas9 DSB at a unique genomic locus is a potent trigger of on-target SCE, with 17-41% of cells undergoing inter-sister strand switching within the same division as cleavage and BrdU labeling. When Cas9 is washed out and cells are labeled in the subsequent cell cycle, the on-target SCE rate drops by 50-74% to a range of 4-17%, confirming that the exchange signal reflects acute repair of the programmed, on-target DSB rather than a heritable alteration of SCE propensity. These strand switches represent non-local interactions between two sister chromatids that are invisible to amplicon sequencing and whole-genome sequencing, yet occur at frequencies comparable to those of the indel outcomes conventionally used to measure editing efficiency. Second, directing Cas9 to 237 repetitive targets produces only mild on-target SCE enrichment in the bulk population, but a subset of high-SCE cells shows more significant enrichment, consistent with a stochastic model in which a fraction of cells sustains a disproportionate share of recombinogenic cleavage events. Even in these high-SCE cells, the number of on-target SCEs per cell remains below 15, far fewer than the 237 available cut sites, indicating that most breaks never formed and/or were resolved by pathways that did not produce SCE. Third, Cas9 nickases (D10A and H840A) neither stall the cell cycle nor produce significant on-target SCE enrichment under the conditions tested, consistent with the expectation that single-strand nicks are less recombinogenic when delivered to a single genomic site without a paired offset nick, although we cannot exclude SCE formation at other time points.

Among the most informative outcomes in this dataset is a subset of Cas9-induced SCE events that display a distinctive structural signature in RDCP analysis: one daughter cell carries a strand switch with a WWC or WCC template-strand pattern at the SCE junction, while the paired daughter cell carries a deletion at the matching SCE junction (**Fig.3**). This “WWC-or- WCC/deletion pair” signature is inconsistent with classical homologous recombination, in which both daughters inherit reciprocal, copy-neutral strand switches. Instead, it resembles the products predicted by the URR resolution model, in which TRAIP-dependent unloading of the CMG helicase at stalled or converging replication forks exposes the template strands to nuclease cleavage, generating one-ended DSBs megabases apart, that are then joined by DNA polymerase theta to produce a sister chromatid with a deletion and an accompanying strand switch (Deng et al., 2019; Sonneville et al., 2019). Independently, van Vugt and colleagues demonstrated that SCEs can form in BRCA1/2-null cell lines, and are enriched at common fragile sites, providing cytogenetic evidence for an HR-independent, URR-coupled SCE pathway (Heijink et al., 2022). Because this pathway does not require end resection or strand invasion, it can operate in BRCA1/2-deficient cancer cells.

**Fig. 3.**
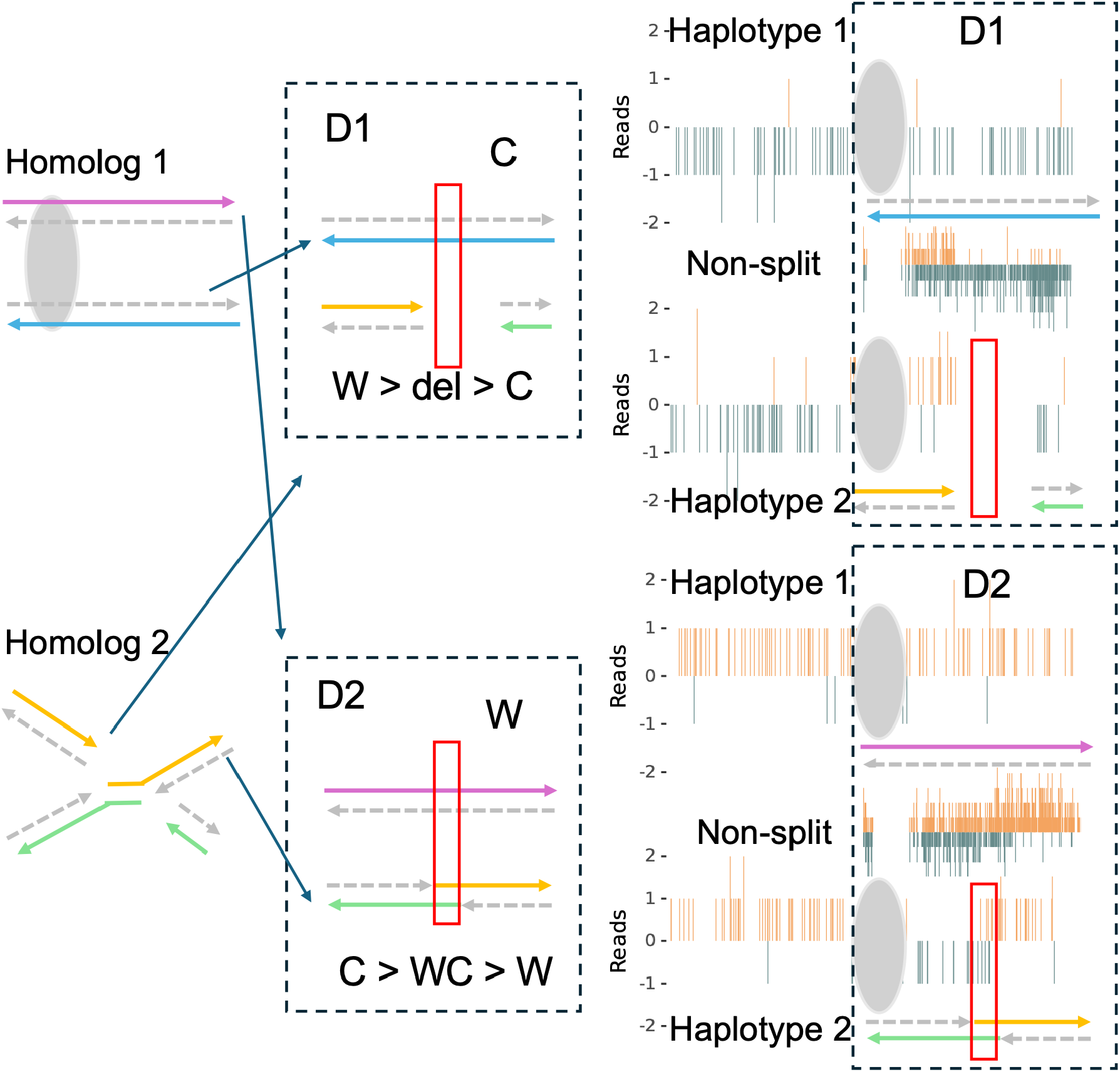
Haplotype-aware analysis of observed RDCP (Pair 4, chr1) shows SV patterns at the SCE junctions consistent with the predicted RDCP signature of SCE mediated by URRs or replication termination zones (green shaded area), although the lagging strands, rather than the leading strands, must be resolved to generate these mitotic breaks. Oval depicts centromere. D1: daughter cell 1. D1 has a deletion region at the SCE junction. The corresponding region in D2 has a WC region within the haplotype with the SCE (WWC if depicted together with the other haplotype) as both strands are segregated without replication. Details in text.

Two recent studies revealed how the CDK1-TTF2-TRAIP axis is cell-cycle regulated to trigger mitotic CMG helicase disassembly and fork cleavage: one study showed a two-fold SCE reduction in mouse ES cells (Fujisawa and Labib, 2026), while the other showed that disrupting the TRAIP-TTF2 interaction reduced common fragile site deletions (Can et al., 2026). Our observation of the “WWC-or-WCC/deletion pair” signature in wild-type cells provides, to our knowledge, the first genetic evidence linking a deletion with SCE and revealing both W and C unreplicated template strands present in the reciprocal daughter cell, consistent with this mechanism at single-cell genomic resolution (illustrated in **Fig.3**), although this is limited by the observation of only one RDCP. Several mechanistic questions follow directly. Does TRAIP-mediated CMG unloading license structure-specific nucleases to process the exposed fork? Which resolvases (MUS81-EME1, SLX1-SLX4, GEN1) execute the cleavage? Note that this specific example actually represents lagging strand cleavage and is possible, although unlikely, due to Cas9 cleavage, RDCP analysis can resolve whether the leading or lagging strands are cleaved, which provides a unique opportunity if coupled with resolvase perturbations. Finally, we observe one such signature among approximately 36 otherwise normal SCEs in wild-type cells; determining the rate of this signature in BRCA1/2-null and other HR-perturbed conditions in spontaneous, rather than Cas9-induced SCE, will be essential for understanding the contribution of URR-type resolution to genome instability in cancer.

## Limitation

A limitation of this study is that all experiments were performed in wild-type cells. In addition, the consequences of Cas9-induced DSBs may depend on both the cell-cycle stage at which cleavage occurs and the number of times a target is cut. Because Cas9 can undergo multiple rounds of cleavage and repair at the same target site, the genomic rearrangements observed here cannot necessarily be attributed to a single cleavage event, limiting mechanistic inference from individual repair outcomes. Approaches that provide tighter temporal control over Cas9 activity and the number of cleavage events will be important for resolving these questions in future studies. The genetic regulation of Cas9-induced SCE by the NHEJ, microhomology-mediated end joining (MMEJ), and HR pathways also remain to be dissected. In our companion paper, we found minimal role of NHEJ in regulating spontaneous SCE. However, it is unknown whether loss of NHEJ would redirect a greater fraction of Cas9 DSBs toward SCE, particularly given that Cas9 DSBs are two-ended, or whether loss of NHEJ abolishes apparent SCEs formed by dNHEJ across sister chromatids. Similarly, the role of MMEJ in competing with or contributing to SCE outcomes at isochromatid breaks and/or at URRs has not been tested. Systematic genetic perturbation in future experiments will be necessary to delineate the pathway hierarchy governing SCE at Cas9-induced isochromatid DSBs.

## Conclusion

In summary, this work establishes that Cas9-induced DSBs trigger SCE at frequencies that are high for single-site cuts and detectable, though modest, for multi-target sites. Beyond quantifying on-target SCE rates, RDCP analysis reveals a “WWC-or-WCC/deletion pair” signature that cannot arise from classical homologous recombination and instead implicates a TRAIP/URR-type resolution pathway operating independently of BRCA1/2. This signature, observed here at single-cell genomic resolution in wild-type cells, provides a molecular framework for the HR-independent SCEs previously inferred from cytogenetic studies in BRCA-deficient backgrounds. More broadly, the isochromatid geometry of Cas9 breaks, in which both sisters are cleaved at the same locus, creates a recombination substrate structurally distinct from the single-ended, replication-born lesions that generate spontaneous SCE (Chovanec et al., 2026a), and this distinction is reflected in a measurable fraction of mutagenic outcomes. These findings underscore that single-cell, strand-level resolution is necessary for a complete accounting of the repair consequences of programmable nucleases.

## Supporting information

Cas9_RNP_scripts.zip

## Acknowledgments

This work was supported by the National Institute of General Medical Sciences (R35GM142511 to Y.Y.), W. M. Keck Foundation (1002444 to Y.Y.) and DOD CDMRP (PC250408 to Y.Y.).

## Author Contributions

Conceptualization: P.C., Y.Y.; Data curation: P.C, S.H., Y.Y.; Software: P.C., Y.Y.; Formal analysis: P.C., Y.Y.; Funding acquisition: Y.Y.; Investigation: P.C., S.H., Y.Y.; Methodology: P.C., S.H., Y.Y.; Supervision: Y.Y.; Writing - original draft: P.C., Y.Y.; Writing - review & editing: P.C., S.H., Y.Y.

## Methods

### Cell culture

BJ-5ta cells (CRL-4001, ATCC) were cultured in DMEM/F12 medium supplemented with 10% fetal bovine serum (FBS) and 1× penicillin-streptomycin at 37 °C in 5% CO_2_.

### Cas9 RNP delivery

Cas9 ribonucleoprotein (RNP) complexes were assembled using Alt-R *S. pyogenes* Cas9 Nuclease V3 (1081058; IDT) or corresponding nickase variants (Alt-R Cas9 D10A Nickase V3; Alt-R Cas9 H840A Nickase V3) together with Alt-R tracrRNA (1072532; IDT) and Alt-R crRNA.

For single-targeting site experiments, crRNAs targeted coding exons of four DNA repair genes (LIG4: 5′-GCTTATACGGATGATCATAA-3′; XRCC4: 5′-ATGGTCATTCAGCATGGACT-3′; XRCC1: 5′-TGCAGGACACGACATGG-3′; LIG3: 5′-CTGTTAGGTACACATCACCG-3′). For the 237-cut condition, a crRNA targeting a repetitive element with 237 predicted genomic sites was used (5′-TCAGCACTTTGGGAGACCA-3′). CrRNA and tracrRNA were diluted to 1 μM in duplex buffer (11010301; IDT), annealed at 95 °C for 5 min, and cooled to room temperature for 30 min to form gRNA. Cas9 (or nickase) was diluted to 1 μM in OptiMEM, combined with gRNA and Cas9 PLUS reagent (CRISPRMAX kit, CMAX00003; Invitrogen), and incubated at room temperature for 5 min to assemble RNPs. Reverse transfection was performed with cells at 2×10^5^ cells/mL.

The 237-site sgRNA targets a repetitive Alu sequence and was selected from a screen of >20,000 sgRNAs designed against repetitive sequences occurring at >200 genomic sites. The guide was selected for an intermediate cellular phenotype that permitted simultaneous targeting of a large number of genomic sites without excessive loss of cells; characterization of the broader repetitive-element screen is outside the scope of this study.

### BrdU labeling conditions

Two labeling schemes were used: in the simultaneous condition (“24 hrs”), BrdU (40 μM final concentration) was added at the time of Cas9 RNP delivery, and cells were cultured for 24 hours prior to fixation, such that Cas9 cleavage and BrdU incorporation occurred within the same cell division. In the washout condition (“24 ws”), Cas9 RNP was delivered without BrdU for 24 hours, after which cells were washed and replated in fresh medium containing 40 μM BrdU for an additional 40 hours before fixation, labeling the division following Cas9 activity.

### Sci-L3-Strand-seq library generation

Sci-L3-Strand-seq was performed as previously described (Chovanec and Yin, 2025; Yin et al., 2019). Briefly, cells were trypsinized and fixed in 1.5% formaldehyde in PBS at 1×10^6^ cells/mL for 10 min at room temperature, followed by quenching with glycine. Cells were snap-frozen in nuclei freezing buffer (NFB: 50 mM Tris pH 8.0, 25% glycerol, 5 mM Mg(OAc)_2_, 0.1 mM EDTA, 5 mM DTT, 1× protease inhibitor).

Tagmentation was performed using Tn5 transposase with 24 first-round barcodes as described (Yin et al., 2019). After tagmentation (55 °C, 15 min), reactions were quenched with lysis buffer (LBT) and nuclei were pooled. Ligation with 72 second-round barcodes was followed by staining with Hoechst-33258 (10 ng/μL) and FACS sorting (200-300 nuclei per well into 96-well plates). Plates were stored at -80 °C.

Following gap extension, UV treatment (270 mJ/cm^2^, 365 nm), USER digestion, and UGI treatment, T7 in vitro transcription was performed (14-16 hours, 37 °C). RNA purification, reverse transcription, second-strand synthesis (third barcode incorporation), and Illumina library preparation were carried out using standard kits (NEB).

### SCE analysis

Raw FASTQ files were processed using the sciL3Pipe Snakemake pipeline (v0.2.1), including barcode demultiplexing, alignment, and single-cell BAM generation. Breakpoint identification and filtering were performed using breakpointR (v1.10.0) and sciStrandR (v0.2.3). Cells passing quality filters (background estimate > 0 and ≤ 0.08; strand-neutral < 0.75; median reads per Mb > 2) were retained.

### On-target SCE identification

SCEs were classified as on-target by two criteria. First, a breakpointR-called breakpoint within +/-3 Mb of the cut site was counted as a direct call. Second, a rescue procedure examined strand-state in two 10 Mb bins flanking the cut site (each offset 3 Mb from the gene boundary) and called a rescued SCE if the flanking bins showed a strand-state transition (e.g., WW-to-CC, WW-to-WC, CC-to-WC etc.). On-target strand switches were defined as the union of called and rescued events.

### On-target SCE enrichment analysis (237-cut condition)

To test enrichment, observed SCEs overlapping cut-site windows were compared to those overlapping size-matched random genomic intervals using a one-sided’s exact test. Analyses were performed across all cells and repeated for high-SCE cells (cells with >=8 or >=9 post-filter SCEs).

### Reciprocal daughter cell identification

The genome was partitioned into 1 Mb bins across autosomes and chromosome X. For each cell, W and C reads were counted per bin, and a strand ratio score was calculated as (W - C)/(W + C), yielding +1 (WW) or -1 (CC). Bins with fewer than four reads were set to 0. For each cell pair, a “sisterness” score was computed by averaging |score_i_ + score_j_| across informative bins (|score| > 0.5 in at least one cell). Scores approach 0 for reciprocal strand inheritance and 2 for identical states. The top 20 candidate pairs with the lowest scores were selected and validated by visual inspection of Strand-seq ideograms.

---

**Figure S1.**
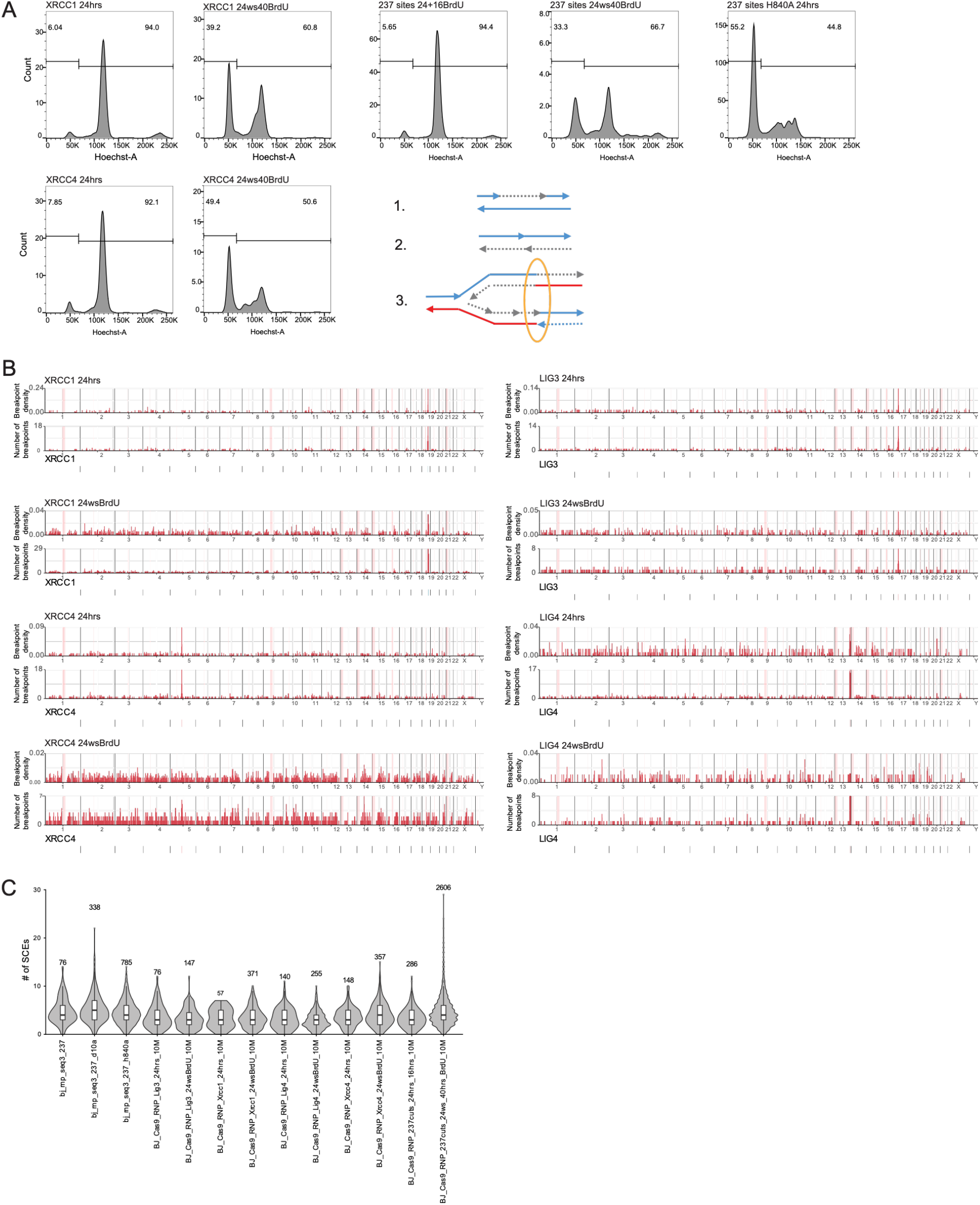
Related to **Fig.1**. **A**. Additional FACS plots showing Cas9-induced cell cycle arrest. Local and non-local repair examples. Local fill-in synthesis (1) & NHEJ (2) preserve continuity of the chromatid; SCE requires cis-chromatid disengagement (3) and are coupled with a template strand directionality switch from blue to red or red to blue. **B**. Additional genome-wide pileup showing on-target SCE. For each sgRNA, top: breakpointR-based SCE calls; boRom: breakpointR-based and rescued SCE calls. **C**. Quantification of spontaneous SCE as a result of polyclonal/bulk Cas9 RNP knockouts (see Tab.S2 for pair-wise statistical testing).

**Figure S2.**
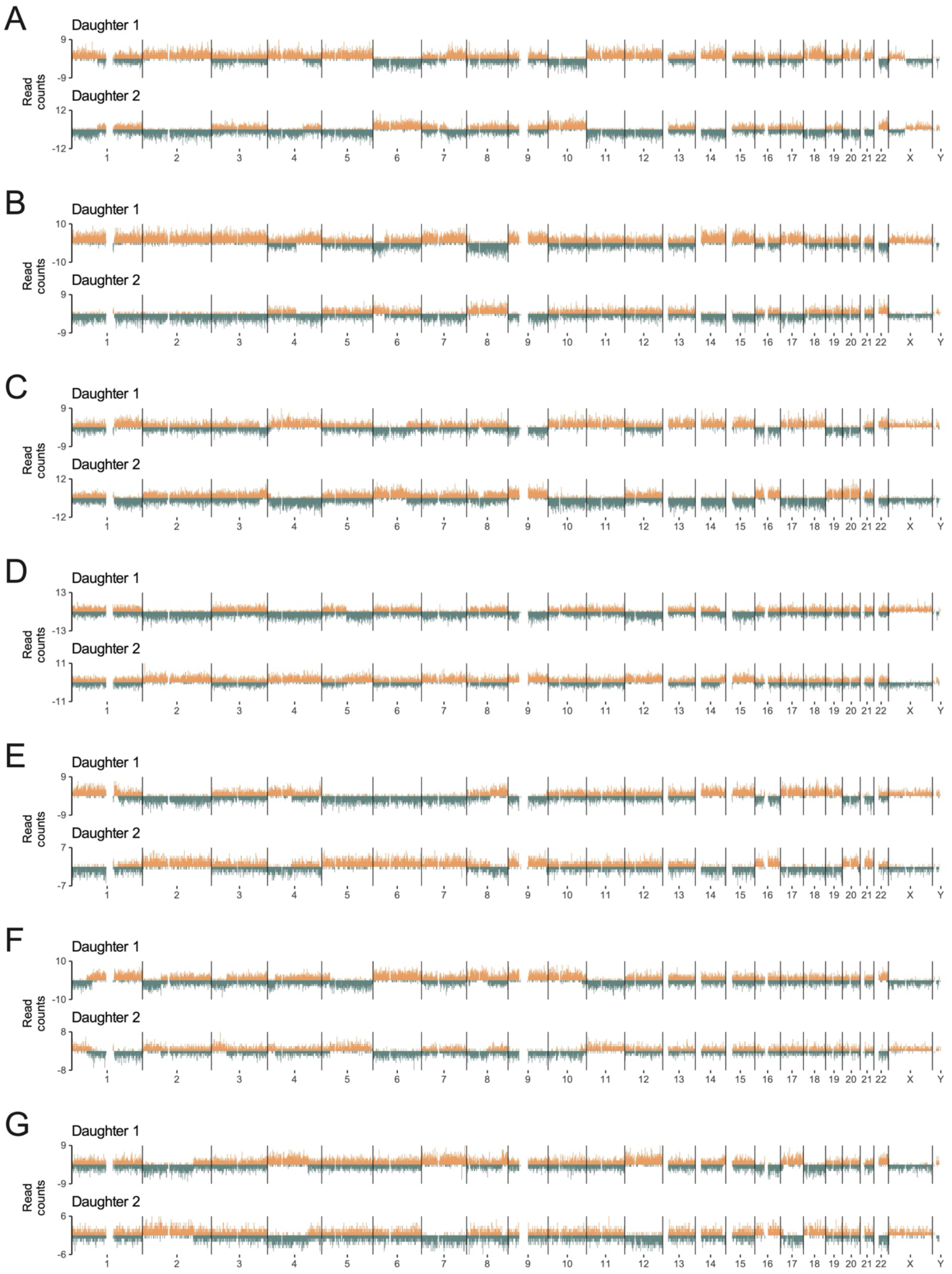
Additional RDCP plots with sgRNA targeting 237 genomic sites. Seven RDCPs without large-scale SVs recovered from cells labeled with BrdU a\er Cas9 RNP washout, corresponding to the second division.

**Figure S3.**
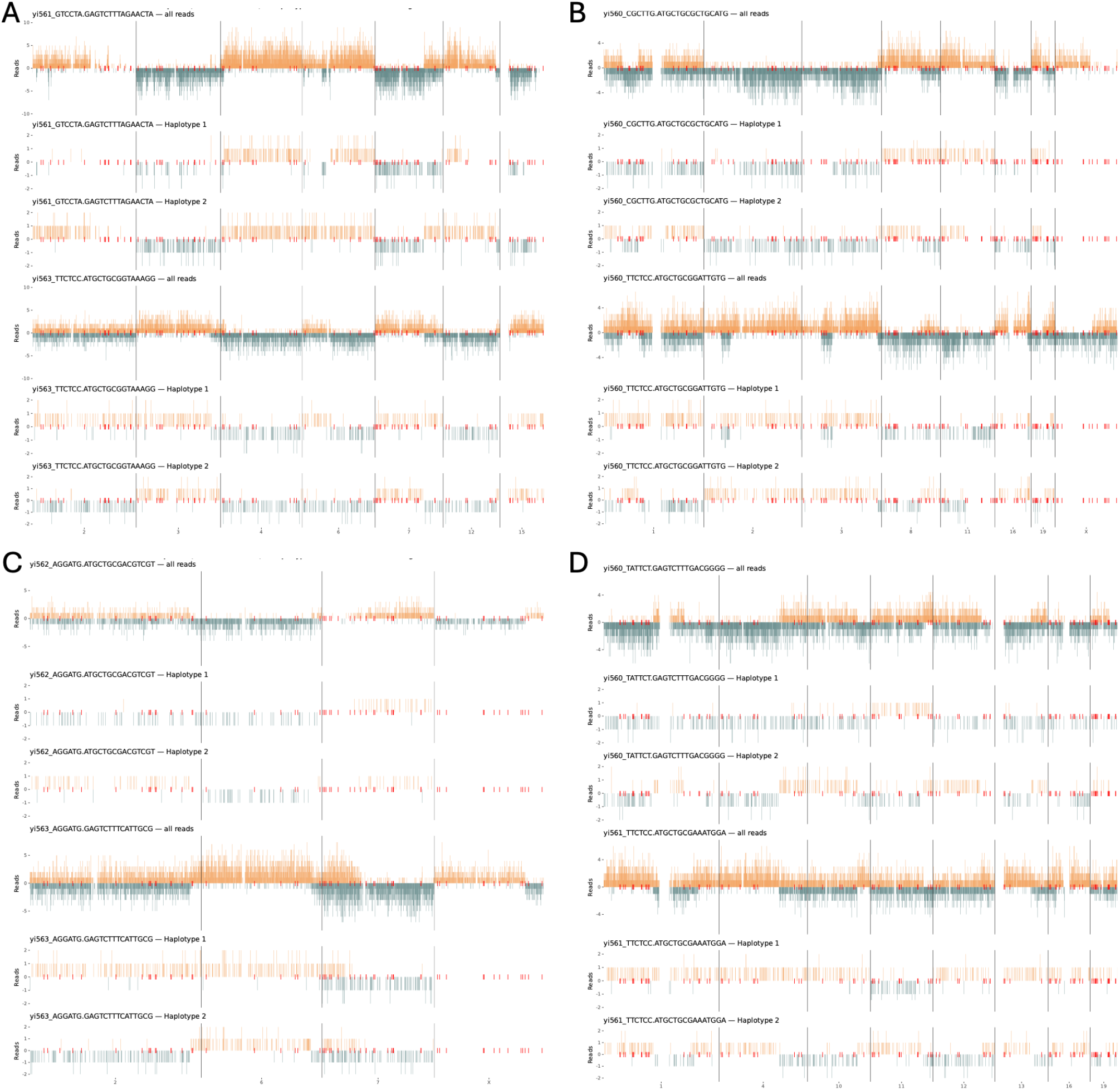
Haplotype-aware analysis of RDCP with large-scale SVs. **A**. Pair1 in Fig.2B. **B**. Pair 2 in Fig.2B. **C**. Pair3 in Fig.2B. **D**. Pair4 in Fig.2B, the mechanistic interpretation is illustrated in Fig.3.

