## Supplementary material for "Genome-wide mapping of Cas9-induced sister chromatid exchange across single and over 200 genomic targets": Cas9_RNP_scripts.zip: 00_setup_functions.R

```
# 00_setup_functions.R
# Defines all helper functions and constants, saves to data/setup_env.RData
# Run with: /u/project/yeastyin/yjiang/miniconda3/envs/sci_bpR_claude/bin/Rscript scripts/00_setup_functions.R

library(sciStrandR)
library(dplyr)
library(tidyr)
library(purrr)
library(readr)
library(stringr)
library(tibble)
library(ggplot2)
library(patchwork)
library(foreach)
library(parallel)
library(regioneR)
library(ggbeeswarm)
library(ggpointdensity)
library(viridis)
library(ggrastr)
library(plyranges)

fig_dir <- "data/figures_10conditions/"
tab_dir <- "data/tables_10conditions/"
dir.create(fig_dir, showWarnings = FALSE, recursive = TRUE)
dir.create(tab_dir, showWarnings = FALSE, recursive = TRUE)

# ---- Theme ----
legend_theme <- theme(legend.position='top',
        legend.justification='right',
        legend.direction='horizontal',
        legend.title = element_blank(),
        legend.text = element_text(size=8),
        legend.key.height= unit(0.4, 'cm'),
        legend.key.width= unit(0.4, 'cm'),
        legend.margin=margin(c(0,0,0,0)),
        legend.spacing.y = unit(0, "mm"),
        legend.spacing.x = unit(0.3, "mm"),
        legend.key=element_blank(),
        plot.title = element_text(size = 11))

# ---- Coordinates (hg19, Ensembl naming) ----
xrcc1_coord <- "19:44047463-44079679"
lig3_coord  <- "17:33307531-33336762"
lig4_coord  <- "13:108859790-108870697"
xrcc4_coord <- "5:82367495-82643756"

# ---- Reference paths ----
deeptools_plotHeatmap_path <- "/u/project/yeastyin/chovanec/miniconda3/envs/deeptools/bin/plotHeatmap"
blacklist_path <- "/u/project/yeastyin/chovanec/genomes/hg19-blacklist.v2.ensembl.bed"
bj_inv_path <- "/u/project/yeastyin/chovanec/software/scistrandfig/inst/extdata/regions/BJ-5ta_inv_bl_name.bed"
guide_bam_path <- "/u/project/yeastyin/chovanec/projects/RNP_workup/YY_SEQ3_237_GUIDE1.srt.bam"

# ---- Functions ----

plot_wrapper <- function(breaks, hotspots, coord, assembly="hg19", ensembl=T,
                         fill_by_col = NULL, custom_colors = NULL, fill_col="grey35", highlights=NULL){
  p1 <- plot_ranges(breaks, assembly, ensembl = ensembl, chr_size_limit = 1000000, subset_coordinates = coord, by_mcols = "sample", min_width = 200000, fill_by_mcol=fill_by_col, custom_colors=custom_colors, fill_col=fill_col, highlights = highlights) +
          theme(axis.ticks.x = element_blank(), axis.text.x = element_blank(), panel.grid.minor.y = element_line()) +
          labs(subtitle = "Breaks")
  p2 <- plot_ranges(hotspots, assembly, ensembl = ensembl, chr_size_limit = 1000000, subset_coordinates = coord, min_width = 200000) +
    theme(axis.ticks.x = element_blank(), axis.text.x = element_blank(), axis.text.y = element_blank(), axis.ticks.y = element_blank()) +
    labs(subtitle = "Hotspots")
  return(list(p1,p2))
}

bins_of_interest <- function(gr, dist_from_gr=3000000, bin_width=10000000){
  L_bin <- plyranges::flank_left(gr, width = bin_width)
  L_bin <- plyranges::shift_left(L_bin, shift = dist_from_gr)
  R_bin <- plyranges::flank_right(gr, width = bin_width)
  R_bin <- plyranges::shift_right(R_bin, shift = dist_from_gr)
  return(c(L_bin,R_bin))
}

state_assignment_bins <- function(bins, frags){
  ranges_tiles_ul <- bins
  strand(ranges_tiles_ul) <- '+'
  ranges_tiles_ul$Ws_unq <- GenomicRanges::countOverlaps(ranges_tiles_ul, frags)
  strand(ranges_tiles_ul) <- '-'
  ranges_tiles_ul$Cs_unq <- GenomicRanges::countOverlaps(ranges_tiles_ul, frags)
  strand(ranges_tiles_ul) <- '*'
  tile_wc_ratio <- (ranges_tiles_ul$Ws_unq-ranges_tiles_ul$Cs_unq)/(ranges_tiles_ul$Ws_unq+ranges_tiles_ul$Cs_unq)
  ranges_tiles_ul$tile_wc_ratio <- tile_wc_ratio
  ranges_tiles_ul$states <- "?"
  ranges_tiles_ul$states[tile_wc_ratio > 0.8] <- "ww"
  ranges_tiles_ul$states[tile_wc_ratio < -0.8] <- "cc"
  ranges_tiles_ul$states[tile_wc_ratio < 0.2 & tile_wc_ratio > -0.2] <- "wc"
  ranges_tiles_ul$states <- factor(ranges_tiles_ul$states, levels = c("wc", "ww", "cc", "?"))
  return(ranges_tiles_ul)
}

run_check_location_for_breakpoints <- function(data_folder,
                                               coord, cent=NULL, samples=NULL,
                                               type=c("diploid", "haploid"),
                                               assembly="hg19", ensembl=TRUE,
                                               num_threads=1, filtered=TRUE,
                                               bin_width = 5e+06, dist_from_gr = 1e+06, limit=NULL){
  if(length(type) > 1) stop("Specify one of the possible options for type.")
  if(is(coord, "character")){
    coord_gr <- coordinates2gr(coord, genome_assembly = assembly)
  }else if(is(coord, "GRanges")){
    coord_gr <- coord
  }else{
    stop("coord not in correct format.")
  }
  data_files <- get_files_to_load(data_folder, samples, filtered=filtered, limit=limit)
  if(is.null(cent)){
    cent <- get_centromere_coordinates(assembly, FALSE, ensembl = ensembl)
  }
  if(num_threads > 1){
    pb <- txtProgressBar(max = length(data_files), style = 3)
    progress <- function(n) setTxtProgressBar(pb, n)
    opts <- list(progress = progress)
    message("\nUsing ",num_threads," threads")
    cl <- parallel::makeCluster(num_threads)
    doSNOW::registerDoSNOW(cl)
    out_df <- foreach(i=1:length(data_files), .packages = c("sciStrandR", "dplyr", "purrr", "plyranges"),
                      .export = c("check_location_for_breakpoints", "bins_of_interest", "state_assignment_bins"),
                      .options.snow = opts, .combine=rbind) %dopar% {
          check_location_for_breakpoints(data_files[i], coord_gr, cent=cent,
                                         type=type, ensembl=ensembl, assembly=assembly,
                                         bin_width=bin_width, dist_from_gr=dist_from_gr)
      }
    parallel::stopCluster(cl)
  }else{
    out_lst <- list()
    for(data_path in data_files){
        out_lst[[data_path]] <- check_location_for_breakpoints(data_path, coord_gr, cent=cent,
                                                              type=type, ensembl=ensembl, assembly=assembly,
                                                              bin_width=bin_width, dist_from_gr=dist_from_gr)
    }
    out_df <- bind_rows(out_lst)
  }
  rownames(out_df) <- NULL
  return(out_df)
}

check_location_for_breakpoints <- function(path, coord, cent,
                                           type=c("diploid", "haploid"), ensembl=TRUE, assembly="hg19",
                                           bin_width=5e+06, dist_from_gr=1e+06){
  # Handle both .rds and .RData formats
  if (grepl("\\.RData$", path, ignore.case = TRUE)) {
    env <- new.env()
    load(path, envir = env)
    f <- ls(env)[1]
    dehcr_filt <- get(f, envir = env)
  } else {
    f <- load_from_rds(path)
    dehcr_filt <- get(f)
  }
  fragments <- dehcr_filt$fragments
  breaks <- dehcr_filt$breaks
  if(is.null(bin_width)){
    if(is.null(cent)) cent <- get_centromere_coordinates(assembly, FALSE, ensembl = ensembl)
    arms <- gaps(cent)
    arms <- arms[strand(arms) == "*"]
    coord_arms <- arms %>% plyranges::filter_by_overlaps(coord) %>% plyranges::setdiff_ranges(coord)
  } else{
    coord_grl <- split(coord)
    coord_arms <- map(coord_grl, \(x) trim(bins_of_interest(x, dist_from_gr = dist_from_gr, bin_width = bin_width)))
  }
  coord_arms_gr <- unlist(GRangesList(coord_arms))
  message("Breaks within interval: ", length(plyranges::filter_by_overlaps(breaks, coord_arms_gr)))
  df_out <- list()
  for(i in seq_along(coord_arms)){
    L_R_states <- suppressWarnings(state_assignment_bins(coord_arms[[i]], fragments))
    L_R_states <- sort(L_R_states)
    new_states <- paste(L_R_states$states, collapse = "_")
    if(type == "haploid"){
      rescue_break <- new_states %in% c("ww_cc", "cc_ww")
    }else if(type == "diploid"){
      rescue_break <- new_states %in% c("ww_cc", "cc_ww", "ww_wc", "wc_ww", "cc_wc", "wc_cc")
    }
    df_out[[i]] <- data.frame(ID = dehcr_filt$ID,
             breakpoint_called = length(plyranges::filter_by_overlaps(breaks, coord_arms[[i]])),
             tiled_states = new_states,
             rescue_break = rescue_break,
             coord_idx = i,
             background.estimate=dehcr_filt$lib.metrics['background.estimate'],
             med.reads.per.MB=dehcr_filt$lib.metrics['med.reads.per.MB'],
             perc.coverage=dehcr_filt$lib.metrics['perc.coverage'])
  }
  out <- bind_rows(df_out)
  assertthat::assert_that(nrow(out) == length(coord), msg = "Not all coordinates are being returned.")
  return(out)
}

read_summary <- function(br_path, num_threads=1){
  out <- breakpoint_summary(br_path, num_threads=num_threads) %>%
    mutate(sample = sub("^([^\\.]*\\.[^\\.]*).*", "\\1", .id)) %>%
    mutate(Tn5=substring(sapply(strsplit(.id, "\\."), "[[", 2),1,8)) %>%
    mutate(ligation=substring(sapply(strsplit(.id, "\\."), "[[", 2),9,15)) %>%
    mutate(sss=sapply(strsplit(.id, "\\.|_"), "[[", 2)) %>%
    mutate(library=sapply(strsplit(.id, "\\.|_"), "[[", 1)) %>%
    mutate(library_sss=sapply(strsplit(.id, "\\."), "[[", 1))

  tn5_barcodes <- data.frame(stringsAsFactors = FALSE,
    Tn5 = c("TGATATTG","GATCCCGT","CTCGATTA","CATCAAGG","TCCTTGTG","GGTCATAT","ATCGCGTT",
            "CATGCCCC","GTTACGCG","CCGCGCTT","TCTTAGTG","TCGGCCTA",
            "CTTTCTCT","TCGCGTTT","GTCAGTAG","CCATGGAA",
            "ATGCTGCG","GAGTCTTT","TACGATAT","ACCATTTA","ATCGGGAC",
            "GACGTCGG","CATTGTGT","TTTGACTC")) %>%
    mutate(barcode_num=seq(1,24,1))

  sss_barcodes <- data.frame(
    sss_num = c("c1","c2","c3","c4","c5","c6","c7","c8",
                "l1","l2","l3","l4","l5","l6","l7","l8",
                "l9","l10","l11","l12","l13","l14","l15","l16","l17","l18"),
    sss = c("CACGTG","GTGCAG","ACTTCG","TGAACG","CCAGTG","TTCGAG","GAACCG","AGGACG",
            "ACGCGA","CGCTTG","GTCCTA","AGGATG","TTCTCC","ACCACT","TTTCGC","CGGTGG",
            "TATTCT","ACTTAA","TAAAGA","GAGTTT","GGGTGC","GGGCCG","AATTGA",
            "TAAGCG","TAATGC","GTCTAT"))

  out <- out %>% left_join(tn5_barcodes) %>% left_join(sss_barcodes)
  return(out)
}

plot_background <- function(df, limits=c(0,0.5), threshold=0.08){
  df %>%
  mutate(filter_col = ifelse(background.estimate > 0 & background.estimate <= threshold, "F", "T")) %>%
   ggplot(aes(x = library, y = background.estimate)) +
    geom_quasirandom(aes(color = filter_col), size = 0.1, dodge.width=NULL) +
    geom_boxplot(width = 0.2, outlier.size = 0.1, outlier.colour = NA, fill = NA, color="black") +
    xlab(NULL) + ylab("Background estimate") +
    theme_classic() +
    geom_hline(yintercept = threshold, alpha=0.5) +
    scale_color_manual(values = c("F" = "darkgrey", "T" = "tomato")) +
    scale_y_continuous(expand = expansion(mult = c(0.02, 0)), limits = limits, breaks = seq(limits[1],limits[2],0.1)) +
    theme(strip.background=element_blank(), legend.position='none',
          axis.text.x = element_text(angle=45, hjust = 1))
}

plot_strand_neutral <- function(df, threshold=0.08){
  df %>%
  ggplot(aes(x=wc.perc+uknw.perc, y=background.estimate)) +
    annotate("rect", xmin = -Inf, xmax = 15, ymin = -Inf, ymax = threshold, fill= "red", alpha=0.1) +
    stat_pointdensity(geom = ggrastr:::GeomPointRast, raster.dpi=300, dev = "cairo", scale = 0.5, adjust = 0.5) +
    scale_color_viridis(name="N neighbors", guide = guide_colorbar(frame.colour = "black", ticks.colour = "black", frame.linewidth = 1, ticks.linewidth = 1)) +
    facet_grid(~library) +
    geom_vline(xintercept = 15) + geom_vline(xintercept = 75) +
    geom_hline(yintercept = threshold) +
    ylab("Background estimate") + xlab("Strand-neutral percentage") +
    theme_bw()+
    theme(legend.position='right', legend.key.width=unit(0.4, 'cm'),
          strip.background=element_blank(), axis.line = element_line(colour = "black"),
          panel.grid.major = element_blank(), panel.grid.minor = element_blank(),
          panel.border = element_blank(), panel.spacing = unit(1, "lines"),
          panel.background = element_blank()) +
    scale_y_continuous(expand = c(0,0)) +
    scale_x_continuous(expand = c(0,0), limits = c(0, 100.01))
}

quality_filter <- function(df, background_estimate=0.08, strand_neutral = 75, mrpm=2){
  df %>% filter(background.estimate <= background_estimate &
                background.estimate > 0 &
                wc.perc+uknw.perc < strand_neutral &
                med.reads.per.MB > mrpm)
}

add_rescue_breaks <- function(filt_gr, rescue_df, rescue_coord_str, stretch_amount=6000000){
  rescue_coord <- coordinates2gr(rescue_coord_str, genome_assembly = "hg19") %>% plyranges::stretch(stretch_amount)
  rescue_samples <- rescue_df %>% filter(rescue_break == TRUE) %>%
    mutate(sample = sub("^([^\\.]*\\.[^\\.]*).*", "\\1", ID)) %>% pull(sample)
  if(length(rescue_samples) == 0) return(filt_gr)
  ignore_rescues <- filt_gr[filt_gr$sample %in% rescue_samples] %>%
    plyranges::filter_by_overlaps(rescue_coord) %>% as.data.frame() %>% pull(sample)
  rescue <- setdiff(rescue_samples, ignore_rescues)
  if(length(rescue) == 0) return(filt_gr)
  rescue_coord <- rep(rescue_coord, length(rescue))
  mcols(rescue_coord) <- DataFrame(genoT=rep("?-?", length(rescue)),
                                    deltaW=rep(0, length(rescue)),
                                    file=rescue, sample=rescue)
  return(c(filt_gr, rescue_coord))
}

run_single_cut_pileup <- function(data_folders, samples, num_threads=10){
  grl_all <- breakpoints_to_list(data_folders, samples=samples, num_threads=num_threads, filtered=FALSE)
  grl_filt <- breakpoints_to_list(data_folders, samples=samples, num_threads=num_threads, filtered=TRUE)
  gr_all <- unlist(grl_all)
  names(gr_all) <- NULL
  mcols(gr_all)$sample <- gsub("\\.(human|hg19)\\.filt\\.bam\\.RData_filtered", "", gr_all$file)
  grl_all <- split(gr_all, gr_all$sample)
  gr_filt <- unlist(grl_filt)
  names(gr_filt) <- NULL
  mcols(gr_filt)$sample <- gsub("\\.(human|hg19)\\.filt\\.bam\\.RData_filtered", "", gr_filt$file)
  list(grl_all=grl_all, gr_filt=gr_filt)
}

build_contingency_table <- function(break_grl, roi_gr, genome, seed=3,
                                    breakpoint_folder=NULL, bin_width = 10000000, dist_from_gr = 3000000,
                                    type="diploid", num_threads=1, filtered=TRUE){
  set.seed(seed)
  random_regions_237 <- randomizeRegions(A = roi_gr, genome = genome, allow.overlaps=FALSE, per.chromosome = TRUE)
  if(!is.null(breakpoint_folder)){
    sample_ids <- sub("^([^\\.]*\\.[^\\.]*).*", "\\1", unique(names(break_grl)))
    sample_breaks_rescue <- run_check_location_for_breakpoints(breakpoint_folder,
                                 coord = roi_gr, samples = sample_ids,
                                 num_threads = num_threads, bin_width = bin_width, dist_from_gr = dist_from_gr, type=type, filtered=filtered)
    random_breaks_rescue <- run_check_location_for_breakpoints(breakpoint_folder,
                                 coord = random_regions_237, samples = sample_ids,
                                 num_threads = num_threads, bin_width = bin_width, dist_from_gr = dist_from_gr, type=type, filtered=filtered)
    sample_breaks_rescue_lst <- split(sample_breaks_rescue, sample_breaks_rescue$ID)
    random_breaks_rescue_lst <- split(random_breaks_rescue, random_breaks_rescue$ID)
    sample_counts <- list()
    for(s in seq_along(sample_breaks_rescue_lst)){
      total_regions <- length(roi_gr)
      roi_overlap <- sum(sample_breaks_rescue_lst[[s]]$rescue_break == TRUE)
      roi_nonoverlap <- total_regions - roi_overlap
      rand_overlap <- sum(random_breaks_rescue_lst[[s]]$rescue_break == TRUE)
      rand_nonoverlap <- total_regions - rand_overlap
      sample_counts[[s]] <- data.frame(sample=sample_ids[[s]],
                                  roi_overlap=roi_overlap, roi_nonoverlap=roi_nonoverlap,
                                  rand_overlap=rand_overlap, rand_nonoverlap=rand_nonoverlap,
                                  total_regions=total_regions)
    }
  }else{
    sample_counts <- list()
    for(s in seq_along(break_grl)){
      sample_breaks <- break_grl[[s]]
      sample_id <- sub("^([^\\.]*\\.[^\\.]*).*", "\\1", unique(sample_breaks$sample))
      total_regions <- length(roi_gr)
      roi_overlap <- sum(countOverlaps(sample_breaks, roi_gr))
      roi_nonoverlap <- total_regions - roi_overlap
      rand_overlap <- sum(countOverlaps(sample_breaks, random_regions_237))
      rand_nonoverlap <- total_regions - rand_overlap
      sample_counts[[s]] <- data.frame(sample=sample_id,
                                  roi_overlap=roi_overlap, roi_nonoverlap=roi_nonoverlap,
                                  rand_overlap=rand_overlap, rand_nonoverlap=rand_nonoverlap,
                                  total_regions=total_regions)
    }
  }
  sample_counts_df <- bind_rows(sample_counts)
  contingency <- matrix(
    c(sum(sample_counts_df$roi_overlap), sum(sample_counts_df$roi_nonoverlap),
      sum(sample_counts_df$rand_overlap), sum(sample_counts_df$rand_nonoverlap)),
    nrow = 2, byrow = TRUE,
    dimnames = list(Region = c("Cut_sites", "Random_sites"), Breakpoint = c("Present", "Absent")))
  return(list(sample_counts_df=sample_counts_df, contigency=contingency))
}

# ---- New data base path ----
new_base_path <- "/u/project/yeastyin/yeastyin/projects/Cas9_RNP/data/bpR_output"
new_conditions <- c("BJ_Cas9_RNP_237cuts_24hrs_16hrs",
                     "BJ_Cas9_RNP_237cuts_24ws_40hrs_BrdU",
                     "BJ_Cas9_RNP_Xrcc1_24hrs",
                     "BJ_Cas9_RNP_Xrcc1_24wsBrdU",
                     "BJ_Cas9_RNP_Lig3_24hrs",
                     "BJ_Cas9_RNP_Lig3_24wsBrdU",
                     "BJ_Cas9_RNP_Lig4_24hrs",
                     "BJ_Cas9_RNP_Lig4_24wsBrdU",
                     "BJ_Cas9_RNP_Xrcc4_24hrs",
                     "BJ_Cas9_RNP_Xrcc4_24wsBrdU")

hg19_cent <- get_centromere_coordinates("hg19", ensembl = TRUE)

message("Setup complete. Saving environment...")
save.image("data/setup_env.RData")
message("Done: data/setup_env.RData")
```
