## Supplementary material for "Genome-wide mapping of Cas9-induced sister chromatid exchange across single and over 200 genomic targets": Cas9_RNP_scripts.zip: 01_load_earlier.R

```
#!/usr/bin/env Rscript
# 01_load_earlier.R
# Load and QC-filter all earlier datasets (threshold 0.08)
# Produces: data/summaries_earlier.RData

message("=== 01_load_earlier.R ===")
source("scripts/load_env.R")

# ---- p20250808: bj_rnp_xrcc1_lig3_seq3_237 (yi464/466/468) ----
message("Loading p20250808...")
summary_p20250808 <- bind_rows(
  read_summary("/u/project/yeastyin/chovanec/projects/novaseq250815/bj_rnp_xrcc1_lig3_seq3_237/human/breakpointR_diploid/data_filtered/", num_threads = 10),
  read_summary("/u/project/yeastyin/chovanec/projects/novaseq250919/bj_rnp_xrcc1_lig3_seq3_237/human/breakpointR_diploid/data_filtered/", num_threads = 10)
)

summary_p20250808_filt <- summary_p20250808 %>%
  filter(background.estimate <= 0.08 &
         background.estimate > 0 &
         wc.perc + uknw.perc < 75 &
         med.reads.per.MB > 2)

message("p20250808: ", nrow(summary_p20250808), " total -> ", nrow(summary_p20250808_filt), " after QC")

# ---- p20250409: BJ_rnp_xrcc1 (yi441) and BJ_rnp_lig3 (yi442/443) ----
message("Loading p20250409...")
summary_p20250409 <- bind_rows(
  read_summary("/u/project/yeastyin/chovanec/projects/nextseq250611/BJ5ta_RNP_XRCC1_LIG3/hg19/breakpointR_diploid/data_filtered/", num_threads = 10),
  read_summary("/u/project/yeastyin/chovanec/projects/novaseq250806/BJ5ta_RNP_XRCC1_LIG3/hg19/breakpointR_diploid/data_filtered/", num_threads = 10)
)

# Load pre-computed summary with condition labels
novaseq250806_summary_bj_rnp <- readRDS("/u/project/yeastyin/chovanec/projects/RNP_workup/data/novaseq250806_summary_bj_rnp.rds")

novaseq250806_summary_bj_rnp_filt <- novaseq250806_summary_bj_rnp %>%
  filter(background.estimate <= 0.08 &
         background.estimate > 0 &
         wc.perc + uknw.perc < 75 &
         med.reads.per.MB > 2)

message("p20250409 (novaseq250806): ", nrow(novaseq250806_summary_bj_rnp), " total -> ", nrow(novaseq250806_summary_bj_rnp_filt), " after QC")
message("  BJ_rnp_xrcc1: ", sum(novaseq250806_summary_bj_rnp_filt$condition == "BJ_rnp_xrcc1"))
message("  BJ_rnp_lig3: ", sum(novaseq250806_summary_bj_rnp_filt$condition == "BJ_rnp_lig3"))

# ---- p20250924: BJ RNP (yi478/479) ----
message("Loading p20250924...")
summary_p20250924 <- read_summary("/u/project/yeastyin/chovanec/projects/novaseq251007/BJ_RNP/human/breakpointR_diploid/data_filtered/", num_threads = 10)

# Assign conditions by Tn5 barcode
summary_p20250924 <- summary_p20250924 %>%
  mutate(condition = case_when(
    barcode_num %in% c(1, 2) ~ "bj_rnp_xrcc1",
    barcode_num %in% c(3, 4, 5) ~ "bj_rnp_lig3",
    barcode_num %in% c(6, 7) ~ "bj_rnp_seq3_237",
    barcode_num %in% c(9, 10) ~ "bj_rnp_seq3_237_d10a",
    barcode_num %in% c(11, 12) ~ "bj_rnp_seq3_237_h840a",
    TRUE ~ "miss"
  ))

summary_p20250924_filt <- summary_p20250924 %>%
  filter(background.estimate <= 0.08 &
         background.estimate > 0 &
         wc.perc + uknw.perc < 75 &
         med.reads.per.MB > 2)

message("p20250924: ", nrow(summary_p20250924), " total -> ", nrow(summary_p20250924_filt), " after QC")
for (cond in unique(summary_p20250924_filt$condition)) {
  message("  ", cond, ": ", sum(summary_p20250924_filt$condition == cond))
}

# ---- HAP1 yi433 ----
message("Loading HAP1...")
novaseq250806_summary_hap1_rnp <- readRDS("/u/project/yeastyin/chovanec/projects/RNP_workup/data/novaseq250806_summary_hap1_rnp.rds")

# Haploid: wc+uknw < 15%
summary_hap1_haploid_filt <- novaseq250806_summary_hap1_rnp %>%
  filter(background.estimate <= 0.08 &
         background.estimate > 0 &
         wc.perc + uknw.perc < 15 &
         med.reads.per.MB > 2)

# Diploid: 15% <= wc+uknw < 75%
summary_hap1_diploid_filt <- novaseq250806_summary_hap1_rnp %>%
  filter(background.estimate <= 0.08 &
         background.estimate > 0 &
         wc.perc + uknw.perc < 75 &
         wc.perc + uknw.perc >= 15 &
         med.reads.per.MB > 2)

message("HAP1: ", nrow(novaseq250806_summary_hap1_rnp), " total")
message("  Haploid: ", nrow(summary_hap1_haploid_filt))
message("  Diploid: ", nrow(summary_hap1_diploid_filt))

# ---- Save cell count table ----
cell_counts_earlier <- data.frame(
  dataset = c("p20250808_xrcc1_lig3_237",
              "p20250409_xrcc1", "p20250409_lig3",
              "p20250924_xrcc1", "p20250924_lig3",
              "p20250924_237", "p20250924_237_d10a", "p20250924_237_h840a",
              "HAP1_haploid", "HAP1_diploid"),
  cells_pre_filter = c(
    nrow(summary_p20250808),
    sum(novaseq250806_summary_bj_rnp$condition == "BJ_rnp_xrcc1"),
    sum(novaseq250806_summary_bj_rnp$condition == "BJ_rnp_lig3"),
    sum(summary_p20250924$condition == "bj_rnp_xrcc1"),
    sum(summary_p20250924$condition == "bj_rnp_lig3"),
    sum(summary_p20250924$condition == "bj_rnp_seq3_237"),
    sum(summary_p20250924$condition == "bj_rnp_seq3_237_d10a"),
    sum(summary_p20250924$condition == "bj_rnp_seq3_237_h840a"),
    nrow(novaseq250806_summary_hap1_rnp),
    nrow(novaseq250806_summary_hap1_rnp)
  ),
  cells_post_filter = c(
    nrow(summary_p20250808_filt),
    sum(novaseq250806_summary_bj_rnp_filt$condition == "BJ_rnp_xrcc1"),
    sum(novaseq250806_summary_bj_rnp_filt$condition == "BJ_rnp_lig3"),
    sum(summary_p20250924_filt$condition == "bj_rnp_xrcc1"),
    sum(summary_p20250924_filt$condition == "bj_rnp_lig3"),
    sum(summary_p20250924_filt$condition == "bj_rnp_seq3_237"),
    sum(summary_p20250924_filt$condition == "bj_rnp_seq3_237_d10a"),
    sum(summary_p20250924_filt$condition == "bj_rnp_seq3_237_h840a"),
    nrow(summary_hap1_haploid_filt),
    nrow(summary_hap1_diploid_filt)
  ),
  qc_threshold = 0.08
)

write.table(cell_counts_earlier, file.path(tab_dir, "cell_counts_earlier.txt"),
            sep = "\t", row.names = FALSE, quote = FALSE)
message("Cell counts saved to ", file.path(tab_dir, "cell_counts_earlier.txt"))

# ---- QC plots ----
message("Generating QC plots...")

# Background plots
bg_p20250808 <- plot_background(summary_p20250808) + ggtitle("p20250808 xrcc1+lig3+237")
bg_p20250409 <- plot_background(novaseq250806_summary_bj_rnp) + ggtitle("p20250409 xrcc1+lig3")
bg_p20250924 <- plot_background(summary_p20250924) + ggtitle("p20250924 all conditions")
bg_hap1 <- plot_background(novaseq250806_summary_hap1_rnp) + ggtitle("HAP1 yi433")

qc_bg_earlier <- bg_p20250808 + bg_p20250409 + bg_p20250924 + bg_hap1 +
  plot_layout(nrow = 2)
ggsave("QC_background_earlier.pdf", qc_bg_earlier, grDevices::cairo_pdf, fig_dir, height = 8, width = 12)
message("Saved QC_background_earlier.pdf")

# Strand-neutral plots
sn_p20250808 <- plot_strand_neutral(summary_p20250808) + ggtitle("p20250808 xrcc1+lig3+237")
sn_p20250409 <- plot_strand_neutral(novaseq250806_summary_bj_rnp) + ggtitle("p20250409 xrcc1+lig3")
sn_p20250924 <- plot_strand_neutral(summary_p20250924) + ggtitle("p20250924 all conditions")
sn_hap1 <- plot_strand_neutral(novaseq250806_summary_hap1_rnp) + ggtitle("HAP1 yi433")

qc_sn_earlier <- sn_p20250808 + sn_p20250409 + sn_p20250924 + sn_hap1 +
  plot_layout(nrow = 2)
ggsave("QC_strand_neutral_earlier.pdf", qc_sn_earlier, grDevices::cairo_pdf, fig_dir, height = 8, width = 12)
message("Saved QC_strand_neutral_earlier.pdf")

# ---- Save environment ----
save(summary_p20250808, summary_p20250808_filt,
     summary_p20250409, novaseq250806_summary_bj_rnp, novaseq250806_summary_bj_rnp_filt,
     summary_p20250924, summary_p20250924_filt,
     novaseq250806_summary_hap1_rnp, summary_hap1_haploid_filt, summary_hap1_diploid_filt,
     file = "data/summaries_earlier.RData")

message("Done: data/summaries_earlier.RData")
```
