## Supplementary material for "Genome-wide mapping of Cas9-induced sister chromatid exchange across single and over 200 genomic targets": Cas9_RNP_scripts.zip: 02_load_new10.R

```
#!/usr/bin/env Rscript
# 02_load_new10.R
# Load and QC-filter 10 new conditions (threshold 0.08)
# Produces: data/summaries_new10.RData

message("=== 02_load_new10.R ===")
source("scripts/load_env.R")

fig_dir <- "data/figures_10conditions/"
tab_dir <- "data/tables_10conditions/"

base_path <- "/u/project/yeastyin/yeastyin/projects/Cas9_RNP/data/bpR_output"

conditions_new <- c(
  "BJ_Cas9_RNP_237cuts_24hrs_16hrs",
  "BJ_Cas9_RNP_237cuts_24ws_40hrs_BrdU",
  "BJ_Cas9_RNP_Xrcc1_24hrs",
  "BJ_Cas9_RNP_Xrcc1_24wsBrdU",
  "BJ_Cas9_RNP_Lig3_24hrs",
  "BJ_Cas9_RNP_Lig3_24wsBrdU",
  "BJ_Cas9_RNP_Lig4_24hrs",
  "BJ_Cas9_RNP_Lig4_24wsBrdU",
  "BJ_Cas9_RNP_Xrcc4_24hrs",
  "BJ_Cas9_RNP_Xrcc4_24wsBrdU"
)

summaries_new <- list()
for (cond in conditions_new) {
  csv_path <- file.path(base_path, cond, "breakPoint_qc_summary_10M.csv")
  message("Loading ", cond, " from ", csv_path, " ...")
  if (!file.exists(csv_path)) {
    message("  WARNING: CSV not found, skipping")
    next
  }
  df <- read.csv(csv_path)
  df$sample <- gsub("\\.hg19\\.filt\\.bam\\.RData\\.rds$|\\.hg19\\.filt\\.bam\\.RData$", "", df$.id)
  df$library <- sub("_.*", "", df$sample)
  df$condition <- cond
  summaries_new[[cond]] <- df
  message("  ", nrow(df), " cells loaded")
}

# Filter with threshold 0.08
summaries_new_filt <- list()
for (cond in names(summaries_new)) {
  summaries_new_filt[[cond]] <- quality_filter(summaries_new[[cond]], background_estimate = 0.08)
  message(cond, ": ", nrow(summaries_new[[cond]]), " -> ", nrow(summaries_new_filt[[cond]]), " after QC (0.08)")
}

# ---- Cell count table ----
cell_counts_new <- data.frame(
  dataset = names(summaries_new),
  cells_pre_filter = sapply(summaries_new, nrow),
  cells_post_filter = sapply(summaries_new_filt, nrow),
  qc_threshold = 0.08,
  row.names = NULL
)

write.table(cell_counts_new, file.path(tab_dir, "cell_counts_new10.txt"),
            sep = "\t", row.names = FALSE, quote = FALSE)
message("Cell counts saved to ", file.path(tab_dir, "cell_counts_new10.txt"))

# ---- QC plots ----
message("Generating QC plots for new conditions...")

bg_plots <- list()
sn_plots <- list()
for (cond in names(summaries_new)) {
  short_name <- gsub("BJ_Cas9_RNP_", "", cond)
  bg_plots[[cond]] <- plot_background(summaries_new[[cond]]) + ggtitle(short_name)
  sn_plots[[cond]] <- plot_strand_neutral(summaries_new[[cond]]) + ggtitle(short_name)
}

# Background: 2 rows x 5 cols
qc_bg_new <- wrap_plots(bg_plots, nrow = 2)
ggsave("QC_background_new10.pdf", qc_bg_new, grDevices::cairo_pdf, fig_dir, height = 8, width = 20)
message("Saved QC_background_new10.pdf")

# Strand-neutral: 2 rows x 5 cols
qc_sn_new <- wrap_plots(sn_plots, nrow = 2)
ggsave("QC_strand_neutral_new10.pdf", qc_sn_new, grDevices::cairo_pdf, fig_dir, height = 8, width = 20)
message("Saved QC_strand_neutral_new10.pdf")

# ---- Save environment ----
save(summaries_new, summaries_new_filt, conditions_new,
     file = "data/summaries_new10.RData")

message("Done: data/summaries_new10.RData")
```
