## Supplementary material for "Genome-wide mapping of Cas9-induced sister chromatid exchange across single and over 200 genomic targets": Cas9_RNP_scripts.zip: 03_hap1_pileup.R

```
#!/usr/bin/env Rscript
# 03_hap1_pileup.R
# HAP1 combined haploid+diploid pileup with manual rescue breakpoints
# Produces genome-wide pileup + zoom at LIG3 (chr17) and XRCC1 (chr19)

message("=== 03_hap1_pileup.R ===")
source("scripts/load_env.R")
load("data/summaries_earlier.RData")
load("data/xrcc1_rescue.RData")
load("data/lig3_rescue.RData")

hap1_bp_dir <- "/u/project/yeastyin/chovanec/projects/novaseq250806/HAP1_RNP_XRCC1_LIG3/hg19/breakpointR_haploid/data_filtered/"

rnp_regions <- GRanges(seqnames = c(19, 17),
                       ranges = IRanges(start = c(44047463, 33307531),
                                        end   = c(44079679, 33336762)))
mcols(rnp_regions)$name <- c("XRCC1", "LIG3")

hg19_cent <- get_centromere_coordinates("hg19", ensembl = TRUE)

# ===================================================================
# 1. Load haploid breakpoints (97 cells)
# ===================================================================
message("--- Loading haploid breakpoints ---")

breakpoints_hap1_hap_grl <- breakpoints_to_list(hap1_bp_dir,
  samples = summary_hap1_haploid_filt$sample,
  num_threads = 5, filtered = FALSE)

breakpoints_hap1_hap_filt_grl <- breakpoints_to_list(hap1_bp_dir,
  samples = summary_hap1_haploid_filt$sample,
  num_threads = 5, filtered = TRUE)

# Clean sample names (unfiltered)
breakpoints_hap1_hap_gr <- unlist(breakpoints_hap1_hap_grl)
names(breakpoints_hap1_hap_gr) <- NULL
mcols(breakpoints_hap1_hap_gr)$sample <- gsub(".hg19.filt.bam.RData_filtered", "", breakpoints_hap1_hap_gr$file)
breakpoints_hap1_hap_grl <- split(breakpoints_hap1_hap_gr, breakpoints_hap1_hap_gr$sample)

# Clean sample names (filtered)
breakpoints_hap1_hap_filt_gr <- unlist(breakpoints_hap1_hap_filt_grl)
names(breakpoints_hap1_hap_filt_gr) <- NULL
mcols(breakpoints_hap1_hap_filt_gr)$sample <- gsub(".hg19.filt.bam.RData_filtered", "", breakpoints_hap1_hap_filt_gr$file)

# ===================================================================
# 2. Add rescue breakpoints for haploid cells
# ===================================================================
message("--- Adding haploid rescue breakpoints ---")

lig3_rescue_coord  <- coordinates2gr("17:33307531-33336762", genome_assembly = "hg19") %>% plyranges::stretch(6000000)
xrcc1_rescue_coord <- coordinates2gr("19:44047463-44079679", genome_assembly = "hg19") %>% plyranges::stretch(6000000)

# LIG3 haploid rescue
lig3_hap_rescue_samples <- lig3_hap1_haploid %>%
  filter(rescue_break == TRUE) %>%
  mutate(sample = sub("^([^\\.]*\\.[^\\.]*).*", "\\1", ID)) %>%
  pull(sample)

lig3_hap_ignore <- breakpoints_hap1_hap_filt_gr[breakpoints_hap1_hap_filt_gr$sample %in% lig3_hap_rescue_samples] %>%
  plyranges::filter_by_overlaps(lig3_rescue_coord) %>%
  as.data.frame() %>% pull(sample)

lig3_hap_rescue <- setdiff(lig3_hap_rescue_samples, lig3_hap_ignore)
message("  LIG3 haploid: ", length(lig3_hap_rescue_samples), " rescued, ",
        length(lig3_hap_ignore), " already have breakpoints, ",
        length(lig3_hap_rescue), " synthetic breaks added")

lig3_hap_rescue_gr <- rep(lig3_rescue_coord, length(lig3_hap_rescue))
mcols(lig3_hap_rescue_gr) <- DataFrame(
  genoT  = rep("?-?", length(lig3_hap_rescue)),
  deltaW = rep(0, length(lig3_hap_rescue)),
  file   = lig3_hap_rescue,
  sample = lig3_hap_rescue)

# XRCC1 haploid rescue
xrcc1_hap_rescue_samples <- xrcc1_hap1_haploid %>%
  filter(rescue_break == TRUE) %>%
  mutate(sample = sub("^([^\\.]*\\.[^\\.]*).*", "\\1", ID)) %>%
  pull(sample)

xrcc1_hap_ignore <- breakpoints_hap1_hap_filt_gr[breakpoints_hap1_hap_filt_gr$sample %in% xrcc1_hap_rescue_samples] %>%
  plyranges::filter_by_overlaps(xrcc1_rescue_coord) %>%
  as.data.frame() %>% pull(sample)

xrcc1_hap_rescue <- setdiff(xrcc1_hap_rescue_samples, xrcc1_hap_ignore)
message("  XRCC1 haploid: ", length(xrcc1_hap_rescue_samples), " rescued, ",
        length(xrcc1_hap_ignore), " already have breakpoints, ",
        length(xrcc1_hap_rescue), " synthetic breaks added")

xrcc1_hap_rescue_gr <- rep(xrcc1_rescue_coord, length(xrcc1_hap_rescue))
mcols(xrcc1_hap_rescue_gr) <- DataFrame(
  genoT  = rep("?-?", length(xrcc1_hap_rescue)),
  deltaW = rep(0, length(xrcc1_hap_rescue)),
  file   = xrcc1_hap_rescue,
  sample = xrcc1_hap_rescue)

# Merge rescue into haploid filtered
breakpoints_hap1_hap_filt_gr <- c(breakpoints_hap1_hap_filt_gr,
                                   lig3_hap_rescue_gr, xrcc1_hap_rescue_gr)
breakpoints_hap1_hap_filt_grl <- split(breakpoints_hap1_hap_filt_gr, breakpoints_hap1_hap_filt_gr$sample)

# ===================================================================
# 3. Load diploid breakpoints (851 cells)
#    NOTE: Original Rmd uses breakpointR_haploid path for diploid too (line 3874)
# ===================================================================
message("--- Loading diploid breakpoints ---")

breakpoints_hap1_dip_grl <- breakpoints_to_list(hap1_bp_dir,
  samples = summary_hap1_diploid_filt$sample,
  num_threads = 5, filtered = FALSE)

breakpoints_hap1_dip_filt_grl <- breakpoints_to_list(hap1_bp_dir,
  samples = summary_hap1_diploid_filt$sample,
  num_threads = 5, filtered = TRUE)

# Clean sample names
breakpoints_hap1_dip_gr <- unlist(breakpoints_hap1_dip_grl)
names(breakpoints_hap1_dip_gr) <- NULL
mcols(breakpoints_hap1_dip_gr)$sample <- gsub(".hg19.filt.bam.RData_filtered", "", breakpoints_hap1_dip_gr$file)
breakpoints_hap1_dip_grl <- split(breakpoints_hap1_dip_gr, breakpoints_hap1_dip_gr$sample)

breakpoints_hap1_dip_filt_gr <- unlist(breakpoints_hap1_dip_filt_grl)
names(breakpoints_hap1_dip_filt_gr) <- NULL
mcols(breakpoints_hap1_dip_filt_gr)$sample <- gsub(".hg19.filt.bam.RData_filtered", "", breakpoints_hap1_dip_filt_gr$file)

# ===================================================================
# 4. Add rescue breakpoints for diploid cells
# ===================================================================
message("--- Adding diploid rescue breakpoints ---")

# LIG3 diploid rescue
lig3_dip_rescue_samples <- lig3_hap1_diploid %>%
  filter(rescue_break == TRUE) %>%
  mutate(sample = sub("^([^\\.]*\\.[^\\.]*).*", "\\1", ID)) %>%
  pull(sample)

lig3_dip_ignore <- breakpoints_hap1_dip_filt_gr[breakpoints_hap1_dip_filt_gr$sample %in% lig3_dip_rescue_samples] %>%
  plyranges::filter_by_overlaps(lig3_rescue_coord) %>%
  as.data.frame() %>% pull(sample)

lig3_dip_rescue <- setdiff(lig3_dip_rescue_samples, lig3_dip_ignore)
message("  LIG3 diploid: ", length(lig3_dip_rescue_samples), " rescued, ",
        length(lig3_dip_ignore), " already have breakpoints, ",
        length(lig3_dip_rescue), " synthetic breaks added")

lig3_dip_rescue_gr <- rep(lig3_rescue_coord, length(lig3_dip_rescue))
mcols(lig3_dip_rescue_gr) <- DataFrame(
  genoT  = rep("?-?", length(lig3_dip_rescue)),
  deltaW = rep(0, length(lig3_dip_rescue)),
  file   = lig3_dip_rescue,
  sample = lig3_dip_rescue,
  ploidy = "diploid")

# XRCC1 diploid rescue
xrcc1_dip_rescue_samples <- xrcc1_hap1_diploid %>%
  filter(rescue_break == TRUE) %>%
  mutate(sample = sub("^([^\\.]*\\.[^\\.]*).*", "\\1", ID)) %>%
  pull(sample)

xrcc1_dip_ignore <- breakpoints_hap1_dip_filt_gr[breakpoints_hap1_dip_filt_gr$sample %in% xrcc1_dip_rescue_samples] %>%
  plyranges::filter_by_overlaps(xrcc1_rescue_coord) %>%
  as.data.frame() %>% pull(sample)

xrcc1_dip_rescue <- setdiff(xrcc1_dip_rescue_samples, xrcc1_dip_ignore)
message("  XRCC1 diploid: ", length(xrcc1_dip_rescue_samples), " rescued, ",
        length(xrcc1_dip_ignore), " already have breakpoints, ",
        length(xrcc1_dip_rescue), " synthetic breaks added")

xrcc1_dip_rescue_gr <- rep(xrcc1_rescue_coord, length(xrcc1_dip_rescue))
mcols(xrcc1_dip_rescue_gr) <- DataFrame(
  genoT  = rep("?-?", length(xrcc1_dip_rescue)),
  deltaW = rep(0, length(xrcc1_dip_rescue)),
  file   = xrcc1_dip_rescue,
  sample = xrcc1_dip_rescue,
  ploidy = "diploid")

# ===================================================================
# 5. Combine haploid + diploid with ploidy metadata
# ===================================================================
message("--- Combining haploid + diploid ---")

mcols(breakpoints_hap1_dip_filt_gr)$ploidy <- "diploid"
mcols(breakpoints_hap1_hap_filt_gr)$ploidy <- "haploid"

breakpoints_hap1_combined_gr <- c(breakpoints_hap1_dip_filt_gr,
                                   lig3_dip_rescue_gr, xrcc1_dip_rescue_gr,
                                   breakpoints_hap1_hap_filt_gr)

breakpoints_hap1_combined_grl <- split(breakpoints_hap1_combined_gr,
                                        breakpoints_hap1_combined_gr$sample)

message("  Combined: ", length(breakpoints_hap1_combined_grl), " samples, ",
        length(breakpoints_hap1_combined_gr), " breakpoints")

# ===================================================================
# 6. Haploid-only pileups (genome-wide)
# ===================================================================
message("--- Haploid pileups ---")

breakpoints_hap1_hap_agg_norm <- plot_aggregate_breaks(breakpoints_hap1_hap_grl,
  tilewidth = 1000000, cent = hg19_cent, hotspot_pval = NULL,
  return_individual_plots = TRUE, normalize_counts = TRUE,
  n = length(breakpoints_hap1_hap_grl))

breakpoints_hap1_hap_agg <- plot_aggregate_breaks(breakpoints_hap1_hap_grl,
  tilewidth = 1000000, cent = hg19_cent, hotspot_pval = NULL,
  return_individual_plots = TRUE, normalize_counts = FALSE,
  n = length(breakpoints_hap1_hap_grl))

breakpoints_hap1_hap_filt_agg_norm <- plot_aggregate_breaks(breakpoints_hap1_hap_filt_grl,
  tilewidth = 1000000, cent = hg19_cent, hotspot_pval = NULL,
  return_individual_plots = TRUE, normalize_counts = TRUE,
  n = length(breakpoints_hap1_hap_filt_grl))

breakpoints_hap1_hap_filt_agg <- plot_aggregate_breaks(breakpoints_hap1_hap_filt_grl,
  tilewidth = 1000000, cent = hg19_cent, hotspot_pval = NULL,
  return_individual_plots = TRUE, normalize_counts = FALSE,
  n = length(breakpoints_hap1_hap_filt_grl), regions = rnp_regions)

# ===================================================================
# 7. Diploid + combined pileups (genome-wide)
# ===================================================================
message("--- Diploid + combined pileups ---")

breakpoints_hap1_dip_agg_norm <- plot_aggregate_breaks(breakpoints_hap1_dip_grl,
  tilewidth = 1000000, cent = hg19_cent, hotspot_pval = NULL,
  return_individual_plots = TRUE, normalize_counts = TRUE,
  n = length(breakpoints_hap1_dip_grl))

breakpoints_hap1_dip_agg <- plot_aggregate_breaks(breakpoints_hap1_dip_grl,
  tilewidth = 1000000, cent = hg19_cent, hotspot_pval = NULL,
  return_individual_plots = TRUE, normalize_counts = FALSE,
  n = length(breakpoints_hap1_dip_grl))

breakpoints_hap1_combined_agg_norm <- plot_aggregate_breaks(breakpoints_hap1_combined_grl,
  tilewidth = 1000000, cent = hg19_cent, hotspot_pval = NULL,
  return_individual_plots = TRUE, normalize_counts = TRUE,
  n = length(breakpoints_hap1_combined_grl))

breakpoints_hap1_combined_agg <- plot_aggregate_breaks(breakpoints_hap1_combined_grl,
  tilewidth = 1000000, cent = hg19_cent, hotspot_pval = NULL,
  return_individual_plots = TRUE, normalize_counts = FALSE,
  n = length(breakpoints_hap1_combined_grl), regions = rnp_regions)

# ===================================================================
# 8. Genome-wide pileup figure (matches original fig_HAP1_433_pileup.pdf)
# ===================================================================
message("--- Genome-wide pileup figure ---")

p_hap1_pileup <- breakpoints_hap1_dip_agg_norm[[1]] +
  breakpoints_hap1_dip_agg[[1]] +
  breakpoints_hap1_combined_agg_norm[[1]] +
  breakpoints_hap1_combined_agg[[1]] +
  breakpoints_hap1_combined_agg[[4]] +
  breakpoints_hap1_combined_agg[[5]] +
  plot_layout(nrow = 6, heights = c(4, 4, 4, 4, 1, 1))

ggsave("pileup_HAP1_433_genome.pdf", p_hap1_pileup, grDevices::cairo_pdf, fig_dir,
       height = 5, width = 12)
message("Saved pileup_HAP1_433_genome.pdf")

# ===================================================================
# 9. Zoom LIG3 (chr17)
# ===================================================================
message("--- LIG3 zoom (chr17) ---")

bp_zoom_lig3 <- breakpoints_hap1_combined_gr
seqlevels(bp_zoom_lig3, pruning.mode = "coarse") <- seqlevels(seqinfo(bp_zoom_lig3)["17"])
seqinfo(bp_zoom_lig3) <- seqinfo(bp_zoom_lig3)["17"]

pileup_hap1_zoom_lig3 <- plot_aggregate_breaks(GRangesList(bp_zoom_lig3),
  tilewidth = 1000000, cent = hg19_cent, hotspot_pval = 1e-08,
  return_individual_plots = TRUE, normalize_counts = TRUE,
  n = length(unique(bp_zoom_lig3$sample)), regions = rnp_regions)

chr_max <- max(pileup_hap1_zoom_lig3[[1]]$layers[[2]]$data$end.genome)

p_zoom_lig3 <- (pileup_hap1_zoom_lig3[[1]] +
  geom_vline(xintercept = chr_max, size = 0.3, col = "black") +
  theme(plot.title = element_text(size = 11)) + legend_theme) +
  (pileup_hap1_zoom_lig3[[4]] +
  geom_vline(xintercept = chr_max, size = 0.3, col = "black") + legend_theme) +
  plot_layout(ncol = 1, heights = c(5, 1))

ggsave("pileup_zoom_HAP1_lig3.pdf", p_zoom_lig3, grDevices::cairo_pdf, fig_dir,
       height = 2, width = 2.1)
message("Saved pileup_zoom_HAP1_lig3.pdf")

# ===================================================================
# 10. Zoom XRCC1 (chr19)
# ===================================================================
message("--- XRCC1 zoom (chr19) ---")

bp_zoom_xrcc1 <- breakpoints_hap1_combined_gr
seqlevels(bp_zoom_xrcc1, pruning.mode = "coarse") <- seqlevels(seqinfo(bp_zoom_xrcc1)["19"])
seqinfo(bp_zoom_xrcc1) <- seqinfo(bp_zoom_xrcc1)["19"]

pileup_hap1_zoom_xrcc1 <- plot_aggregate_breaks(GRangesList(bp_zoom_xrcc1),
  tilewidth = 1000000, cent = hg19_cent, hotspot_pval = 1e-08,
  return_individual_plots = TRUE, normalize_counts = TRUE,
  n = length(unique(bp_zoom_xrcc1$sample)), regions = rnp_regions)

chr_max <- max(pileup_hap1_zoom_xrcc1[[1]]$layers[[2]]$data$end.genome)

p_zoom_xrcc1 <- (pileup_hap1_zoom_xrcc1[[1]] +
  geom_vline(xintercept = chr_max, size = 0.3, col = "black") +
  theme(plot.title = element_text(size = 11)) + legend_theme) +
  (pileup_hap1_zoom_xrcc1[[4]] +
  geom_vline(xintercept = chr_max, size = 0.3, col = "black") + legend_theme) +
  plot_layout(ncol = 1, heights = c(5, 1))

ggsave("pileup_zoom_HAP1_xrcc1.pdf", p_zoom_xrcc1, grDevices::cairo_pdf, fig_dir,
       height = 2, width = 2.1)
message("Saved pileup_zoom_HAP1_xrcc1.pdf")

message("Done: 03_hap1_pileup.R")
```
