## Supplementary material for "Genome-wide mapping of Cas9-induced sister chromatid exchange across single and over 200 genomic targets": Cas9_RNP_scripts.zip: 04_07_rescue_update_yi564.R

```
#!/usr/bin/env Rscript
# 04_07_rescue_update_yi564.R
# Re-run rescue analysis ONLY for the new siRNA conditions that got yi564 cells.
# Preserves old condition results from existing .RData files.
# Covers: XRCC1, LIG3, LIG4, XRCC4

message("=== 04_07_rescue_update_yi564.R ===")
source("scripts/load_env.R")
load("data/summaries_earlier.RData")
load("data/summaries_new10.RData")

hg19_cent <- get_centromere_coordinates("hg19", ensembl = TRUE)

# ===================================================================
# XRCC1 rescue (chr19:44047463-44079679)
# ===================================================================
message("\n========== XRCC1 ==========")
load("data/xrcc1_rescue.RData")
xrcc1_coord_str <- "19:44047463-44079679"

message("--- Re-running Xrcc1_24hrs ---")
xrcc1_new_24hrs <- run_check_location_for_breakpoints(
  "/u/project/yeastyin/yeastyin/projects/Cas9_RNP/data/bpR_output/BJ_Cas9_RNP_Xrcc1_24hrs/data_filtered_10M/",
  coord = xrcc1_coord_str,
  samples = summaries_new_filt[["BJ_Cas9_RNP_Xrcc1_24hrs"]]$sample,
  num_threads = 10, bin_width = 10000000, dist_from_gr = 3000000, type = "diploid")
message("  Xrcc1_24hrs rescue rate: ", sum(xrcc1_new_24hrs$rescue_break), "/", nrow(xrcc1_new_24hrs))

message("--- Re-running Xrcc1_24wsBrdU ---")
xrcc1_new_24wsBrdU <- run_check_location_for_breakpoints(
  "/u/project/yeastyin/yeastyin/projects/Cas9_RNP/data/bpR_output/BJ_Cas9_RNP_Xrcc1_24wsBrdU/data_filtered_10M/",
  coord = xrcc1_coord_str,
  samples = summaries_new_filt[["BJ_Cas9_RNP_Xrcc1_24wsBrdU"]]$sample,
  num_threads = 10, bin_width = 10000000, dist_from_gr = 3000000, type = "diploid")
message("  Xrcc1_24wsBrdU rescue rate: ", sum(xrcc1_new_24wsBrdU$rescue_break), "/", nrow(xrcc1_new_24wsBrdU))

# Update summary
xrcc1_rescue_summary <- data.frame(
  condition = c("p20250808_xrcc1_lig3_237", "p20250409_xrcc1",
                "p20250924_xrcc1", "HAP1_haploid", "HAP1_diploid",
                "Xrcc1_24hrs", "Xrcc1_24wsBrdU"),
  total_cells = c(nrow(xrcc1_p20250808), nrow(xrcc1_p20250409),
                  nrow(xrcc1_p20250924), nrow(xrcc1_hap1_haploid), nrow(xrcc1_hap1_diploid),
                  nrow(xrcc1_new_24hrs), nrow(xrcc1_new_24wsBrdU)),
  rescued = c(sum(xrcc1_p20250808$rescue_break), sum(xrcc1_p20250409$rescue_break),
              sum(xrcc1_p20250924$rescue_break), sum(xrcc1_hap1_haploid$rescue_break),
              sum(xrcc1_hap1_diploid$rescue_break),
              sum(xrcc1_new_24hrs$rescue_break), sum(xrcc1_new_24wsBrdU$rescue_break))
)
xrcc1_rescue_summary$rescue_rate <- round(xrcc1_rescue_summary$rescued / xrcc1_rescue_summary$total_cells, 4)
write.table(xrcc1_rescue_summary, file.path(tab_dir, "xrcc1_rescue_summary.txt"),
            sep = "\t", row.names = FALSE, quote = FALSE)
print(xrcc1_rescue_summary)

save(xrcc1_p20250808, xrcc1_p20250409, xrcc1_p20250924,
     xrcc1_hap1_haploid, xrcc1_hap1_diploid,
     xrcc1_new_24hrs, xrcc1_new_24wsBrdU,
     xrcc1_rescue_summary,
     bp_p20250808_filt_rescue_grl,
     bp_xrcc1_409_filt_rescue_grl,
     bp_xrcc1_924_filt_rescue_grl,
     file = "data/xrcc1_rescue.RData")
message("Saved data/xrcc1_rescue.RData")

# ===================================================================
# LIG3 rescue (chr17:33307531-33336762)
# ===================================================================
message("\n========== LIG3 ==========")
load("data/lig3_rescue.RData")
lig3_coord_str <- "17:33307531-33336762"

message("--- Re-running Lig3_24hrs ---")
lig3_new_24hrs <- run_check_location_for_breakpoints(
  "/u/project/yeastyin/yeastyin/projects/Cas9_RNP/data/bpR_output/BJ_Cas9_RNP_Lig3_24hrs/data_filtered_10M/",
  coord = lig3_coord_str,
  samples = summaries_new_filt[["BJ_Cas9_RNP_Lig3_24hrs"]]$sample,
  num_threads = 10, bin_width = 10000000, dist_from_gr = 3000000, type = "diploid")
message("  Lig3_24hrs rescue rate: ", sum(lig3_new_24hrs$rescue_break), "/", nrow(lig3_new_24hrs))

message("--- Re-running Lig3_24wsBrdU ---")
lig3_new_24wsBrdU <- run_check_location_for_breakpoints(
  "/u/project/yeastyin/yeastyin/projects/Cas9_RNP/data/bpR_output/BJ_Cas9_RNP_Lig3_24wsBrdU/data_filtered_10M/",
  coord = lig3_coord_str,
  samples = summaries_new_filt[["BJ_Cas9_RNP_Lig3_24wsBrdU"]]$sample,
  num_threads = 10, bin_width = 10000000, dist_from_gr = 3000000, type = "diploid")
message("  Lig3_24wsBrdU rescue rate: ", sum(lig3_new_24wsBrdU$rescue_break), "/", nrow(lig3_new_24wsBrdU))

# Update summary (need to check what variables exist in lig3_rescue.RData)
lig3_rescue_summary <- data.frame(
  condition = c("p20250808_xrcc1_lig3_237", "p20250409_lig3",
                "p20250924_lig3", "HAP1_haploid", "HAP1_diploid",
                "Lig3_24hrs", "Lig3_24wsBrdU"),
  total_cells = c(nrow(lig3_p20250808), nrow(lig3_p20250409),
                  nrow(lig3_p20250924), nrow(lig3_hap1_haploid), nrow(lig3_hap1_diploid),
                  nrow(lig3_new_24hrs), nrow(lig3_new_24wsBrdU)),
  rescued = c(sum(lig3_p20250808$rescue_break), sum(lig3_p20250409$rescue_break),
              sum(lig3_p20250924$rescue_break), sum(lig3_hap1_haploid$rescue_break),
              sum(lig3_hap1_diploid$rescue_break),
              sum(lig3_new_24hrs$rescue_break), sum(lig3_new_24wsBrdU$rescue_break))
)
lig3_rescue_summary$rescue_rate <- round(lig3_rescue_summary$rescued / lig3_rescue_summary$total_cells, 4)
write.table(lig3_rescue_summary, file.path(tab_dir, "lig3_rescue_summary.txt"),
            sep = "\t", row.names = FALSE, quote = FALSE)
print(lig3_rescue_summary)

save(lig3_p20250808, lig3_p20250409, lig3_p20250924,
     lig3_hap1_haploid, lig3_hap1_diploid,
     lig3_new_24hrs, lig3_new_24wsBrdU,
     lig3_rescue_summary,
     bp_p20250808_filt_rescue_grl,
     bp_lig3_409_filt_rescue_grl,
     bp_lig3_924_filt_rescue_grl,
     file = "data/lig3_rescue.RData")
message("Saved data/lig3_rescue.RData")

# ===================================================================
# LIG4 rescue (chr13:108859790-108870697)
# Only has new conditions (no old p20250924/HAP1 data)
# ===================================================================
message("\n========== LIG4 ==========")
lig4_coord_str <- "13:108859790-108870697"

message("--- Re-running Lig4_24hrs ---")
lig4_new_24hrs <- run_check_location_for_breakpoints(
  "/u/project/yeastyin/yeastyin/projects/Cas9_RNP/data/bpR_output/BJ_Cas9_RNP_Lig4_24hrs/data_filtered_10M/",
  coord = lig4_coord_str,
  samples = summaries_new_filt[["BJ_Cas9_RNP_Lig4_24hrs"]]$sample,
  num_threads = 10, bin_width = 10000000, dist_from_gr = 3000000, type = "diploid")
message("  Lig4_24hrs rescue rate: ", sum(lig4_new_24hrs$rescue_break), "/", nrow(lig4_new_24hrs))

message("--- Re-running Lig4_24wsBrdU ---")
lig4_new_24wsBrdU <- run_check_location_for_breakpoints(
  "/u/project/yeastyin/yeastyin/projects/Cas9_RNP/data/bpR_output/BJ_Cas9_RNP_Lig4_24wsBrdU/data_filtered_10M/",
  coord = lig4_coord_str,
  samples = summaries_new_filt[["BJ_Cas9_RNP_Lig4_24wsBrdU"]]$sample,
  num_threads = 10, bin_width = 10000000, dist_from_gr = 3000000, type = "diploid")
message("  Lig4_24wsBrdU rescue rate: ", sum(lig4_new_24wsBrdU$rescue_break), "/", nrow(lig4_new_24wsBrdU))

lig4_rescue_summary <- data.frame(
  condition = c("Lig4_24hrs", "Lig4_24wsBrdU"),
  total_cells = c(nrow(lig4_new_24hrs), nrow(lig4_new_24wsBrdU)),
  rescued = c(sum(lig4_new_24hrs$rescue_break), sum(lig4_new_24wsBrdU$rescue_break))
)
lig4_rescue_summary$rescue_rate <- round(lig4_rescue_summary$rescued / lig4_rescue_summary$total_cells, 4)
write.table(lig4_rescue_summary, file.path(tab_dir, "lig4_rescue_summary.txt"),
            sep = "\t", row.names = FALSE, quote = FALSE)
print(lig4_rescue_summary)

save(lig4_new_24hrs, lig4_new_24wsBrdU,
     lig4_rescue_summary,
     file = "data/lig4_rescue.RData")
message("Saved data/lig4_rescue.RData")

# ===================================================================
# XRCC4 rescue (chr5:82367495-82643756)
# Only has new conditions (no old p20250924/HAP1 data)
# ===================================================================
message("\n========== XRCC4 ==========")
xrcc4_coord_str <- "5:82367495-82643756"

message("--- Re-running Xrcc4_24hrs ---")
xrcc4_new_24hrs <- run_check_location_for_breakpoints(
  "/u/project/yeastyin/yeastyin/projects/Cas9_RNP/data/bpR_output/BJ_Cas9_RNP_Xrcc4_24hrs/data_filtered_10M/",
  coord = xrcc4_coord_str,
  samples = summaries_new_filt[["BJ_Cas9_RNP_Xrcc4_24hrs"]]$sample,
  num_threads = 10, bin_width = 10000000, dist_from_gr = 3000000, type = "diploid")
message("  Xrcc4_24hrs rescue rate: ", sum(xrcc4_new_24hrs$rescue_break), "/", nrow(xrcc4_new_24hrs))

message("--- Re-running Xrcc4_24wsBrdU ---")
xrcc4_new_24wsBrdU <- run_check_location_for_breakpoints(
  "/u/project/yeastyin/yeastyin/projects/Cas9_RNP/data/bpR_output/BJ_Cas9_RNP_Xrcc4_24wsBrdU/data_filtered_10M/",
  coord = xrcc4_coord_str,
  samples = summaries_new_filt[["BJ_Cas9_RNP_Xrcc4_24wsBrdU"]]$sample,
  num_threads = 10, bin_width = 10000000, dist_from_gr = 3000000, type = "diploid")
message("  Xrcc4_24wsBrdU rescue rate: ", sum(xrcc4_new_24wsBrdU$rescue_break), "/", nrow(xrcc4_new_24wsBrdU))

xrcc4_rescue_summary <- data.frame(
  condition = c("Xrcc4_24hrs", "Xrcc4_24wsBrdU"),
  total_cells = c(nrow(xrcc4_new_24hrs), nrow(xrcc4_new_24wsBrdU)),
  rescued = c(sum(xrcc4_new_24hrs$rescue_break), sum(xrcc4_new_24wsBrdU$rescue_break))
)
xrcc4_rescue_summary$rescue_rate <- round(xrcc4_rescue_summary$rescued / xrcc4_rescue_summary$total_cells, 4)
write.table(xrcc4_rescue_summary, file.path(tab_dir, "xrcc4_rescue_summary.txt"),
            sep = "\t", row.names = FALSE, quote = FALSE)
print(xrcc4_rescue_summary)

save(xrcc4_new_24hrs, xrcc4_new_24wsBrdU,
     xrcc4_rescue_summary,
     file = "data/xrcc4_rescue.RData")
message("Saved data/xrcc4_rescue.RData")

message("\n=== Done: All rescue analyses updated ===")
```
