## Supplementary material for "Genome-wide mapping of Cas9-induced sister chromatid exchange across single and over 200 genomic targets": Cas9_RNP_scripts.zip: 04_xrcc1_rescue.R

```
#!/usr/bin/env Rscript
# 04_xrcc1_rescue.R
# Rescue breakpoint analysis at XRCC1 locus (chr19:44047463-44079679)
# Datasets with XRCC1 cut: p20250808, p20250409_xrcc1, p20250924_xrcc1, HAP1,
#   + new Xrcc1_24hrs, Xrcc1_24wsBrdU

message("=== 04_xrcc1_rescue.R ===")
source("scripts/load_env.R")
load("data/summaries_earlier.RData")
load("data/summaries_new10.RData")

xrcc1_coord_str <- "19:44047463-44079679"

# ===================================================================
# 1. p20250808: xrcc1+lig3+237 (has XRCC1 cut)
# ===================================================================
message("--- p20250808 XRCC1 rescue ---")

xrcc1_p20250808 <- run_check_location_for_breakpoints(
  c("/u/project/yeastyin/chovanec/projects/novaseq250815/bj_rnp_xrcc1_lig3_seq3_237/human/breakpointR_diploid/data_filtered/",
    "/u/project/yeastyin/chovanec/projects/novaseq250919/bj_rnp_xrcc1_lig3_seq3_237/human/breakpointR_diploid/data_filtered/"),
  coord = xrcc1_coord_str, samples = summary_p20250808_filt$sample,
  num_threads = 10, bin_width = 10000000, dist_from_gr = 3000000, type = "diploid")

message("p20250808 XRCC1 rescue rate: ", sum(xrcc1_p20250808$rescue_break), "/", nrow(xrcc1_p20250808))

# Pileup with rescue
bp_p20250808_filt_grl <- breakpoints_to_list(
  c("/u/project/yeastyin/chovanec/projects/novaseq250815/bj_rnp_xrcc1_lig3_seq3_237/human/breakpointR_diploid/data_filtered/",
    "/u/project/yeastyin/chovanec/projects/novaseq250919/bj_rnp_xrcc1_lig3_seq3_237/human/breakpointR_diploid/data_filtered/"),
  samples = summary_p20250808_filt$sample, num_threads = 10, filtered = TRUE)

bp_p20250808_filt_gr <- unlist(bp_p20250808_filt_grl)
names(bp_p20250808_filt_gr) <- NULL
mcols(bp_p20250808_filt_gr)$sample <- gsub("\\.human\\.filt\\.bam\\.RData_filtered|\\.hg19\\.filt\\.bam\\.RData_filtered", "", bp_p20250808_filt_gr$file)

bp_p20250808_filt_rescue_gr <- add_rescue_breaks(bp_p20250808_filt_gr, xrcc1_p20250808, xrcc1_coord_str)
bp_p20250808_filt_rescue_grl <- split(bp_p20250808_filt_rescue_gr, bp_p20250808_filt_rescue_gr$sample)

hg19_cent <- get_centromere_coordinates("hg19", ensembl = TRUE)
rnp_xrcc1 <- GRanges(seqnames = 19, ranges = IRanges(start = 44047463, end = 44079679))
mcols(rnp_xrcc1)$name <- "XRCC1"

# Genome-wide pileup
pileup_p20250808_xrcc1 <- plot_aggregate_breaks(bp_p20250808_filt_rescue_grl,
  tilewidth = 1000000, cent = hg19_cent, hotspot_pval = NULL,
  return_individual_plots = TRUE, normalize_counts = TRUE,
  n = length(bp_p20250808_filt_rescue_grl), regions = rnp_xrcc1)

# Zoom chr19
bp_zoom <- bp_p20250808_filt_rescue_gr
seqlevels(bp_zoom, pruning.mode = "coarse") <- seqlevels(seqinfo(bp_zoom)["19"])
seqinfo(bp_zoom) <- seqinfo(bp_zoom)["19"]

pileup_zoom_p20250808_xrcc1 <- plot_aggregate_breaks(GRangesList(bp_zoom),
  tilewidth = 1000000, cent = hg19_cent, hotspot_pval = 1e-08,
  return_individual_plots = TRUE, normalize_counts = TRUE,
  n = length(unique(bp_zoom$sample)), regions = rnp_xrcc1)

chr_max <- max(pileup_zoom_p20250808_xrcc1[[1]]$layers[[2]]$data$end.genome)
p_zoom <- (pileup_zoom_p20250808_xrcc1[[1]] + geom_vline(xintercept = chr_max, size = 0.3) + legend_theme) +
  (pileup_zoom_p20250808_xrcc1[[4]] + geom_vline(xintercept = chr_max, size = 0.3) + legend_theme) +
  plot_layout(ncol = 1, heights = c(5, 1))
ggsave("pileup_zoom_p20250808_xrcc1.pdf", p_zoom, grDevices::cairo_pdf, fig_dir, height = 2, width = 2.1)
message("Saved pileup_zoom_p20250808_xrcc1.pdf")

# ===================================================================
# 2. p20250409: BJ_rnp_xrcc1 (yi441 via novaseq250806)
# ===================================================================
message("--- p20250409 XRCC1 rescue ---")

xrcc1_p20250409 <- run_check_location_for_breakpoints(
  "/u/project/yeastyin/chovanec/projects/novaseq250806/BJ5ta_RNP_XRCC1_LIG3/hg19/breakpointR_diploid/data_filtered/",
  coord = xrcc1_coord_str,
  samples = novaseq250806_summary_bj_rnp_filt$sample[novaseq250806_summary_bj_rnp_filt$condition == "BJ_rnp_xrcc1"],
  num_threads = 10, bin_width = 10000000, dist_from_gr = 3000000, type = "diploid")

message("p20250409 XRCC1 rescue rate: ", sum(xrcc1_p20250409$rescue_break), "/", nrow(xrcc1_p20250409))

# Pileup
bp_xrcc1_409_filt_grl <- breakpoints_to_list(
  "/u/project/yeastyin/chovanec/projects/novaseq250806/BJ5ta_RNP_XRCC1_LIG3/hg19/breakpointR_diploid/data_filtered/",
  samples = novaseq250806_summary_bj_rnp_filt %>% filter(condition == "BJ_rnp_xrcc1") %>% pull(sample),
  num_threads = 5, filtered = TRUE)

bp_xrcc1_409_filt_gr <- unlist(bp_xrcc1_409_filt_grl)
names(bp_xrcc1_409_filt_gr) <- NULL
mcols(bp_xrcc1_409_filt_gr)$sample <- gsub("\\.hg19\\.filt\\.bam\\.RData_filtered", "", bp_xrcc1_409_filt_gr$file)

bp_xrcc1_409_filt_rescue_gr <- add_rescue_breaks(bp_xrcc1_409_filt_gr, xrcc1_p20250409, xrcc1_coord_str)
bp_xrcc1_409_filt_rescue_grl <- split(bp_xrcc1_409_filt_rescue_gr, bp_xrcc1_409_filt_rescue_gr$sample)

# Zoom chr19
bp_zoom <- bp_xrcc1_409_filt_gr  # no rescue for zoom per original code
seqlevels(bp_zoom, pruning.mode = "coarse") <- seqlevels(seqinfo(bp_zoom)["19"])
seqinfo(bp_zoom) <- seqinfo(bp_zoom)["19"]

pileup_zoom_p20250409_xrcc1 <- plot_aggregate_breaks(GRangesList(bp_zoom),
  tilewidth = 1000000, cent = hg19_cent, hotspot_pval = 1e-08,
  return_individual_plots = TRUE, normalize_counts = TRUE,
  n = length(bp_xrcc1_409_filt_rescue_grl), regions = rnp_xrcc1)

chr_max <- max(pileup_zoom_p20250409_xrcc1[[1]]$layers[[2]]$data$end.genome)
p_zoom <- (pileup_zoom_p20250409_xrcc1[[1]] + geom_vline(xintercept = chr_max, size = 0.3) + legend_theme) +
  (pileup_zoom_p20250409_xrcc1[[4]] + geom_vline(xintercept = chr_max, size = 0.3) + legend_theme) +
  plot_layout(ncol = 1, heights = c(5, 1))
ggsave("pileup_zoom_p20250409_xrcc1.pdf", p_zoom, grDevices::cairo_pdf, fig_dir, height = 2, width = 2.1)
message("Saved pileup_zoom_p20250409_xrcc1.pdf")

# ===================================================================
# 3. p20250924: bj_rnp_xrcc1 (yi478)
# ===================================================================
message("--- p20250924 XRCC1 rescue ---")

xrcc1_p20250924 <- run_check_location_for_breakpoints(
  "/u/project/yeastyin/chovanec/projects/novaseq251007/BJ_RNP/human/breakpointR_diploid/data_filtered/",
  coord = xrcc1_coord_str,
  samples = summary_p20250924_filt %>% filter(condition == "bj_rnp_xrcc1") %>% pull(sample),
  num_threads = 10, bin_width = 10000000, dist_from_gr = 3000000, type = "diploid")

message("p20250924 XRCC1 rescue rate: ", sum(xrcc1_p20250924$rescue_break), "/", nrow(xrcc1_p20250924))

# Pileup
bp_xrcc1_924_filt_grl <- breakpoints_to_list(
  "/u/project/yeastyin/chovanec/projects/novaseq251007/BJ_RNP/human/breakpointR_diploid/data_filtered/",
  samples = summary_p20250924_filt %>% filter(condition == "bj_rnp_xrcc1") %>% pull(sample),
  num_threads = 5, filtered = TRUE)

bp_xrcc1_924_filt_gr <- unlist(bp_xrcc1_924_filt_grl)
names(bp_xrcc1_924_filt_gr) <- NULL
mcols(bp_xrcc1_924_filt_gr)$sample <- gsub("\\.hg19\\.filt\\.bam\\.RData_filtered", "", bp_xrcc1_924_filt_gr$file)

bp_xrcc1_924_filt_rescue_gr <- add_rescue_breaks(bp_xrcc1_924_filt_gr, xrcc1_p20250924, xrcc1_coord_str)
bp_xrcc1_924_filt_rescue_grl <- split(bp_xrcc1_924_filt_rescue_gr, bp_xrcc1_924_filt_rescue_gr$sample)

# Zoom chr19
bp_zoom <- bp_xrcc1_924_filt_gr
seqlevels(bp_zoom, pruning.mode = "coarse") <- seqlevels(seqinfo(bp_zoom)["19"])
seqinfo(bp_zoom) <- seqinfo(bp_zoom)["19"]

pileup_zoom_p20250924_xrcc1 <- plot_aggregate_breaks(GRangesList(bp_zoom),
  tilewidth = 1000000, cent = hg19_cent, hotspot_pval = 1e-08,
  return_individual_plots = TRUE, normalize_counts = TRUE,
  n = length(bp_xrcc1_924_filt_rescue_grl), regions = rnp_xrcc1)

chr_max <- max(pileup_zoom_p20250924_xrcc1[[1]]$layers[[2]]$data$end.genome)
p_zoom <- (pileup_zoom_p20250924_xrcc1[[1]] + geom_vline(xintercept = chr_max, size = 0.3) + legend_theme) +
  (pileup_zoom_p20250924_xrcc1[[4]] + geom_vline(xintercept = chr_max, size = 0.3) + legend_theme) +
  plot_layout(ncol = 1, heights = c(5, 1))
ggsave("pileup_zoom_p20250924_xrcc1.pdf", p_zoom, grDevices::cairo_pdf, fig_dir, height = 2, width = 2.1)
message("Saved pileup_zoom_p20250924_xrcc1.pdf")

# ===================================================================
# 4. HAP1 haploid
# ===================================================================
message("--- HAP1 haploid XRCC1 rescue ---")

xrcc1_hap1_haploid <- run_check_location_for_breakpoints(
  "/u/project/yeastyin/chovanec/projects/novaseq250806/HAP1_RNP_XRCC1_LIG3/hg19/breakpointR_haploid/data_filtered/",
  coord = xrcc1_coord_str, samples = summary_hap1_haploid_filt$sample,
  num_threads = 10, bin_width = 10000000, dist_from_gr = 3000000, type = "haploid")

message("HAP1 haploid XRCC1 rescue rate: ", sum(xrcc1_hap1_haploid$rescue_break), "/", nrow(xrcc1_hap1_haploid))

# ===================================================================
# 5. HAP1 diploid
# ===================================================================
message("--- HAP1 diploid XRCC1 rescue ---")

xrcc1_hap1_diploid <- run_check_location_for_breakpoints(
  "/u/project/yeastyin/chovanec/projects/novaseq250806/HAP1_RNP_XRCC1_LIG3/hg19/breakpointR_diploid/data_filtered/",
  coord = xrcc1_coord_str, samples = summary_hap1_diploid_filt$sample,
  num_threads = 10, bin_width = 10000000, dist_from_gr = 3000000, type = "diploid")

message("HAP1 diploid XRCC1 rescue rate: ", sum(xrcc1_hap1_diploid$rescue_break), "/", nrow(xrcc1_hap1_diploid))

# ===================================================================
# 6. New: BJ_Cas9_RNP_Xrcc1_24hrs
# ===================================================================
message("--- New Xrcc1_24hrs rescue ---")

xrcc1_new_24hrs <- run_check_location_for_breakpoints(
  "/u/project/yeastyin/yeastyin/projects/Cas9_RNP/data/bpR_output/BJ_Cas9_RNP_Xrcc1_24hrs/data_filtered_10M/",
  coord = xrcc1_coord_str,
  samples = summaries_new_filt[["BJ_Cas9_RNP_Xrcc1_24hrs"]]$sample,
  num_threads = 10, bin_width = 10000000, dist_from_gr = 3000000, type = "diploid")

message("Xrcc1_24hrs rescue rate: ", sum(xrcc1_new_24hrs$rescue_break), "/", nrow(xrcc1_new_24hrs))

# ===================================================================
# 7. New: BJ_Cas9_RNP_Xrcc1_24wsBrdU
# ===================================================================
message("--- New Xrcc1_24wsBrdU rescue ---")

xrcc1_new_24wsBrdU <- run_check_location_for_breakpoints(
  "/u/project/yeastyin/yeastyin/projects/Cas9_RNP/data/bpR_output/BJ_Cas9_RNP_Xrcc1_24wsBrdU/data_filtered_10M/",
  coord = xrcc1_coord_str,
  samples = summaries_new_filt[["BJ_Cas9_RNP_Xrcc1_24wsBrdU"]]$sample,
  num_threads = 10, bin_width = 10000000, dist_from_gr = 3000000, type = "diploid")

message("Xrcc1_24wsBrdU rescue rate: ", sum(xrcc1_new_24wsBrdU$rescue_break), "/", nrow(xrcc1_new_24wsBrdU))

# ===================================================================
# Summary rescue table for XRCC1
# ===================================================================
xrcc1_rescue_summary <- data.frame(
  condition = c("p20250808_xrcc1_lig3_237", "p20250409_xrcc1",
                "p20250924_xrcc1", "HAP1_haploid", "HAP1_diploid",
                "Xrcc1_24hrs", "Xrcc1_24wsBrdU"),
  total_cells = c(nrow(xrcc1_p20250808), nrow(xrcc1_p20250409),
                  nrow(xrcc1_p20250924), nrow(xrcc1_hap1_haploid), nrow(xrcc1_hap1_diploid),
                  nrow(xrcc1_new_24hrs), nrow(xrcc1_new_24wsBrdU)),
  rescued = c(sum(xrcc1_p20250808$rescue_break), sum(xrcc1_p20250409$rescue_break),
              sum(xrcc1_p20250924$rescue_break), sum(xrcc1_hap1_haploid$rescue_break),
              sum(xrcc1_hap1_diploid$rescue_break),
              sum(xrcc1_new_24hrs$rescue_break), sum(xrcc1_new_24wsBrdU$rescue_break))
)
xrcc1_rescue_summary$rescue_rate <- round(xrcc1_rescue_summary$rescued / xrcc1_rescue_summary$total_cells, 4)

write.table(xrcc1_rescue_summary, file.path(tab_dir, "xrcc1_rescue_summary.txt"),
            sep = "\t", row.names = FALSE, quote = FALSE)
message("Saved xrcc1_rescue_summary.txt")
print(xrcc1_rescue_summary)

# ---- Save ----
save(xrcc1_p20250808, xrcc1_p20250409, xrcc1_p20250924,
     xrcc1_hap1_haploid, xrcc1_hap1_diploid,
     xrcc1_new_24hrs, xrcc1_new_24wsBrdU,
     xrcc1_rescue_summary,
     bp_p20250808_filt_rescue_grl,
     bp_xrcc1_409_filt_rescue_grl,
     bp_xrcc1_924_filt_rescue_grl,
     file = "data/xrcc1_rescue.RData")

message("Done: data/xrcc1_rescue.RData")
```
