## Supplementary material for "Genome-wide mapping of Cas9-induced sister chromatid exchange across single and over 200 genomic targets": Cas9_RNP_scripts.zip: 04b_xrcc1_new_pileup.R

```
#!/usr/bin/env Rscript
# 04b_xrcc1_new_pileup.R
# Pileup zoom plots for new XRCC1 conditions: Xrcc1_24hrs, Xrcc1_24wsBrdU

message("=== 04b_xrcc1_new_pileup.R ===")
source("scripts/load_env.R")
load("data/summaries_new10.RData")
load("data/xrcc1_rescue.RData")

xrcc1_coord_str <- "19:44047463-44079679"
hg19_cent <- get_centromere_coordinates("hg19", ensembl = TRUE)
rnp_xrcc1 <- GRanges(seqnames = 19, ranges = IRanges(start = 44047463, end = 44079679))
mcols(rnp_xrcc1)$name <- "XRCC1"

# ===================================================================
# 1. BJ_Cas9_RNP_Xrcc1_24hrs
# ===================================================================
message("--- Xrcc1_24hrs pileup ---")

bp_xrcc1_24hrs_filt_grl <- breakpoints_to_list(
  "/u/project/yeastyin/yeastyin/projects/Cas9_RNP/data/bpR_output/BJ_Cas9_RNP_Xrcc1_24hrs/data_filtered_10M/",
  samples = summaries_new_filt[["BJ_Cas9_RNP_Xrcc1_24hrs"]]$sample,
  num_threads = 10)

bp_xrcc1_24hrs_filt_gr <- unlist(bp_xrcc1_24hrs_filt_grl)
names(bp_xrcc1_24hrs_filt_gr) <- NULL
mcols(bp_xrcc1_24hrs_filt_gr)$sample <- gsub("\\.hg19\\.filt\\.bam\\.RData_filtered|\\.hg19\\.filt\\.bam\\.RData", "", bp_xrcc1_24hrs_filt_gr$file)

bp_xrcc1_24hrs_filt_rescue_gr <- add_rescue_breaks(bp_xrcc1_24hrs_filt_gr, xrcc1_new_24hrs, xrcc1_coord_str)
bp_xrcc1_24hrs_filt_rescue_grl <- split(bp_xrcc1_24hrs_filt_rescue_gr, bp_xrcc1_24hrs_filt_rescue_gr$sample)

# Zoom chr19
bp_zoom <- bp_xrcc1_24hrs_filt_rescue_gr
seqlevels(bp_zoom, pruning.mode = "coarse") <- seqlevels(seqinfo(bp_zoom)["19"])
seqinfo(bp_zoom) <- seqinfo(bp_zoom)["19"]

pileup_zoom_xrcc1_24hrs <- plot_aggregate_breaks(GRangesList(bp_zoom),
  tilewidth = 1000000, cent = hg19_cent, hotspot_pval = 1e-08,
  return_individual_plots = TRUE, normalize_counts = TRUE,
  n = length(unique(bp_zoom$sample)), regions = rnp_xrcc1)

chr_max <- max(pileup_zoom_xrcc1_24hrs[[1]]$layers[[2]]$data$end.genome)
p_zoom <- (pileup_zoom_xrcc1_24hrs[[1]] + geom_vline(xintercept = chr_max, size = 0.3) + legend_theme) +
  (pileup_zoom_xrcc1_24hrs[[4]] + geom_vline(xintercept = chr_max, size = 0.3) + legend_theme) +
  plot_layout(ncol = 1, heights = c(5, 1))
ggsave("pileup_zoom_Xrcc1_24hrs.pdf", p_zoom, grDevices::cairo_pdf, fig_dir, height = 2, width = 2.1)
message("Saved pileup_zoom_Xrcc1_24hrs.pdf")

# ===================================================================
# 2. BJ_Cas9_RNP_Xrcc1_24wsBrdU
# ===================================================================
message("--- Xrcc1_24wsBrdU pileup ---")

bp_xrcc1_brdu_filt_grl <- breakpoints_to_list(
  "/u/project/yeastyin/yeastyin/projects/Cas9_RNP/data/bpR_output/BJ_Cas9_RNP_Xrcc1_24wsBrdU/data_filtered_10M/",
  samples = summaries_new_filt[["BJ_Cas9_RNP_Xrcc1_24wsBrdU"]]$sample,
  num_threads = 10)

bp_xrcc1_brdu_filt_gr <- unlist(bp_xrcc1_brdu_filt_grl)
names(bp_xrcc1_brdu_filt_gr) <- NULL
mcols(bp_xrcc1_brdu_filt_gr)$sample <- gsub("\\.hg19\\.filt\\.bam\\.RData_filtered|\\.hg19\\.filt\\.bam\\.RData", "", bp_xrcc1_brdu_filt_gr$file)

bp_xrcc1_brdu_filt_rescue_gr <- add_rescue_breaks(bp_xrcc1_brdu_filt_gr, xrcc1_new_24wsBrdU, xrcc1_coord_str)
bp_xrcc1_brdu_filt_rescue_grl <- split(bp_xrcc1_brdu_filt_rescue_gr, bp_xrcc1_brdu_filt_rescue_gr$sample)

# Zoom chr19
bp_zoom <- bp_xrcc1_brdu_filt_rescue_gr
seqlevels(bp_zoom, pruning.mode = "coarse") <- seqlevels(seqinfo(bp_zoom)["19"])
seqinfo(bp_zoom) <- seqinfo(bp_zoom)["19"]

pileup_zoom_xrcc1_brdu <- plot_aggregate_breaks(GRangesList(bp_zoom),
  tilewidth = 1000000, cent = hg19_cent, hotspot_pval = 1e-08,
  return_individual_plots = TRUE, normalize_counts = TRUE,
  n = length(unique(bp_zoom$sample)), regions = rnp_xrcc1)

chr_max <- max(pileup_zoom_xrcc1_brdu[[1]]$layers[[2]]$data$end.genome)
p_zoom <- (pileup_zoom_xrcc1_brdu[[1]] + geom_vline(xintercept = chr_max, size = 0.3) + legend_theme) +
  (pileup_zoom_xrcc1_brdu[[4]] + geom_vline(xintercept = chr_max, size = 0.3) + legend_theme) +
  plot_layout(ncol = 1, heights = c(5, 1))
ggsave("pileup_zoom_Xrcc1_24wsBrdU.pdf", p_zoom, grDevices::cairo_pdf, fig_dir, height = 2, width = 2.1)
message("Saved pileup_zoom_Xrcc1_24wsBrdU.pdf")

message("Done: 04b_xrcc1_new_pileup.R")
```
