## Supplementary material for "Genome-wide mapping of Cas9-induced sister chromatid exchange across single and over 200 genomic targets": Cas9_RNP_scripts.zip: 05_lig3_rescue.R

```
#!/usr/bin/env Rscript
# 05_lig3_rescue.R
# Rescue breakpoint analysis at LIG3 locus (chr17:33307531-33336762)
# Datasets with LIG3 cut: p20250808, p20250409_lig3, p20250924_lig3, HAP1,
#   + new Lig3_24hrs, Lig3_24wsBrdU

message("=== 05_lig3_rescue.R ===")
source("scripts/load_env.R")
load("data/summaries_earlier.RData")
load("data/summaries_new10.RData")

lig3_coord_str <- "17:33307531-33336762"

# ===================================================================
# 1. p20250808: xrcc1+lig3+237 (has LIG3 cut)
# ===================================================================
message("--- p20250808 LIG3 rescue ---")

lig3_p20250808 <- run_check_location_for_breakpoints(
  c("/u/project/yeastyin/chovanec/projects/novaseq250815/bj_rnp_xrcc1_lig3_seq3_237/human/breakpointR_diploid/data_filtered/",
    "/u/project/yeastyin/chovanec/projects/novaseq250919/bj_rnp_xrcc1_lig3_seq3_237/human/breakpointR_diploid/data_filtered/"),
  coord = lig3_coord_str, samples = summary_p20250808_filt$sample,
  num_threads = 10, bin_width = 10000000, dist_from_gr = 3000000, type = "diploid")

message("p20250808 LIG3 rescue rate: ", sum(lig3_p20250808$rescue_break), "/", nrow(lig3_p20250808))

# Pileup with rescue
bp_p20250808_filt_grl <- breakpoints_to_list(
  c("/u/project/yeastyin/chovanec/projects/novaseq250815/bj_rnp_xrcc1_lig3_seq3_237/human/breakpointR_diploid/data_filtered/",
    "/u/project/yeastyin/chovanec/projects/novaseq250919/bj_rnp_xrcc1_lig3_seq3_237/human/breakpointR_diploid/data_filtered/"),
  samples = summary_p20250808_filt$sample, num_threads = 10, filtered = TRUE)

bp_p20250808_filt_gr <- unlist(bp_p20250808_filt_grl)
names(bp_p20250808_filt_gr) <- NULL
mcols(bp_p20250808_filt_gr)$sample <- gsub("\\.human\\.filt\\.bam\\.RData_filtered|\\.hg19\\.filt\\.bam\\.RData_filtered", "", bp_p20250808_filt_gr$file)

bp_p20250808_filt_rescue_gr <- add_rescue_breaks(bp_p20250808_filt_gr, lig3_p20250808, lig3_coord_str)
bp_p20250808_filt_rescue_grl <- split(bp_p20250808_filt_rescue_gr, bp_p20250808_filt_rescue_gr$sample)

hg19_cent <- get_centromere_coordinates("hg19", ensembl = TRUE)
rnp_lig3 <- GRanges(seqnames = 17, ranges = IRanges(start = 33307531, end = 33336762))
mcols(rnp_lig3)$name <- "LIG3"

# Zoom chr17
bp_zoom <- bp_p20250808_filt_rescue_gr
seqlevels(bp_zoom, pruning.mode = "coarse") <- seqlevels(seqinfo(bp_zoom)["17"])
seqinfo(bp_zoom) <- seqinfo(bp_zoom)["17"]

pileup_zoom_p20250808_lig3 <- plot_aggregate_breaks(GRangesList(bp_zoom),
  tilewidth = 1000000, cent = hg19_cent, hotspot_pval = 1e-08,
  return_individual_plots = TRUE, normalize_counts = TRUE,
  n = length(unique(bp_zoom$sample)), regions = rnp_lig3)

chr_max <- max(pileup_zoom_p20250808_lig3[[1]]$layers[[2]]$data$end.genome)
p_zoom <- (pileup_zoom_p20250808_lig3[[1]] + geom_vline(xintercept = chr_max, size = 0.3) + legend_theme) +
  (pileup_zoom_p20250808_lig3[[4]] + geom_vline(xintercept = chr_max, size = 0.3) + legend_theme) +
  plot_layout(ncol = 1, heights = c(5, 1))
ggsave("pileup_zoom_p20250808_lig3.pdf", p_zoom, grDevices::cairo_pdf, fig_dir, height = 2, width = 2.1)
message("Saved pileup_zoom_p20250808_lig3.pdf")

# ===================================================================
# 2. p20250409: BJ_rnp_lig3 (yi442/443 via novaseq250806)
# ===================================================================
message("--- p20250409 LIG3 rescue ---")

lig3_p20250409 <- run_check_location_for_breakpoints(
  "/u/project/yeastyin/chovanec/projects/novaseq250806/BJ5ta_RNP_XRCC1_LIG3/hg19/breakpointR_diploid/data_filtered/",
  coord = lig3_coord_str,
  samples = novaseq250806_summary_bj_rnp_filt$sample[novaseq250806_summary_bj_rnp_filt$condition == "BJ_rnp_lig3"],
  num_threads = 10, bin_width = 10000000, dist_from_gr = 3000000, type = "diploid")

message("p20250409 LIG3 rescue rate: ", sum(lig3_p20250409$rescue_break), "/", nrow(lig3_p20250409))

# Pileup
bp_lig3_409_filt_grl <- breakpoints_to_list(
  "/u/project/yeastyin/chovanec/projects/novaseq250806/BJ5ta_RNP_XRCC1_LIG3/hg19/breakpointR_diploid/data_filtered/",
  samples = novaseq250806_summary_bj_rnp_filt %>% filter(condition == "BJ_rnp_lig3") %>% pull(sample),
  num_threads = 5, filtered = TRUE)

bp_lig3_409_filt_gr <- unlist(bp_lig3_409_filt_grl)
names(bp_lig3_409_filt_gr) <- NULL
mcols(bp_lig3_409_filt_gr)$sample <- gsub("\\.hg19\\.filt\\.bam\\.RData_filtered", "", bp_lig3_409_filt_gr$file)

bp_lig3_409_filt_rescue_gr <- add_rescue_breaks(bp_lig3_409_filt_gr, lig3_p20250409, lig3_coord_str)
bp_lig3_409_filt_rescue_grl <- split(bp_lig3_409_filt_rescue_gr, bp_lig3_409_filt_rescue_gr$sample)

# Zoom chr17
bp_zoom <- bp_lig3_409_filt_gr
seqlevels(bp_zoom, pruning.mode = "coarse") <- seqlevels(seqinfo(bp_zoom)["17"])
seqinfo(bp_zoom) <- seqinfo(bp_zoom)["17"]

pileup_zoom_p20250409_lig3 <- plot_aggregate_breaks(GRangesList(bp_zoom),
  tilewidth = 1000000, cent = hg19_cent, hotspot_pval = 1e-08,
  return_individual_plots = TRUE, normalize_counts = TRUE,
  n = length(bp_lig3_409_filt_rescue_grl), regions = rnp_lig3)

chr_max <- max(pileup_zoom_p20250409_lig3[[1]]$layers[[2]]$data$end.genome)
p_zoom <- (pileup_zoom_p20250409_lig3[[1]] + geom_vline(xintercept = chr_max, size = 0.3) + legend_theme) +
  (pileup_zoom_p20250409_lig3[[4]] + geom_vline(xintercept = chr_max, size = 0.3) + legend_theme) +
  plot_layout(ncol = 1, heights = c(5, 1))
ggsave("pileup_zoom_p20250409_lig3.pdf", p_zoom, grDevices::cairo_pdf, fig_dir, height = 2, width = 2.1)
message("Saved pileup_zoom_p20250409_lig3.pdf")

# ===================================================================
# 3. p20250924: bj_rnp_lig3 (yi479)
# ===================================================================
message("--- p20250924 LIG3 rescue ---")

lig3_p20250924 <- run_check_location_for_breakpoints(
  "/u/project/yeastyin/chovanec/projects/novaseq251007/BJ_RNP/human/breakpointR_diploid/data_filtered/",
  coord = lig3_coord_str,
  samples = summary_p20250924_filt %>% filter(condition == "bj_rnp_lig3") %>% pull(sample),
  num_threads = 10, bin_width = 10000000, dist_from_gr = 3000000, type = "diploid")

message("p20250924 LIG3 rescue rate: ", sum(lig3_p20250924$rescue_break), "/", nrow(lig3_p20250924))

# Pileup
bp_lig3_924_filt_grl <- breakpoints_to_list(
  "/u/project/yeastyin/chovanec/projects/novaseq251007/BJ_RNP/human/breakpointR_diploid/data_filtered/",
  samples = summary_p20250924_filt %>% filter(condition == "bj_rnp_lig3") %>% pull(sample),
  num_threads = 5, filtered = TRUE)

bp_lig3_924_filt_gr <- unlist(bp_lig3_924_filt_grl)
names(bp_lig3_924_filt_gr) <- NULL
mcols(bp_lig3_924_filt_gr)$sample <- gsub("\\.hg19\\.filt\\.bam\\.RData_filtered", "", bp_lig3_924_filt_gr$file)

bp_lig3_924_filt_rescue_gr <- add_rescue_breaks(bp_lig3_924_filt_gr, lig3_p20250924, lig3_coord_str)
bp_lig3_924_filt_rescue_grl <- split(bp_lig3_924_filt_rescue_gr, bp_lig3_924_filt_rescue_gr$sample)

# Zoom chr17
bp_zoom <- bp_lig3_924_filt_gr
seqlevels(bp_zoom, pruning.mode = "coarse") <- seqlevels(seqinfo(bp_zoom)["17"])
seqinfo(bp_zoom) <- seqinfo(bp_zoom)["17"]

pileup_zoom_p20250924_lig3 <- plot_aggregate_breaks(GRangesList(bp_zoom),
  tilewidth = 1000000, cent = hg19_cent, hotspot_pval = 1e-08,
  return_individual_plots = TRUE, normalize_counts = TRUE,
  n = length(bp_lig3_924_filt_rescue_grl), regions = rnp_lig3)

chr_max <- max(pileup_zoom_p20250924_lig3[[1]]$layers[[2]]$data$end.genome)
p_zoom <- (pileup_zoom_p20250924_lig3[[1]] + geom_vline(xintercept = chr_max, size = 0.3) + legend_theme) +
  (pileup_zoom_p20250924_lig3[[4]] + geom_vline(xintercept = chr_max, size = 0.3) + legend_theme) +
  plot_layout(ncol = 1, heights = c(5, 1))
ggsave("pileup_zoom_p20250924_lig3.pdf", p_zoom, grDevices::cairo_pdf, fig_dir, height = 2, width = 2.1)
message("Saved pileup_zoom_p20250924_lig3.pdf")

# ===================================================================
# 4. HAP1 haploid
# ===================================================================
message("--- HAP1 haploid LIG3 rescue ---")

lig3_hap1_haploid <- run_check_location_for_breakpoints(
  "/u/project/yeastyin/chovanec/projects/novaseq250806/HAP1_RNP_XRCC1_LIG3/hg19/breakpointR_haploid/data_filtered/",
  coord = lig3_coord_str, samples = summary_hap1_haploid_filt$sample,
  num_threads = 10, bin_width = 10000000, dist_from_gr = 3000000, type = "haploid")

message("HAP1 haploid LIG3 rescue rate: ", sum(lig3_hap1_haploid$rescue_break), "/", nrow(lig3_hap1_haploid))

# ===================================================================
# 5. HAP1 diploid
# ===================================================================
message("--- HAP1 diploid LIG3 rescue ---")

lig3_hap1_diploid <- run_check_location_for_breakpoints(
  "/u/project/yeastyin/chovanec/projects/novaseq250806/HAP1_RNP_XRCC1_LIG3/hg19/breakpointR_diploid/data_filtered/",
  coord = lig3_coord_str, samples = summary_hap1_diploid_filt$sample,
  num_threads = 10, bin_width = 10000000, dist_from_gr = 3000000, type = "diploid")

message("HAP1 diploid LIG3 rescue rate: ", sum(lig3_hap1_diploid$rescue_break), "/", nrow(lig3_hap1_diploid))

# ===================================================================
# 6. New: BJ_Cas9_RNP_Lig3_24hrs
# ===================================================================
message("--- New Lig3_24hrs rescue ---")

lig3_new_24hrs <- run_check_location_for_breakpoints(
  "/u/project/yeastyin/yeastyin/projects/Cas9_RNP/data/bpR_output/BJ_Cas9_RNP_Lig3_24hrs/data_filtered_10M/",
  coord = lig3_coord_str,
  samples = summaries_new_filt[["BJ_Cas9_RNP_Lig3_24hrs"]]$sample,
  num_threads = 10, bin_width = 10000000, dist_from_gr = 3000000, type = "diploid")

message("Lig3_24hrs rescue rate: ", sum(lig3_new_24hrs$rescue_break), "/", nrow(lig3_new_24hrs))

# ===================================================================
# 7. New: BJ_Cas9_RNP_Lig3_24wsBrdU
# ===================================================================
message("--- New Lig3_24wsBrdU rescue ---")

lig3_new_24wsBrdU <- run_check_location_for_breakpoints(
  "/u/project/yeastyin/yeastyin/projects/Cas9_RNP/data/bpR_output/BJ_Cas9_RNP_Lig3_24wsBrdU/data_filtered_10M/",
  coord = lig3_coord_str,
  samples = summaries_new_filt[["BJ_Cas9_RNP_Lig3_24wsBrdU"]]$sample,
  num_threads = 10, bin_width = 10000000, dist_from_gr = 3000000, type = "diploid")

message("Lig3_24wsBrdU rescue rate: ", sum(lig3_new_24wsBrdU$rescue_break), "/", nrow(lig3_new_24wsBrdU))

# ===================================================================
# Summary rescue table for LIG3
# ===================================================================
lig3_rescue_summary <- data.frame(
  condition = c("p20250808_xrcc1_lig3_237", "p20250409_lig3",
                "p20250924_lig3", "HAP1_haploid", "HAP1_diploid",
                "Lig3_24hrs", "Lig3_24wsBrdU"),
  total_cells = c(nrow(lig3_p20250808), nrow(lig3_p20250409),
                  nrow(lig3_p20250924), nrow(lig3_hap1_haploid), nrow(lig3_hap1_diploid),
                  nrow(lig3_new_24hrs), nrow(lig3_new_24wsBrdU)),
  rescued = c(sum(lig3_p20250808$rescue_break), sum(lig3_p20250409$rescue_break),
              sum(lig3_p20250924$rescue_break), sum(lig3_hap1_haploid$rescue_break),
              sum(lig3_hap1_diploid$rescue_break),
              sum(lig3_new_24hrs$rescue_break), sum(lig3_new_24wsBrdU$rescue_break))
)
lig3_rescue_summary$rescue_rate <- round(lig3_rescue_summary$rescued / lig3_rescue_summary$total_cells, 4)

write.table(lig3_rescue_summary, file.path(tab_dir, "lig3_rescue_summary.txt"),
            sep = "\t", row.names = FALSE, quote = FALSE)
message("Saved lig3_rescue_summary.txt")
print(lig3_rescue_summary)

# ---- Save ----
save(lig3_p20250808, lig3_p20250409, lig3_p20250924,
     lig3_hap1_haploid, lig3_hap1_diploid,
     lig3_new_24hrs, lig3_new_24wsBrdU,
     lig3_rescue_summary,
     bp_p20250808_filt_rescue_grl,
     bp_lig3_409_filt_rescue_grl,
     bp_lig3_924_filt_rescue_grl,
     file = "data/lig3_rescue.RData")

message("Done: data/lig3_rescue.RData")
```
