## Supplementary material for "Genome-wide mapping of Cas9-induced sister chromatid exchange across single and over 200 genomic targets": Cas9_RNP_scripts.zip: 07_xrcc4_rescue.R

```
#!/usr/bin/env Rscript
# 07_xrcc4_rescue.R
# Rescue breakpoint analysis at XRCC4 locus (chr5:82367495-82643756)
# NEW locus - only new conditions: Xrcc4_24hrs, Xrcc4_24wsBrdU

message("=== 07_xrcc4_rescue.R ===")
source("scripts/load_env.R")
load("data/summaries_new10.RData")

xrcc4_coord_str <- "5:82367495-82643756"

# ===================================================================
# 1. New: BJ_Cas9_RNP_Xrcc4_24hrs
# ===================================================================
message("--- Xrcc4_24hrs rescue ---")

xrcc4_new_24hrs <- run_check_location_for_breakpoints(
  "/u/project/yeastyin/yeastyin/projects/Cas9_RNP/data/bpR_output/BJ_Cas9_RNP_Xrcc4_24hrs/data_filtered_10M/",
  coord = xrcc4_coord_str,
  samples = summaries_new_filt[["BJ_Cas9_RNP_Xrcc4_24hrs"]]$sample,
  num_threads = 10, bin_width = 10000000, dist_from_gr = 3000000, type = "diploid")

message("Xrcc4_24hrs rescue rate: ", sum(xrcc4_new_24hrs$rescue_break), "/", nrow(xrcc4_new_24hrs))

# Pileup with rescue
bp_xrcc4_24hrs_filt_grl <- breakpoints_to_list(
  "/u/project/yeastyin/yeastyin/projects/Cas9_RNP/data/bpR_output/BJ_Cas9_RNP_Xrcc4_24hrs/data_filtered_10M/",
  samples = summaries_new_filt[["BJ_Cas9_RNP_Xrcc4_24hrs"]]$sample,
  num_threads = 5, filtered = TRUE)

bp_xrcc4_24hrs_filt_gr <- unlist(bp_xrcc4_24hrs_filt_grl)
names(bp_xrcc4_24hrs_filt_gr) <- NULL
mcols(bp_xrcc4_24hrs_filt_gr)$sample <- gsub("\\.hg19\\.filt\\.bam\\.RData_filtered|\\.hg19\\.filt\\.bam\\.RData", "", bp_xrcc4_24hrs_filt_gr$file)

bp_xrcc4_24hrs_filt_rescue_gr <- add_rescue_breaks(bp_xrcc4_24hrs_filt_gr, xrcc4_new_24hrs, xrcc4_coord_str)
bp_xrcc4_24hrs_filt_rescue_grl <- split(bp_xrcc4_24hrs_filt_rescue_gr, bp_xrcc4_24hrs_filt_rescue_gr$sample)

hg19_cent <- get_centromere_coordinates("hg19", ensembl = TRUE)
rnp_xrcc4 <- GRanges(seqnames = 5, ranges = IRanges(start = 82367495, end = 82643756))
mcols(rnp_xrcc4)$name <- "XRCC4"

# Zoom chr5
bp_zoom <- bp_xrcc4_24hrs_filt_rescue_gr
seqlevels(bp_zoom, pruning.mode = "coarse") <- seqlevels(seqinfo(bp_zoom)["5"])
seqinfo(bp_zoom) <- seqinfo(bp_zoom)["5"]

pileup_zoom_xrcc4_24hrs <- plot_aggregate_breaks(GRangesList(bp_zoom),
  tilewidth = 1000000, cent = hg19_cent, hotspot_pval = 1e-08,
  return_individual_plots = TRUE, normalize_counts = TRUE,
  n = length(unique(bp_zoom$sample)), regions = rnp_xrcc4)

chr_max <- max(pileup_zoom_xrcc4_24hrs[[1]]$layers[[2]]$data$end.genome)
p_zoom <- (pileup_zoom_xrcc4_24hrs[[1]] + geom_vline(xintercept = chr_max, size = 0.3) + legend_theme) +
  (pileup_zoom_xrcc4_24hrs[[4]] + geom_vline(xintercept = chr_max, size = 0.3) + legend_theme) +
  plot_layout(ncol = 1, heights = c(5, 1))
ggsave("pileup_zoom_Xrcc4_24hrs.pdf", p_zoom, grDevices::cairo_pdf, fig_dir, height = 2, width = 2.1)
message("Saved pileup_zoom_Xrcc4_24hrs.pdf")

# ===================================================================
# 2. New: BJ_Cas9_RNP_Xrcc4_24wsBrdU
# ===================================================================
message("--- Xrcc4_24wsBrdU rescue ---")

xrcc4_new_24wsBrdU <- run_check_location_for_breakpoints(
  "/u/project/yeastyin/yeastyin/projects/Cas9_RNP/data/bpR_output/BJ_Cas9_RNP_Xrcc4_24wsBrdU/data_filtered_10M/",
  coord = xrcc4_coord_str,
  samples = summaries_new_filt[["BJ_Cas9_RNP_Xrcc4_24wsBrdU"]]$sample,
  num_threads = 10, bin_width = 10000000, dist_from_gr = 3000000, type = "diploid")

message("Xrcc4_24wsBrdU rescue rate: ", sum(xrcc4_new_24wsBrdU$rescue_break), "/", nrow(xrcc4_new_24wsBrdU))

# Pileup with rescue
bp_xrcc4_brdu_filt_grl <- breakpoints_to_list(
  "/u/project/yeastyin/yeastyin/projects/Cas9_RNP/data/bpR_output/BJ_Cas9_RNP_Xrcc4_24wsBrdU/data_filtered_10M/",
  samples = summaries_new_filt[["BJ_Cas9_RNP_Xrcc4_24wsBrdU"]]$sample,
  num_threads = 5, filtered = TRUE)

bp_xrcc4_brdu_filt_gr <- unlist(bp_xrcc4_brdu_filt_grl)
names(bp_xrcc4_brdu_filt_gr) <- NULL
mcols(bp_xrcc4_brdu_filt_gr)$sample <- gsub("\\.hg19\\.filt\\.bam\\.RData_filtered|\\.hg19\\.filt\\.bam\\.RData", "", bp_xrcc4_brdu_filt_gr$file)

bp_xrcc4_brdu_filt_rescue_gr <- add_rescue_breaks(bp_xrcc4_brdu_filt_gr, xrcc4_new_24wsBrdU, xrcc4_coord_str)
bp_xrcc4_brdu_filt_rescue_grl <- split(bp_xrcc4_brdu_filt_rescue_gr, bp_xrcc4_brdu_filt_rescue_gr$sample)

# Zoom chr5
bp_zoom <- bp_xrcc4_brdu_filt_rescue_gr
seqlevels(bp_zoom, pruning.mode = "coarse") <- seqlevels(seqinfo(bp_zoom)["5"])
seqinfo(bp_zoom) <- seqinfo(bp_zoom)["5"]

pileup_zoom_xrcc4_brdu <- plot_aggregate_breaks(GRangesList(bp_zoom),
  tilewidth = 1000000, cent = hg19_cent, hotspot_pval = 1e-08,
  return_individual_plots = TRUE, normalize_counts = TRUE,
  n = length(unique(bp_zoom$sample)), regions = rnp_xrcc4)

chr_max <- max(pileup_zoom_xrcc4_brdu[[1]]$layers[[2]]$data$end.genome)
p_zoom <- (pileup_zoom_xrcc4_brdu[[1]] + geom_vline(xintercept = chr_max, size = 0.3) + legend_theme) +
  (pileup_zoom_xrcc4_brdu[[4]] + geom_vline(xintercept = chr_max, size = 0.3) + legend_theme) +
  plot_layout(ncol = 1, heights = c(5, 1))
ggsave("pileup_zoom_Xrcc4_24wsBrdU.pdf", p_zoom, grDevices::cairo_pdf, fig_dir, height = 2, width = 2.1)
message("Saved pileup_zoom_Xrcc4_24wsBrdU.pdf")

# ===================================================================
# Summary rescue table for XRCC4
# ===================================================================
xrcc4_rescue_summary <- data.frame(
  condition = c("Xrcc4_24hrs", "Xrcc4_24wsBrdU"),
  total_cells = c(nrow(xrcc4_new_24hrs), nrow(xrcc4_new_24wsBrdU)),
  rescued = c(sum(xrcc4_new_24hrs$rescue_break), sum(xrcc4_new_24wsBrdU$rescue_break))
)
xrcc4_rescue_summary$rescue_rate <- round(xrcc4_rescue_summary$rescued / xrcc4_rescue_summary$total_cells, 4)

write.table(xrcc4_rescue_summary, file.path(tab_dir, "xrcc4_rescue_summary.txt"),
            sep = "\t", row.names = FALSE, quote = FALSE)
message("Saved xrcc4_rescue_summary.txt")
print(xrcc4_rescue_summary)

# ---- Save ----
save(xrcc4_new_24hrs, xrcc4_new_24wsBrdU, xrcc4_rescue_summary,
     bp_xrcc4_24hrs_filt_rescue_grl, bp_xrcc4_brdu_filt_rescue_grl,
     file = "data/xrcc4_rescue.RData")

message("Done: data/xrcc4_rescue.RData")
```
