## Supplementary material for "Genome-wide mapping of Cas9-induced sister chromatid exchange across single and over 200 genomic targets": Cas9_RNP_scripts.zip: 08_237_pileup.R

```
#!/usr/bin/env Rscript
# 08_237_pileup.R
# 237 guide pileup analysis: WT, d10a nickase, h840a nickase
# Datasets: p20250808, p20250924 (WT/d10a/h840a), new 237cuts_24hrs, 237cuts_24ws_BrdU

message("=== 08_237_pileup.R ===")
source("scripts/load_env.R")
load("data/summaries_earlier.RData")
load("data/summaries_new10.RData")

# ---- Load 237 guide coordinates ----
message("Loading 237 guide coordinates from BAM...")
yy_237_gr <- bam2granges("/u/project/yeastyin/chovanec/projects/RNP_workup/YY_SEQ3_237_GUIDE1.srt.bam", paired_end = FALSE)
names(yy_237_gr) <- NULL

# Remove hs37d5 and chrY
seqlevels(yy_237_gr, pruning.mode = "coarse") <- standardChromosomes(yy_237_gr)[1:23]

# Load BJ-5ta inversions and blacklist for breakpoint filtering (panels 3-4)
bj_region_filt <- rtracklayer::import.bed("/u/project/yeastyin/chovanec/software/scistrandfig/inst/extdata/regions/BJ-5ta_inv_bl.bed")
hg19_blk <- rtracklayer::import.bed("/u/project/yeastyin/yeastyin/pipelines/sciL3StrandPipe/resources/blacklist/hg19-blacklist.v2.ensembl.bed")

hg19_cent <- get_centromere_coordinates("hg19", ensembl = TRUE)

# ===================================================================
# Helper: run pileup for a 237 condition
# ===================================================================
run_237_pileup <- function(data_folder, samples, condition_name, use_filtered = TRUE) {
  message("--- ", condition_name, " pileup ---")

  bp_grl <- breakpoints_to_list(data_folder, samples = samples, num_threads = 5, filtered = FALSE)
  bp_filt_grl <- breakpoints_to_list(data_folder, samples = samples, num_threads = 5, filtered = use_filtered)

  bp_gr <- unlist(bp_grl)
  names(bp_gr) <- NULL
  mcols(bp_gr)$sample <- gsub("\\.human\\.filt\\.bam\\.RData_filtered|\\.hg19\\.filt\\.bam\\.RData_filtered|\\.hg19\\.filt\\.bam\\.RData", "", bp_gr$file)
  bp_grl <- split(bp_gr, bp_gr$sample)

  bp_filt_gr <- unlist(bp_filt_grl)
  names(bp_filt_gr) <- NULL
  mcols(bp_filt_gr)$sample <- gsub("\\.human\\.filt\\.bam\\.RData_filtered|\\.hg19\\.filt\\.bam\\.RData_filtered|\\.hg19\\.filt\\.bam\\.RData", "", bp_filt_gr$file)
  # Filter breakpoints against BJ-5ta inversions and blacklist (panels 3-4)
  bp_filt_gr <- plyranges::filter_by_non_overlaps(bp_filt_gr, c(bj_region_filt, hg19_blk))
  bp_filt_grl <- split(bp_filt_gr, bp_filt_gr$sample)

  # Normalized pileup
  pileup_norm <- plot_aggregate_breaks(bp_grl,
    tilewidth = 1000000, cent = hg19_cent, hotspot_pval = NULL,
    return_individual_plots = TRUE, normalize_counts = TRUE,
    n = length(bp_grl))

  pileup_raw <- plot_aggregate_breaks(bp_grl,
    tilewidth = 1000000, cent = hg19_cent, hotspot_pval = NULL,
    return_individual_plots = TRUE, normalize_counts = FALSE,
    n = length(bp_grl))

  pileup_filt_norm <- plot_aggregate_breaks(bp_filt_grl,
    tilewidth = 1000000, cent = hg19_cent, hotspot_pval = NULL,
    return_individual_plots = TRUE, normalize_counts = TRUE,
    n = length(bp_filt_grl))

  pileup_filt_raw <- plot_aggregate_breaks(bp_filt_grl,
    tilewidth = 1000000, cent = hg19_cent, hotspot_pval = NULL,
    return_individual_plots = TRUE, normalize_counts = FALSE,
    n = length(bp_filt_grl), regions = yy_237_gr)

  # Combined figure
  p <- pileup_norm[[1]] + ggtitle(paste(condition_name, "- all breaks (normalized)")) +
    pileup_raw[[1]] + ggtitle("all breaks (raw)") +
    pileup_filt_norm[[1]] + ggtitle("filtered breaks (normalized)") +
    pileup_filt_raw[[1]] + ggtitle("filtered breaks (raw)") +
    pileup_filt_raw[[4]] +
    plot_layout(nrow = 5, heights = c(4, 4, 4, 4, 1))

  fname <- paste0("pileup_237_", gsub(" ", "_", condition_name), ".pdf")
  ggsave(fname, p, grDevices::cairo_pdf, fig_dir, height = 4.5, width = 12)
  message("Saved ", fname)

  return(list(bp_grl = bp_grl, bp_filt_grl = bp_filt_grl,
              bp_filt_gr = bp_filt_gr, n_cells = length(bp_grl)))
}

# ===================================================================
# p20250808: WT 237 (has 237+XRCC1+LIG3 cuts)
# ===================================================================
p20250808_237 <- run_237_pileup(
  c("/u/project/yeastyin/chovanec/projects/novaseq250815/bj_rnp_xrcc1_lig3_seq3_237/human/breakpointR_diploid/data_filtered/",
    "/u/project/yeastyin/chovanec/projects/novaseq250919/bj_rnp_xrcc1_lig3_seq3_237/human/breakpointR_diploid/data_filtered/"),
  samples = summary_p20250808_filt$sample,
  condition_name = "p20250808_WT237")

# ===================================================================
# p20250924: WT 237
# ===================================================================
p20250924_237_wt <- run_237_pileup(
  "/u/project/yeastyin/chovanec/projects/novaseq251007/BJ_RNP/human/breakpointR_diploid/data_filtered/",
  samples = summary_p20250924_filt %>% filter(condition == "bj_rnp_seq3_237") %>% pull(sample),
  condition_name = "p20250924_WT237")

# ===================================================================
# p20250924: d10a nickase
# ===================================================================
p20250924_237_d10a <- run_237_pileup(
  "/u/project/yeastyin/chovanec/projects/novaseq251007/BJ_RNP/human/breakpointR_diploid/data_filtered/",
  samples = summary_p20250924_filt %>% filter(condition == "bj_rnp_seq3_237_d10a") %>% pull(sample),
  condition_name = "p20250924_d10a")

# ===================================================================
# p20250924: h840a nickase
# ===================================================================
p20250924_237_h840a <- run_237_pileup(
  "/u/project/yeastyin/chovanec/projects/novaseq251007/BJ_RNP/human/breakpointR_diploid/data_filtered/",
  samples = summary_p20250924_filt %>% filter(condition == "bj_rnp_seq3_237_h840a") %>% pull(sample),
  condition_name = "p20250924_h840a")

# ===================================================================
# New: 237cuts_24hrs_16hrs
# ===================================================================
new_237_24hrs <- run_237_pileup(
  "/u/project/yeastyin/yeastyin/projects/Cas9_RNP/data/bpR_output/BJ_Cas9_RNP_237cuts_24hrs_16hrs/data_filtered_10M/",
  samples = summaries_new_filt[["BJ_Cas9_RNP_237cuts_24hrs_16hrs"]]$sample,
  condition_name = "237cuts_24hrs", use_filtered = TRUE)

# ===================================================================
# New: 237cuts_24ws_40hrs_BrdU
# ===================================================================
new_237_brdu <- run_237_pileup(
  "/u/project/yeastyin/yeastyin/projects/Cas9_RNP/data/bpR_output/BJ_Cas9_RNP_237cuts_24ws_40hrs_BrdU/data_filtered_10M/",
  samples = summaries_new_filt[["BJ_Cas9_RNP_237cuts_24ws_40hrs_BrdU"]]$sample,
  condition_name = "237cuts_24ws_BrdU", use_filtered = TRUE)

# ---- Save ----
save(yy_237_gr,
     p20250808_237, p20250924_237_wt, p20250924_237_d10a, p20250924_237_h840a,
     new_237_24hrs, new_237_brdu,
     file = "data/pileup_237.RData")

message("Done: data/pileup_237.RData")
```
