## Supplementary material for "Genome-wide mapping of Cas9-induced sister chromatid exchange across single and over 200 genomic targets": Cas9_RNP_scripts.zip: 08b_237_pileup_update_yi564.R

```
#!/usr/bin/env Rscript
# 08b_237_pileup_update_yi564.R
# Re-run 237 pileup for ONLY the 2 new conditions (24hrs, BrdU) that got yi564 cells.
# Preserves existing pileup results for old conditions (WT, d10a, h840a).

message("=== 08b_237_pileup_update_yi564.R ===")
source("scripts/load_env.R")
load("data/summaries_earlier.RData")
load("data/summaries_new10.RData")

# Load existing pileup (contains old condition results we want to keep)
load("data/pileup_237.RData")
message("Loaded existing pileup_237.RData — keeping old conditions intact")
message("  p20250808_237: ", p20250808_237$n_cells, " cells")
message("  p20250924_237_wt: ", p20250924_237_wt$n_cells, " cells")
message("  p20250924_237_d10a: ", p20250924_237_d10a$n_cells, " cells")
message("  p20250924_237_h840a: ", p20250924_237_h840a$n_cells, " cells")

# ---- Load resources for breakpoint filtering ----
bj_region_filt <- rtracklayer::import.bed("/u/project/yeastyin/chovanec/software/scistrandfig/inst/extdata/regions/BJ-5ta_inv_bl.bed")
hg19_blk <- rtracklayer::import.bed("/u/project/yeastyin/yeastyin/pipelines/sciL3StrandPipe/resources/blacklist/hg19-blacklist.v2.ensembl.bed")
hg19_cent <- get_centromere_coordinates("hg19", ensembl = TRUE)

# ===================================================================
# Helper: run pileup for a 237 condition (same as 08_237_pileup.R)
# ===================================================================
run_237_pileup <- function(data_folder, samples, condition_name, use_filtered = TRUE) {
  message("--- ", condition_name, " pileup ---")

  bp_grl <- breakpoints_to_list(data_folder, samples = samples, num_threads = 5, filtered = FALSE)
  bp_filt_grl <- breakpoints_to_list(data_folder, samples = samples, num_threads = 5, filtered = use_filtered)

  bp_gr <- unlist(bp_grl)
  names(bp_gr) <- NULL
  mcols(bp_gr)$sample <- gsub("\\.human\\.filt\\.bam\\.RData_filtered|\\.hg19\\.filt\\.bam\\.RData_filtered|\\.hg19\\.filt\\.bam\\.RData", "", bp_gr$file)
  bp_grl <- split(bp_gr, bp_gr$sample)

  bp_filt_gr <- unlist(bp_filt_grl)
  names(bp_filt_gr) <- NULL
  mcols(bp_filt_gr)$sample <- gsub("\\.human\\.filt\\.bam\\.RData_filtered|\\.hg19\\.filt\\.bam\\.RData_filtered|\\.hg19\\.filt\\.bam\\.RData", "", bp_filt_gr$file)
  # Filter breakpoints against BJ-5ta inversions and blacklist
  bp_filt_gr <- plyranges::filter_by_non_overlaps(bp_filt_gr, c(bj_region_filt, hg19_blk))
  bp_filt_grl <- split(bp_filt_gr, bp_filt_gr$sample)

  # Normalized pileup
  pileup_norm <- plot_aggregate_breaks(bp_grl,
    tilewidth = 1000000, cent = hg19_cent, hotspot_pval = NULL,
    return_individual_plots = TRUE, normalize_counts = TRUE,
    n = length(bp_grl))

  pileup_raw <- plot_aggregate_breaks(bp_grl,
    tilewidth = 1000000, cent = hg19_cent, hotspot_pval = NULL,
    return_individual_plots = TRUE, normalize_counts = FALSE,
    n = length(bp_grl))

  pileup_filt_norm <- plot_aggregate_breaks(bp_filt_grl,
    tilewidth = 1000000, cent = hg19_cent, hotspot_pval = NULL,
    return_individual_plots = TRUE, normalize_counts = TRUE,
    n = length(bp_filt_grl))

  pileup_filt_raw <- plot_aggregate_breaks(bp_filt_grl,
    tilewidth = 1000000, cent = hg19_cent, hotspot_pval = NULL,
    return_individual_plots = TRUE, normalize_counts = FALSE,
    n = length(bp_filt_grl), regions = yy_237_gr)

  # Combined figure
  p <- pileup_norm[[1]] + ggtitle(paste(condition_name, "- all breaks (normalized)")) +
    pileup_raw[[1]] + ggtitle("all breaks (raw)") +
    pileup_filt_norm[[1]] + ggtitle("filtered breaks (normalized)") +
    pileup_filt_raw[[1]] + ggtitle("filtered breaks (raw)") +
    pileup_filt_raw[[4]] +
    plot_layout(nrow = 5, heights = c(4, 4, 4, 4, 1))

  fname <- paste0("pileup_237_", gsub(" ", "_", condition_name), ".pdf")
  ggsave(fname, p, grDevices::cairo_pdf, fig_dir, height = 4.5, width = 12)
  message("Saved ", fname)

  return(list(bp_grl = bp_grl, bp_filt_grl = bp_filt_grl,
              bp_filt_gr = bp_filt_gr, n_cells = length(bp_grl)))
}

# ===================================================================
# Re-run ONLY the 2 new conditions with expanded yi564 data
# ===================================================================

message("\n=== Re-running 237cuts_24hrs (now includes yi564 cells) ===")
new_237_24hrs <- run_237_pileup(
  "/u/project/yeastyin/yeastyin/projects/Cas9_RNP/data/bpR_output/BJ_Cas9_RNP_237cuts_24hrs_16hrs/data_filtered_10M/",
  samples = summaries_new_filt[["BJ_Cas9_RNP_237cuts_24hrs_16hrs"]]$sample,
  condition_name = "237cuts_24hrs", use_filtered = TRUE)
message("  237cuts_24hrs: ", new_237_24hrs$n_cells, " cells (was ",
        length(grep("yi564", names(new_237_24hrs$bp_grl), invert=TRUE)), " old + ",
        length(grep("yi564", names(new_237_24hrs$bp_grl))), " yi564)")

message("\n=== Re-running 237cuts_24ws_BrdU (now includes yi564 cells) ===")
new_237_brdu <- run_237_pileup(
  "/u/project/yeastyin/yeastyin/projects/Cas9_RNP/data/bpR_output/BJ_Cas9_RNP_237cuts_24ws_40hrs_BrdU/data_filtered_10M/",
  samples = summaries_new_filt[["BJ_Cas9_RNP_237cuts_24ws_40hrs_BrdU"]]$sample,
  condition_name = "237cuts_24ws_BrdU", use_filtered = TRUE)
message("  237cuts_24ws_BrdU: ", new_237_brdu$n_cells, " cells")

# ---- Save (overwrites with updated new conditions, preserved old) ----
save(yy_237_gr,
     p20250808_237, p20250924_237_wt, p20250924_237_d10a, p20250924_237_h840a,
     new_237_24hrs, new_237_brdu,
     file = "data/pileup_237.RData")

message("\nDone: data/pileup_237.RData updated (old conditions preserved, new conditions re-run)")
```
