## Supplementary material for "Genome-wide mapping of Cas9-induced sister chromatid exchange across single and over 200 genomic targets": Cas9_RNP_scripts.zip: 09_237_enrichment.R

```
#!/usr/bin/env Rscript
# 09_237_enrichment.R
# Contingency table + chi-square enrichment tests for 237 guide datasets
# Tests whether breakpoints are enriched at 237 guide sites vs random regions

message("=== 09_237_enrichment.R ===")
source("scripts/load_env.R")
load("data/summaries_earlier.RData")
load("data/summaries_new10.RData")
load("data/pileup_237.RData")

# ---- Prepare chromosome arms (exclude Y, centromeres) ----
cent_hg19 <- get_centromere_coordinates("hg19", FALSE, ensembl = TRUE)
arms <- gaps(cent_hg19)
arms <- arms[strand(arms) == "*"]
# Exclude Y chr: first 42 elements cover chr1-22 + X (2 arms each = 44 - 2 for chrX short arm = 42?)
arms_filt <- arms[1:42]
seqlevels(arms_filt, pruning.mode = "coarse") <- seqlevels(arms_filt)[1:23]

# ===================================================================
# 1. p20250924 WT 237 - with rescue
# ===================================================================
message("--- Contingency: p20250924 WT237 (with rescue) ---")
c237_wt_resc <- build_contingency_table(
  p20250924_237_wt$bp_filt_grl, yy_237_gr,
  genome = arms_filt, seed = 3,
  breakpoint_folder = "/u/project/yeastyin/chovanec/projects/novaseq251007/BJ_RNP/human/breakpointR_diploid/data_filtered/",
  num_threads = 10)
saveRDS(c237_wt_resc, "data/c237_wt_cont_tab_resc.rds")

message("WT 237 (rescue) chi-sq:")
print(c237_wt_resc$contigency)
ct_wt_resc <- chisq.test(c237_wt_resc$contigency)
message("p-value: ", ct_wt_resc$p.value)

# Without rescue (simple overlap)
message("--- Contingency: p20250924 WT237 (no rescue) ---")
c237_wt_filt <- build_contingency_table(
  p20250924_237_wt$bp_filt_grl, yy_237_gr,
  genome = arms_filt, seed = 3)
ct_wt_filt <- chisq.test(c237_wt_filt$contigency)
message("p-value (no rescue): ", ct_wt_filt$p.value)

# ===================================================================
# 2. p20250924 d10a - with rescue
# ===================================================================
message("--- Contingency: p20250924 d10a (with rescue) ---")
c237_d10a_resc <- build_contingency_table(
  p20250924_237_d10a$bp_filt_grl, yy_237_gr,
  genome = arms_filt, seed = 3,
  breakpoint_folder = "/u/project/yeastyin/chovanec/projects/novaseq251007/BJ_RNP/human/breakpointR_diploid/data_filtered/",
  num_threads = 10)
saveRDS(c237_d10a_resc, "data/c237_d10a_cont_tab_resc.rds")

print(c237_d10a_resc$contigency)
ct_d10a_resc <- chisq.test(c237_d10a_resc$contigency)
message("p-value: ", ct_d10a_resc$p.value)

c237_d10a_filt <- build_contingency_table(
  p20250924_237_d10a$bp_filt_grl, yy_237_gr,
  genome = arms_filt, seed = 3)
ct_d10a_filt <- chisq.test(c237_d10a_filt$contigency)
message("p-value (no rescue): ", ct_d10a_filt$p.value)

# ===================================================================
# 3. p20250924 h840a - with rescue
# ===================================================================
message("--- Contingency: p20250924 h840a (with rescue) ---")
c237_h840a_resc <- build_contingency_table(
  p20250924_237_h840a$bp_filt_grl, yy_237_gr,
  genome = arms_filt, seed = 3,
  breakpoint_folder = "/u/project/yeastyin/chovanec/projects/novaseq251007/BJ_RNP/human/breakpointR_diploid/data_filtered/",
  num_threads = 10)
saveRDS(c237_h840a_resc, "data/c237_h840a_cont_tab_resc.rds")

print(c237_h840a_resc$contigency)
ct_h840a_resc <- chisq.test(c237_h840a_resc$contigency)
message("p-value: ", ct_h840a_resc$p.value)

c237_h840a_filt <- build_contingency_table(
  p20250924_237_h840a$bp_filt_grl, yy_237_gr,
  genome = arms_filt, seed = 3)
ct_h840a_filt <- chisq.test(c237_h840a_filt$contigency)
message("p-value (no rescue): ", ct_h840a_filt$p.value)

# ===================================================================
# 4. New: 237cuts_24hrs
# ===================================================================
message("--- Contingency: 237cuts_24hrs (with rescue) ---")
c237_24hrs_resc <- build_contingency_table(
  new_237_24hrs$bp_filt_grl, yy_237_gr,
  genome = arms_filt, seed = 3,
  breakpoint_folder = "/u/project/yeastyin/yeastyin/projects/Cas9_RNP/data/bpR_output/BJ_Cas9_RNP_237cuts_24hrs_16hrs/data_filtered_10M/",
  num_threads = 10)
saveRDS(c237_24hrs_resc, "data/c237_24hrs_cont_tab_resc.rds")

print(c237_24hrs_resc$contigency)
ct_24hrs_resc <- chisq.test(c237_24hrs_resc$contigency)
message("p-value: ", ct_24hrs_resc$p.value)

c237_24hrs_filt <- build_contingency_table(
  new_237_24hrs$bp_filt_grl, yy_237_gr,
  genome = arms_filt, seed = 3)
ct_24hrs_filt <- chisq.test(c237_24hrs_filt$contigency)
message("p-value (no rescue): ", ct_24hrs_filt$p.value)

# ===================================================================
# 5. New: 237cuts_24ws_BrdU
# ===================================================================
message("--- Contingency: 237cuts_24ws_BrdU (with rescue) ---")
c237_brdu_resc <- build_contingency_table(
  new_237_brdu$bp_filt_grl, yy_237_gr,
  genome = arms_filt, seed = 3,
  breakpoint_folder = "/u/project/yeastyin/yeastyin/projects/Cas9_RNP/data/bpR_output/BJ_Cas9_RNP_237cuts_24ws_40hrs_BrdU/data_filtered_10M/",
  num_threads = 10)
saveRDS(c237_brdu_resc, "data/c237_brdu_cont_tab_resc.rds")

print(c237_brdu_resc$contigency)
ct_brdu_resc <- chisq.test(c237_brdu_resc$contigency)
message("p-value: ", ct_brdu_resc$p.value)

c237_brdu_filt <- build_contingency_table(
  new_237_brdu$bp_filt_grl, yy_237_gr,
  genome = arms_filt, seed = 3)
ct_brdu_filt <- chisq.test(c237_brdu_filt$contigency)
message("p-value (no rescue): ", ct_brdu_filt$p.value)

# ===================================================================
# Summary table
# ===================================================================
enrichment_summary <- data.frame(
  condition = c("p20250924_WT237", "p20250924_d10a", "p20250924_h840a",
                "237cuts_24hrs", "237cuts_24ws_BrdU"),
  n_cells = c(p20250924_237_wt$n_cells, p20250924_237_d10a$n_cells,
              p20250924_237_h840a$n_cells,
              new_237_24hrs$n_cells, new_237_brdu$n_cells),
  rescue_CutSites_present = c(c237_wt_resc$contigency["Cut_sites", "Present"],
                         c237_d10a_resc$contigency["Cut_sites", "Present"],
                         c237_h840a_resc$contigency["Cut_sites", "Present"],
                         c237_24hrs_resc$contigency["Cut_sites", "Present"],
                         c237_brdu_resc$contigency["Cut_sites", "Present"]),
  rescue_Random_present = c(c237_wt_resc$contigency["Random_sites", "Present"],
                            c237_d10a_resc$contigency["Random_sites", "Present"],
                            c237_h840a_resc$contigency["Random_sites", "Present"],
                            c237_24hrs_resc$contigency["Random_sites", "Present"],
                            c237_brdu_resc$contigency["Random_sites", "Present"]),
  rescue_pvalue = c(ct_wt_resc$p.value, ct_d10a_resc$p.value,
                    ct_h840a_resc$p.value,
                    ct_24hrs_resc$p.value, ct_brdu_resc$p.value),
  filt_pvalue = c(ct_wt_filt$p.value, ct_d10a_filt$p.value,
                  ct_h840a_filt$p.value,
                  ct_24hrs_filt$p.value, ct_brdu_filt$p.value)
)

write.table(enrichment_summary, file.path(tab_dir, "enrichment_237_summary.txt"),
            sep = "\t", row.names = FALSE, quote = FALSE)
message("\nEnrichment summary:")
print(enrichment_summary)

# Save individual contingency tables as text
for (name_tab in list(
  list("c237_wt_rescue", c237_wt_resc),
  list("c237_d10a_rescue", c237_d10a_resc),
  list("c237_h840a_rescue", c237_h840a_resc),
  list("c237_24hrs_rescue", c237_24hrs_resc),
  list("c237_brdu_rescue", c237_brdu_resc),
  list("c237_wt_filt", c237_wt_filt),
  list("c237_d10a_filt", c237_d10a_filt),
  list("c237_h840a_filt", c237_h840a_filt),
  list("c237_24hrs_filt", c237_24hrs_filt),
  list("c237_brdu_filt", c237_brdu_filt)
)) {
  ct <- name_tab[[2]]$contigency
  write.table(ct, file.path(tab_dir, paste0("contingency_", name_tab[[1]], ".txt")),
              sep = "\t", quote = FALSE)
}
message("Contingency tables saved to ", tab_dir)

message("Done: 09_237_enrichment.R")
```
