## Supplementary material for "Genome-wide mapping of Cas9-induced sister chromatid exchange across single and over 200 genomic targets": Cas9_RNP_scripts.zip: 09_verify_old_c237.R

```
#!/usr/bin/env Rscript
# 09_verify_old_c237.R
# Verify old condition enrichment tables reproduce HTML reference values
# Run this BEFORE the full 09_237_enrichment.R to check old conditions only

message("=== 09_verify_old_c237.R ===")
source("scripts/load_env.R")
load("data/summaries_earlier.RData")
load("data/pileup_237.RData")

# ---- Prepare chromosome arms (exclude Y, centromeres) ----
cent_hg19 <- get_centromere_coordinates("hg19", FALSE, ensembl = TRUE)
arms <- gaps(cent_hg19)
arms <- arms[strand(arms) == "*"]
arms_filt <- arms[1:42]
seqlevels(arms_filt, pruning.mode = "coarse") <- seqlevels(arms_filt)[1:23]

message("Using yy_237_gr (unfiltered guides): ", length(yy_237_gr), " regions")
message("  (vs yy_237_filt_gr: ", length(yy_237_filt_gr), " regions)")

# ===================================================================
# 1. p20250924 WT 237
# ===================================================================
message("\n--- Contingency: p20250924 WT237 (with rescue, seed=3) ---")
c237_wt_resc <- build_contingency_table(
  p20250924_237_wt$bp_filt_grl, yy_237_gr,
  genome = arms_filt, seed = 3,
  breakpoint_folder = "/u/project/yeastyin/chovanec/projects/novaseq251007/BJ_RNP/human/breakpointR_diploid/data_filtered/",
  num_threads = 10)

message("WT237 rescue contingency:")
print(c237_wt_resc$contigency)
ct_wt_resc <- chisq.test(c237_wt_resc$contigency)
message("p-value: ", ct_wt_resc$p.value)
message("Expected from HTML: p = 0.5204, Cut_sites Present ~ 248")

# Non-rescue
c237_wt_filt <- build_contingency_table(
  p20250924_237_wt$bp_filt_grl, yy_237_gr,
  genome = arms_filt, seed = 3)
ct_wt_filt <- chisq.test(c237_wt_filt$contigency)
message("WT237 filt p-value: ", ct_wt_filt$p.value)

# ===================================================================
# 2. p20250924 d10a
# ===================================================================
message("\n--- Contingency: p20250924 d10a (with rescue, seed=3) ---")
c237_d10a_resc <- build_contingency_table(
  p20250924_237_d10a$bp_filt_grl, yy_237_gr,
  genome = arms_filt, seed = 3,
  breakpoint_folder = "/u/project/yeastyin/chovanec/projects/novaseq251007/BJ_RNP/human/breakpointR_diploid/data_filtered/",
  num_threads = 10)

message("d10a rescue contingency:")
print(c237_d10a_resc$contigency)
ct_d10a_resc <- chisq.test(c237_d10a_resc$contigency)
message("p-value: ", ct_d10a_resc$p.value)
message("Expected from HTML: p = 0.637, Cut_sites Present ~ 1303")

# Non-rescue
c237_d10a_filt <- build_contingency_table(
  p20250924_237_d10a$bp_filt_grl, yy_237_gr,
  genome = arms_filt, seed = 3)
ct_d10a_filt <- chisq.test(c237_d10a_filt$contigency)
message("d10a filt p-value: ", ct_d10a_filt$p.value)

# ===================================================================
# 3. p20250924 h840a
# ===================================================================
message("\n--- Contingency: p20250924 h840a (with rescue, seed=3) ---")
c237_h840a_resc <- build_contingency_table(
  p20250924_237_h840a$bp_filt_grl, yy_237_gr,
  genome = arms_filt, seed = 3,
  breakpoint_folder = "/u/project/yeastyin/chovanec/projects/novaseq251007/BJ_RNP/human/breakpointR_diploid/data_filtered/",
  num_threads = 10)

message("h840a rescue contingency:")
print(c237_h840a_resc$contigency)
ct_h840a_resc <- chisq.test(c237_h840a_resc$contigency)
message("p-value: ", ct_h840a_resc$p.value)
message("Expected from HTML: p = 0.001592")

# Non-rescue
c237_h840a_filt <- build_contingency_table(
  p20250924_237_h840a$bp_filt_grl, yy_237_gr,
  genome = arms_filt, seed = 3)
ct_h840a_filt <- chisq.test(c237_h840a_filt$contigency)
message("h840a filt p-value: ", ct_h840a_filt$p.value)

# ===================================================================
# Save RDS files (overwrite with corrected values)
# ===================================================================
saveRDS(c237_wt_resc, "data/c237_wt_cont_tab_resc.rds")
saveRDS(c237_d10a_resc, "data/c237_d10a_cont_tab_resc.rds")
saveRDS(c237_h840a_resc, "data/c237_h840a_cont_tab_resc.rds")
message("Saved corrected RDS files to data/")

# ===================================================================
# Save contingency tables to tables_10conditions/
# ===================================================================
tab_dir <- "data/tables_10conditions/"

for (name_tab in list(
  list("c237_wt_rescue", c237_wt_resc),
  list("c237_d10a_rescue", c237_d10a_resc),
  list("c237_h840a_rescue", c237_h840a_resc),
  list("c237_wt_filt", c237_wt_filt),
  list("c237_d10a_filt", c237_d10a_filt),
  list("c237_h840a_filt", c237_h840a_filt)
)) {
  ct <- name_tab[[2]]$contigency
  write.table(ct, file.path(tab_dir, paste0("contingency_", name_tab[[1]], ".txt")),
              sep = "\t", quote = FALSE)
}
message("Updated contingency tables in ", tab_dir)

# Update enrichment_237_summary.txt — old 3 rows only
# Read existing table, update old rows, write back
existing <- read.delim(file.path(tab_dir, "enrichment_237_summary.txt"), stringsAsFactors = FALSE)
old_rows <- data.frame(
  condition = c("p20250924_WT237", "p20250924_d10a", "p20250924_h840a"),
  n_cells = c(p20250924_237_wt$n_cells, p20250924_237_d10a$n_cells,
              p20250924_237_h840a$n_cells),
  rescue_CutSites_present = c(c237_wt_resc$contigency["Cut_sites", "Present"],
                               c237_d10a_resc$contigency["Cut_sites", "Present"],
                               c237_h840a_resc$contigency["Cut_sites", "Present"]),
  rescue_Random_present = c(c237_wt_resc$contigency["Random_sites", "Present"],
                             c237_d10a_resc$contigency["Random_sites", "Present"],
                             c237_h840a_resc$contigency["Random_sites", "Present"]),
  rescue_pvalue = c(ct_wt_resc$p.value, ct_d10a_resc$p.value, ct_h840a_resc$p.value),
  filt_pvalue = c(ct_wt_filt$p.value, ct_d10a_filt$p.value, ct_h840a_filt$p.value),
  stringsAsFactors = FALSE
)

# Match column names to existing table
colnames(old_rows) <- colnames(existing)

# Replace old rows, keep new rows
new_rows <- existing[!existing[[1]] %in% old_rows[[1]], , drop = FALSE]
updated <- rbind(old_rows, new_rows)

write.table(updated, file.path(tab_dir, "enrichment_237_summary.txt"),
            sep = "\t", row.names = FALSE, quote = FALSE)
message("Updated enrichment_237_summary.txt (old 3 rows only)")

# ===================================================================
# Summary comparison
# ===================================================================
message("\n=== COMPARISON WITH HTML REFERENCE ===")
message("HTML rescue p-values: WT=0.5204, d10a=0.637, h840a=0.001592")
message("Got rescue p-values:  WT=", round(ct_wt_resc$p.value, 4),
        ", d10a=", round(ct_d10a_resc$p.value, 4),
        ", h840a=", round(ct_h840a_resc$p.value, 6))

message("\nDone: 09_verify_old_c237.R")
```
