## Supplementary material for "Genome-wide mapping of Cas9-induced sister chromatid exchange across single and over 200 genomic targets": Cas9_RNP_scripts.zip: 09b_237_enrichment_summary.R

```
#!/usr/bin/env Rscript
# 09b_237_enrichment_summary.R
# Build summary table + save contingency text files from pre-computed RDS
# (Avoids re-running the expensive rescue computation)

message("=== 09b_237_enrichment_summary.R ===")
source("scripts/load_env.R")
load("data/pileup_237.RData")

# Load staged RDS files
c237_wt_resc <- readRDS("data/c237_wt_cont_tab_resc.rds")
c237_d10a_resc <- readRDS("data/c237_d10a_cont_tab_resc.rds")
c237_h840a_resc <- readRDS("data/c237_h840a_cont_tab_resc.rds")
c237_24hrs_resc <- readRDS("data/c237_24hrs_cont_tab_resc.rds")
c237_brdu_resc <- readRDS("data/c237_brdu_cont_tab_resc.rds")

# Chi-square on rescue results
ct_wt_resc <- chisq.test(c237_wt_resc$contigency)
ct_d10a_resc <- chisq.test(c237_d10a_resc$contigency)
ct_h840a_resc <- chisq.test(c237_h840a_resc$contigency)
ct_24hrs_resc <- chisq.test(c237_24hrs_resc$contigency)
ct_brdu_resc <- chisq.test(c237_brdu_resc$contigency)

# Also run "no rescue" (simple overlap) — these are fast
load("data/summaries_earlier.RData")
load("data/summaries_new10.RData")

cent_hg19 <- get_centromere_coordinates("hg19", FALSE, ensembl = TRUE)
arms <- gaps(cent_hg19)
arms <- arms[strand(arms) == "*"]
arms_filt <- arms[1:42]
seqlevels(arms_filt, pruning.mode = "coarse") <- seqlevels(arms_filt)[1:23]

c237_wt_filt <- build_contingency_table(
  p20250924_237_wt$bp_filt_grl, yy_237_gr, genome = arms_filt, seed = 3)
ct_wt_filt <- chisq.test(c237_wt_filt$contigency)

c237_d10a_filt <- build_contingency_table(
  p20250924_237_d10a$bp_filt_grl, yy_237_gr, genome = arms_filt, seed = 3)
ct_d10a_filt <- chisq.test(c237_d10a_filt$contigency)

c237_h840a_filt <- build_contingency_table(
  p20250924_237_h840a$bp_filt_grl, yy_237_gr, genome = arms_filt, seed = 3)
ct_h840a_filt <- chisq.test(c237_h840a_filt$contigency)

c237_24hrs_filt <- build_contingency_table(
  new_237_24hrs$bp_filt_grl, yy_237_gr, genome = arms_filt, seed = 3)
ct_24hrs_filt <- chisq.test(c237_24hrs_filt$contigency)

c237_brdu_filt <- build_contingency_table(
  new_237_brdu$bp_filt_grl, yy_237_gr, genome = arms_filt, seed = 3)
ct_brdu_filt <- chisq.test(c237_brdu_filt$contigency)

# Summary table
enrichment_summary <- data.frame(
  condition = c("p20250924_WT237", "p20250924_d10a", "p20250924_h840a",
                "237cuts_24hrs", "237cuts_24ws_BrdU"),
  n_cells = c(p20250924_237_wt$n_cells, p20250924_237_d10a$n_cells,
              p20250924_237_h840a$n_cells,
              new_237_24hrs$n_cells, new_237_brdu$n_cells),
  rescue_CutSites_present = c(c237_wt_resc$contigency["Cut_sites", "Present"],
                         c237_d10a_resc$contigency["Cut_sites", "Present"],
                         c237_h840a_resc$contigency["Cut_sites", "Present"],
                         c237_24hrs_resc$contigency["Cut_sites", "Present"],
                         c237_brdu_resc$contigency["Cut_sites", "Present"]),
  rescue_Random_present = c(c237_wt_resc$contigency["Random_sites", "Present"],
                            c237_d10a_resc$contigency["Random_sites", "Present"],
                            c237_h840a_resc$contigency["Random_sites", "Present"],
                            c237_24hrs_resc$contigency["Random_sites", "Present"],
                            c237_brdu_resc$contigency["Random_sites", "Present"]),
  rescue_pvalue = c(ct_wt_resc$p.value, ct_d10a_resc$p.value,
                    ct_h840a_resc$p.value,
                    ct_24hrs_resc$p.value, ct_brdu_resc$p.value),
  filt_pvalue = c(ct_wt_filt$p.value, ct_d10a_filt$p.value,
                  ct_h840a_filt$p.value,
                  ct_24hrs_filt$p.value, ct_brdu_filt$p.value)
)

write.table(enrichment_summary, file.path(tab_dir, "enrichment_237_summary.txt"),
            sep = "\t", row.names = FALSE, quote = FALSE)
message("\nEnrichment summary:")
print(enrichment_summary)

# Save individual contingency tables as text
for (name_tab in list(
  list("c237_wt_rescue", c237_wt_resc),
  list("c237_d10a_rescue", c237_d10a_resc),
  list("c237_h840a_rescue", c237_h840a_resc),
  list("c237_24hrs_rescue", c237_24hrs_resc),
  list("c237_brdu_rescue", c237_brdu_resc),
  list("c237_wt_filt", c237_wt_filt),
  list("c237_d10a_filt", c237_d10a_filt),
  list("c237_h840a_filt", c237_h840a_filt),
  list("c237_24hrs_filt", c237_24hrs_filt),
  list("c237_brdu_filt", c237_brdu_filt)
)) {
  ct <- name_tab[[2]]$contigency
  write.table(ct, file.path(tab_dir, paste0("contingency_", name_tab[[1]], ".txt")),
              sep = "\t", quote = FALSE)
}
message("Contingency tables saved to ", tab_dir)

message("Done: 09b_237_enrichment_summary.R")
```
