## Supplementary material for "Genome-wide mapping of Cas9-induced sister chromatid exchange across single and over 200 genomic targets": Cas9_RNP_scripts.zip: 09d_237_enrichment_top10pct_sce.R

```
#!/usr/bin/env Rscript
# 09d_237_enrichment_top10pct_sce.R
# Enrichment analysis restricted to cells in the TOP 10% of SCE (post_filt)
# per condition. Uses quantile threshold (not fixed cutoff like 09b).

message("=== 09d_237_enrichment_top10pct_sce.R ===")
source("scripts/load_env.R")
load("data/summaries_earlier.RData")
load("data/summaries_new10.RData")
load("data/pileup_237.RData")

suffix <- "_top10pct_sce"
int_dir <- "data/tables_10conditions/intermediate"
out_dir <- "data/tables_10conditions"

# ---- Chromosome arms ----
cent_hg19 <- get_centromere_coordinates("hg19", FALSE, ensembl = TRUE)
arms <- gaps(cent_hg19)
arms <- arms[strand(arms) == "*"]
arms_filt <- arms[1:42]
seqlevels(arms_filt, pruning.mode = "coarse") <- seqlevels(arms_filt)[1:23]

# ---- Build QC data with post_filt per condition ----
# Old conditions from summary_p20250924_filt
old_qc <- summary_p20250924_filt %>%
  filter(condition %in% c("bj_rnp_seq3_237", "bj_rnp_seq3_237_d10a", "bj_rnp_seq3_237_h840a")) %>%
  select(sample, condition, post_filt)

# New conditions from 10M CSVs
new_base_path <- "/u/project/yeastyin/yeastyin/projects/Cas9_RNP/data/bpR_output"

read_new_qc <- function(csv_path) {
  df <- read.csv(csv_path)
  df$sample <- sub("^([^\\.]*\\.[^\\.]*).*", "\\1", df$.id)
  df_filt <- df[df$background.estimate > 0 & df$background.estimate <= 0.08 &
                df$med.reads.per.MB > 2 & (df$wc.perc + df$uknw.perc) < 75, ]
  df_filt[, c("sample", "post_filt")]
}

qc_24hrs <- read_new_qc(file.path(new_base_path, "BJ_Cas9_RNP_237cuts_24hrs_16hrs", "breakPoint_qc_summary_10M.csv"))
qc_brdu  <- read_new_qc(file.path(new_base_path, "BJ_Cas9_RNP_237cuts_24ws_40hrs_BrdU", "breakPoint_qc_summary_10M.csv"))

# ---- Helper: get top 10% threshold per condition ----
get_top10_samples <- function(qc_df) {
  threshold <- quantile(qc_df$post_filt, probs = 0.90, na.rm = TRUE)
  high_samples <- qc_df$sample[qc_df$post_filt >= threshold]
  message("  Top 10% threshold: post_filt >= ", threshold,
          " (", length(high_samples), " / ", nrow(qc_df), " cells)")
  high_samples
}

# ---- Helper: subset rescue contingency table to high-SCE cells ----
subset_rescue_contingency <- function(rds_path, high_samples, label) {
  resc <- readRDS(rds_path)
  scd <- resc$sample_counts_df

  scd_sub <- scd[scd$sample %in% high_samples, ]
  message(label, ": ", nrow(scd), " -> ", nrow(scd_sub), " cells (top 10% SCE)")

  contingency <- matrix(
    c(sum(scd_sub$roi_overlap), sum(scd_sub$roi_nonoverlap),
      sum(scd_sub$rand_overlap), sum(scd_sub$rand_nonoverlap)),
    nrow = 2, byrow = TRUE,
    dimnames = list(Region = c("Cut_sites", "Random_sites"), Breakpoint = c("Present", "Absent")))

  list(sample_counts_df = scd_sub, contigency = contingency)
}

# ---- Helper: subset bp_filt_grl to high-SCE cells ----
subset_grl_by_samples <- function(grl, high_samples) {
  grl_names <- sub("^([^\\.]*\\.[^\\.]*).*", "\\1", names(grl))
  keep <- grl_names %in% high_samples
  grl[keep]
}

# ===================================================================
# 1. WT237 (rescue)
# ===================================================================
message("\n--- WT237 ---")
wt_qc <- old_qc %>% filter(condition == "bj_rnp_seq3_237")
wt_high <- get_top10_samples(wt_qc)
c237_wt_resc_top10 <- subset_rescue_contingency(
  file.path(int_dir, "c237_wt_cont_tab_resc.rds"), wt_high, "WT237 rescue")
saveRDS(c237_wt_resc_top10, file.path(int_dir, paste0("c237_wt_cont_tab_resc", suffix, ".rds")))
print(c237_wt_resc_top10$contigency)
ct_wt_resc <- chisq.test(c237_wt_resc_top10$contigency)
message("p-value: ", ct_wt_resc$p.value)

# WT237 (no rescue / filt)
grl_wt_top10 <- subset_grl_by_samples(p20250924_237_wt$bp_filt_grl, wt_high)
message("WT237 filt: ", length(p20250924_237_wt$bp_filt_grl), " -> ", length(grl_wt_top10), " elements")
c237_wt_filt_top10 <- build_contingency_table(grl_wt_top10, yy_237_gr, genome = arms_filt, seed = 3)
ct_wt_filt <- chisq.test(c237_wt_filt_top10$contigency)
message("p-value (no rescue): ", ct_wt_filt$p.value)

# ===================================================================
# 2. d10a (rescue)
# ===================================================================
message("\n--- d10a ---")
d10a_qc <- old_qc %>% filter(condition == "bj_rnp_seq3_237_d10a")
d10a_high <- get_top10_samples(d10a_qc)
c237_d10a_resc_top10 <- subset_rescue_contingency(
  file.path(int_dir, "c237_d10a_cont_tab_resc.rds"), d10a_high, "d10a rescue")
saveRDS(c237_d10a_resc_top10, file.path(int_dir, paste0("c237_d10a_cont_tab_resc", suffix, ".rds")))
print(c237_d10a_resc_top10$contigency)
ct_d10a_resc <- chisq.test(c237_d10a_resc_top10$contigency)
message("p-value: ", ct_d10a_resc$p.value)

grl_d10a_top10 <- subset_grl_by_samples(p20250924_237_d10a$bp_filt_grl, d10a_high)
message("d10a filt: ", length(p20250924_237_d10a$bp_filt_grl), " -> ", length(grl_d10a_top10), " elements")
c237_d10a_filt_top10 <- build_contingency_table(grl_d10a_top10, yy_237_gr, genome = arms_filt, seed = 3)
ct_d10a_filt <- chisq.test(c237_d10a_filt_top10$contigency)
message("p-value (no rescue): ", ct_d10a_filt$p.value)

# ===================================================================
# 3. h840a (rescue)
# ===================================================================
message("\n--- h840a ---")
h840a_qc <- old_qc %>% filter(condition == "bj_rnp_seq3_237_h840a")
h840a_high <- get_top10_samples(h840a_qc)
c237_h840a_resc_top10 <- subset_rescue_contingency(
  file.path(int_dir, "c237_h840a_cont_tab_resc.rds"), h840a_high, "h840a rescue")
saveRDS(c237_h840a_resc_top10, file.path(int_dir, paste0("c237_h840a_cont_tab_resc", suffix, ".rds")))
print(c237_h840a_resc_top10$contigency)
ct_h840a_resc <- chisq.test(c237_h840a_resc_top10$contigency)
message("p-value: ", ct_h840a_resc$p.value)

grl_h840a_top10 <- subset_grl_by_samples(p20250924_237_h840a$bp_filt_grl, h840a_high)
message("h840a filt: ", length(p20250924_237_h840a$bp_filt_grl), " -> ", length(grl_h840a_top10), " elements")
c237_h840a_filt_top10 <- build_contingency_table(grl_h840a_top10, yy_237_gr, genome = arms_filt, seed = 3)
ct_h840a_filt <- chisq.test(c237_h840a_filt_top10$contigency)
message("p-value (no rescue): ", ct_h840a_filt$p.value)

# ===================================================================
# 4. 237cuts_24hrs (rescue)
# ===================================================================
message("\n--- 237cuts_24hrs ---")
hrs24_high <- get_top10_samples(qc_24hrs)
c237_24hrs_resc_top10 <- subset_rescue_contingency(
  "data/c237_24hrs_cont_tab_resc.rds", hrs24_high, "24hrs rescue")
saveRDS(c237_24hrs_resc_top10, file.path(int_dir, paste0("c237_24hrs_cont_tab_resc", suffix, ".rds")))
print(c237_24hrs_resc_top10$contigency)
ct_24hrs_resc <- chisq.test(c237_24hrs_resc_top10$contigency)
message("p-value: ", ct_24hrs_resc$p.value)

grl_24hrs_top10 <- subset_grl_by_samples(new_237_24hrs$bp_filt_grl, hrs24_high)
message("24hrs filt: ", length(new_237_24hrs$bp_filt_grl), " -> ", length(grl_24hrs_top10), " elements")
c237_24hrs_filt_top10 <- build_contingency_table(grl_24hrs_top10, yy_237_gr, genome = arms_filt, seed = 3)
ct_24hrs_filt <- chisq.test(c237_24hrs_filt_top10$contigency)
message("p-value (no rescue): ", ct_24hrs_filt$p.value)

# ===================================================================
# 5. 237cuts_24ws_BrdU (rescue)
# ===================================================================
message("\n--- 237cuts_24ws_BrdU ---")
brdu_high <- get_top10_samples(qc_brdu)
c237_brdu_resc_top10 <- subset_rescue_contingency(
  "data/c237_brdu_cont_tab_resc.rds", brdu_high, "BrdU rescue")
saveRDS(c237_brdu_resc_top10, file.path(int_dir, paste0("c237_brdu_cont_tab_resc", suffix, ".rds")))
print(c237_brdu_resc_top10$contigency)
ct_brdu_resc <- chisq.test(c237_brdu_resc_top10$contigency)
message("p-value: ", ct_brdu_resc$p.value)

grl_brdu_top10 <- subset_grl_by_samples(new_237_brdu$bp_filt_grl, brdu_high)
message("BrdU filt: ", length(new_237_brdu$bp_filt_grl), " -> ", length(grl_brdu_top10), " elements")
c237_brdu_filt_top10 <- build_contingency_table(grl_brdu_top10, yy_237_gr, genome = arms_filt, seed = 3)
ct_brdu_filt <- chisq.test(c237_brdu_filt_top10$contigency)
message("p-value (no rescue): ", ct_brdu_filt$p.value)

# ===================================================================
# Summary table
# ===================================================================
summary_top10 <- data.frame(
  condition = c("p20250924_WT237", "p20250924_d10a", "p20250924_h840a",
                "237cuts_24hrs", "237cuts_24ws_BrdU"),
  n_cells_total = c(nrow(wt_qc), nrow(d10a_qc), nrow(h840a_qc),
                    nrow(qc_24hrs), nrow(qc_brdu)),
  n_cells_top10 = c(length(wt_high), length(d10a_high), length(h840a_high),
                    length(hrs24_high), length(brdu_high)),
  rescue_CutSites_present = c(c237_wt_resc_top10$contigency["Cut_sites", "Present"],
                               c237_d10a_resc_top10$contigency["Cut_sites", "Present"],
                               c237_h840a_resc_top10$contigency["Cut_sites", "Present"],
                               c237_24hrs_resc_top10$contigency["Cut_sites", "Present"],
                               c237_brdu_resc_top10$contigency["Cut_sites", "Present"]),
  rescue_Random_present = c(c237_wt_resc_top10$contigency["Random_sites", "Present"],
                             c237_d10a_resc_top10$contigency["Random_sites", "Present"],
                             c237_h840a_resc_top10$contigency["Random_sites", "Present"],
                             c237_24hrs_resc_top10$contigency["Random_sites", "Present"],
                             c237_brdu_resc_top10$contigency["Random_sites", "Present"]),
  rescue_pvalue = c(ct_wt_resc$p.value, ct_d10a_resc$p.value, ct_h840a_resc$p.value,
                    ct_24hrs_resc$p.value, ct_brdu_resc$p.value),
  filt_pvalue = c(ct_wt_filt$p.value, ct_d10a_filt$p.value, ct_h840a_filt$p.value,
                  ct_24hrs_filt$p.value, ct_brdu_filt$p.value)
)

message("\n===== Enrichment Summary (Top 10% SCE) =====")
print(summary_top10)

out_path <- file.path(out_dir, paste0("enrichment_237_summary", suffix, ".txt"))
write.table(summary_top10, out_path, sep = "\t", row.names = FALSE, quote = FALSE)
message("\nSaved: ", out_path)

# Save contingency text files
for (item in list(
  list(paste0("c237_wt_rescue", suffix), c237_wt_resc_top10),
  list(paste0("c237_d10a_rescue", suffix), c237_d10a_resc_top10),
  list(paste0("c237_h840a_rescue", suffix), c237_h840a_resc_top10),
  list(paste0("c237_24hrs_rescue", suffix), c237_24hrs_resc_top10),
  list(paste0("c237_brdu_rescue", suffix), c237_brdu_resc_top10)
)) {
  txt_path <- file.path(out_dir, paste0("contingency_", item[[1]], ".txt"))
  ct <- item[[2]]$contigency
  write.table(ct, txt_path, sep = "\t", quote = FALSE)
  message("Saved: ", txt_path)
}

message("\nDone: 09d_237_enrichment_top10pct_sce.R")
```
