## Supplementary material for "Genome-wide mapping of Cas9-induced sister chromatid exchange across single and over 200 genomic targets": Cas9_RNP_scripts.zip: 10_summary.R

```
#!/usr/bin/env Rscript
# 10_summary.R
# Combined rescue rate figure and cell count table across ALL conditions

message("=== 10_summary.R ===")
source("scripts/load_env.R")

# Load rescue results
load("data/xrcc1_rescue.RData")
load("data/lig3_rescue.RData")
load("data/lig4_rescue.RData")
load("data/xrcc4_rescue.RData")

# ---- Combined rescue rate table ----
xrcc1_rescue_summary$gene <- "XRCC1"
lig3_rescue_summary$gene <- "LIG3"
lig4_rescue_summary$gene <- "LIG4"
xrcc4_rescue_summary$gene <- "XRCC4"

all_rescue <- bind_rows(xrcc1_rescue_summary, lig3_rescue_summary,
                        lig4_rescue_summary, xrcc4_rescue_summary)

write.table(all_rescue, file.path(tab_dir, "all_rescue_summary.txt"),
            sep = "\t", row.names = FALSE, quote = FALSE)
message("Saved all_rescue_summary.txt")
print(all_rescue)

# ---- Combined rescue rate figure ----
all_rescue$label <- paste0(all_rescue$gene, ": ", all_rescue$condition)
all_rescue$label <- factor(all_rescue$label, levels = rev(all_rescue$label))

p_rescue <- ggplot(all_rescue, aes(x = label, y = rescue_rate, fill = gene)) +
  geom_col() +
  geom_text(aes(label = paste0(rescued, "/", total_cells)), hjust = -0.1, size = 2.5) +
  coord_flip() +
  facet_wrap(~gene, scales = "free_y", ncol = 1) +
  scale_y_continuous(expand = expansion(mult = c(0, 0.3)), labels = scales::percent) +
  theme_classic() +
  theme(strip.background = element_blank(),
        legend.position = "none") +
  xlab(NULL) + ylab("Rescue rate") +
  scale_fill_manual(values = c("XRCC1" = "#E41A1C", "LIG3" = "#377EB8",
                                "LIG4" = "#4DAF4A", "XRCC4" = "#984EA3"))

ggsave("rescue_rate_all_conditions.pdf", p_rescue, grDevices::cairo_pdf, fig_dir,
       height = 6, width = 8)
message("Saved rescue_rate_all_conditions.pdf")

# ---- Combined cell count table ----
counts_earlier <- read.table(file.path(tab_dir, "cell_counts_earlier.txt"),
                             header = TRUE, sep = "\t")
counts_new <- read.table(file.path(tab_dir, "cell_counts_new10.txt"),
                          header = TRUE, sep = "\t")

all_counts <- bind_rows(counts_earlier, counts_new)
write.table(all_counts, file.path(tab_dir, "cell_counts_all.txt"),
            sep = "\t", row.names = FALSE, quote = FALSE)
message("Saved cell_counts_all.txt")
print(all_counts)

message("Done: 10_summary.R")
```
