## Supplementary material for "Genome-wide mapping of Cas9-induced sister chromatid exchange across single and over 200 genomic targets": Cas9_RNP_scripts.zip: 11_sce_violin.R

```
#!/usr/bin/env Rscript
# 11_sce_violin.R
# SCE quantification violin plots: pre_filt and post_filt for all 15 BJ conditions
# (5 earlier p20250924 + 10 new conditions)

message("=== 11_sce_violin.R ===")
source("scripts/load_env.R")
load("data/summaries_earlier.RData")
load("data/summaries_new10.RData")

# ---- Combine all BJ conditions ----
# Earlier: p20250924 has the 5 BJ conditions with pre_filt/post_filt
earlier_df <- summary_p20250924_filt %>%
  select(sample, condition, pre_filt, post_filt)

# New: bind all 10 conditions
new_df <- bind_rows(summaries_new_filt) %>%
  select(sample, condition, pre_filt, post_filt)

all_df <- bind_rows(earlier_df, new_df)

# ---- Set condition order ----
cond_levels <- c(
  # Earlier 5
  "bj_rnp_lig3", "bj_rnp_xrcc1",
  "bj_rnp_seq3_237", "bj_rnp_seq3_237_d10a", "bj_rnp_seq3_237_h840a",
  # New 10
  "BJ_Cas9_RNP_Lig3_24hrs", "BJ_Cas9_RNP_Lig3_24wsBrdU",
  "BJ_Cas9_RNP_Xrcc1_24hrs", "BJ_Cas9_RNP_Xrcc1_24wsBrdU",
  "BJ_Cas9_RNP_Lig4_24hrs", "BJ_Cas9_RNP_Lig4_24wsBrdU",
  "BJ_Cas9_RNP_Xrcc4_24hrs", "BJ_Cas9_RNP_Xrcc4_24wsBrdU",
  "BJ_Cas9_RNP_237cuts_24hrs_16hrs", "BJ_Cas9_RNP_237cuts_24ws_40hrs_BrdU"
)

all_df$condition <- factor(all_df$condition, levels = cond_levels)

# ---- Stats summaries (n and max for label placement) ----
stats_pre  <- all_df %>% group_by(condition) %>%
  summarise(n = dplyr::n(), max_val = max(pre_filt), .groups = "drop")
stats_post <- all_df %>% group_by(condition) %>%
  summarise(n = dplyr::n(), max_val = max(post_filt), .groups = "drop")

# ---- p1: pre_filt ----
p1 <- all_df %>%
  ggplot(aes(x = condition, y = pre_filt)) +
  geom_violin(fill = "grey", scale = "width") +
  geom_boxplot(width = 0.15, fill = "white",
               outlier.colour = "grey50", outlier.alpha = 0.5, outlier.size = 0.8) +
  theme_classic() +
  theme(axis.text.x = element_text(angle = 90, vjust = 0.5, hjust = 1)) +
  scale_y_continuous(expand = c(0, 0), limits = c(0, 30)) +
  ylab("pre_filt") + xlab(NULL) +
  ggrepel::geom_text_repel(data = stats_pre,
    aes(y = max_val * 1.15, label = n), size = 3,
    direction = "y", segment.alpha = 0)

# ---- p2: post_filt ----
p2 <- all_df %>%
  ggplot(aes(x = condition, y = post_filt)) +
  geom_violin(fill = "grey", scale = "width") +
  geom_boxplot(width = 0.15, fill = "white",
               outlier.colour = "grey50", outlier.alpha = 0.5, outlier.size = 0.8) +
  theme_classic() +
  theme(axis.text.x = element_text(angle = 90, vjust = 0.5, hjust = 1)) +
  scale_y_continuous(expand = c(0, 0), limits = c(0, 30)) +
  ylab("post_filt") + xlab(NULL) +
  ggrepel::geom_text_repel(data = stats_post,
    aes(y = max_val * 1.15, label = n), size = 3,
    direction = "y", segment.alpha = 0)

# ---- Combined figure ----
p_sce <- p1 + p2 + plot_layout(nrow = 1)

ggsave("SCE_violin_all_conditions.pdf", p_sce, grDevices::cairo_pdf, fig_dir,
       height = 6, width = 14)
message("Saved SCE_violin_all_conditions.pdf")

message("Done: 11_sce_violin.R")
```
