## Supplementary material for "Genome-wide mapping of Cas9-induced sister chromatid exchange across single and over 200 genomic targets": Cas9_RNP_scripts.zip: 11b_sce_violin_coriell.R

```
#!/usr/bin/env Rscript
# 11b_sce_violin_coriell.R
# SCE violin plots for 8 coriell conditions (GM14877_XPC is empty, skipped)
# post_filt is NA for coriell data, so only pre_filt is plotted

message("=== 11b_sce_violin_coriell.R ===")
source("scripts/load_env.R")

base <- "/u/project/yeastyin/yeastyin/projects/coriell/data/bpR_output"
conds <- list.dirs(base, recursive = FALSE, full.names = FALSE)

# Additional conditions from novaseq251219
extra_base_251219 <- "/u/project/yeastyin/chovanec/projects/novaseq251219/split_bam_bwa_hg19_filtered/test_output"
extra_conds_251219 <- c("output_GM04313_ERCC4", "output_GM16693_XPC", "output_GM16633_FANCD2")

# Additional conditions from novaseq251217
extra_base_251217 <- "/u/project/yeastyin/chovanec/projects/novaseq251217/split_bam_bwa_hg19_filtered/test_output"
extra_conds_251217 <- c("output_GM16180_ERCC5", "output_GM16957_ERCC2", "output_GM21072_ERCC3",
                        "output_GM16096_LIG1", "output_GM13155_POLH")

# Load and QC-filter each condition
all_summ <- list()
for (cond in conds) {
  folder <- file.path(base, cond, "data")
  if (!dir.exists(folder)) {
    message("  Skipping ", cond, " — no data/ folder")
    next
  }
  flist <- list.files(folder, pattern = "\\.RData$", full.names = TRUE)
  if (length(flist) == 0) {
    message("  Skipping ", cond, " — no .RData files")
    next
  }
  message("  Loading ", cond, " ...")
  summ <- read_summary(folder, num_threads = 5)
  summ$condition <- cond
  all_summ[[cond]] <- summ
}

# Load extra conditions (strip "output_" prefix for condition name)
extra_all <- c(
  setNames(file.path(extra_base_251219, extra_conds_251219), extra_conds_251219),
  setNames(file.path(extra_base_251217, extra_conds_251217), extra_conds_251217)
)
for (econd in names(extra_all)) {
  folder <- file.path(extra_all[[econd]], "data")
  if (!dir.exists(folder)) {
    message("  Skipping ", econd, " — no data/ folder")
    next
  }
  cond_name <- sub("^output_", "", econd)
  src <- if (grepl("251219", extra_all[[econd]])) "novaseq251219" else "novaseq251217"
  message("  Loading ", cond_name, " (", src, ") ...")
  summ <- read_summary(folder, num_threads = 5)
  summ$condition <- cond_name
  all_summ[[cond_name]] <- summ
}

all_df <- bind_rows(all_summ)
message("Total cells loaded: ", nrow(all_df))

# Apply QC filter (0.06 for GM lines)
all_filt <- all_df %>%
  filter(background.estimate <= 0.06, background.estimate > 0,
         (wc.perc + uknw.perc) < 75,
         med.reads.per.MB >= 2)
message("Cells after QC: ", nrow(all_filt))

# Print per-condition counts
counts <- all_filt %>% group_by(condition) %>%
  summarise(n_pre = dplyr::n(),
            pre_filt_median = median(pre_filt),
            .groups = "drop")
print(counts)

# ---- Condition order ----
cond_levels <- c("GM04959_IKBKAP", "GM05509_XPA",
                 "GM11613_ERCC2", "GM11638_XPC",
                 "GM15709_XPC", "GM16181_ERCC5",
                 "GM16381_XPC", "GM16398_ERCC5",
                 "GM04313_ERCC4", "GM16693_XPC", "GM16633_FANCD2",
                 "GM16180_ERCC5", "GM16957_ERCC2", "GM21072_ERCC3",
                 "GM16096_LIG1", "GM13155_POLH")
all_filt$condition <- factor(all_filt$condition, levels = cond_levels)

# ---- Stats summary ----
stats_pre <- all_filt %>% group_by(condition) %>%
  summarise(n = dplyr::n(), max_val = max(pre_filt), .groups = "drop")

# ---- Violin plot (pre_filt only, since post_filt is NA) ----
p1 <- all_filt %>%
  ggplot(aes(x = condition, y = pre_filt)) +
  geom_violin(fill = "grey", scale = "width") +
  geom_boxplot(width = 0.15, fill = "white",
               outlier.colour = "grey50", outlier.alpha = 0.5, outlier.size = 0.8) +
  theme_classic() +
  theme(axis.text.x = element_text(angle = 90, vjust = 0.5, hjust = 1)) +
  scale_y_continuous(expand = c(0, 0), limits = c(0, 30)) +
  ylab("SCE count (pre_filt)") + xlab(NULL) +
  ggrepel::geom_text_repel(data = stats_pre,
    aes(y = max_val * 1.15, label = n), size = 3,
    direction = "y", segment.alpha = 0)

ggsave("SCE_violin_coriell.pdf", p1, grDevices::cairo_pdf, fig_dir,
       height = 5, width = 13)
message("Saved SCE_violin_coriell.pdf")

# Save filtered summaries for potential reuse
summaries_coriell_filt <- all_filt
save(summaries_coriell_filt, file = "data/summaries_coriell.RData")
message("Saved data/summaries_coriell.RData")

# Cell counts table
cell_counts <- all_df %>% group_by(condition) %>%
  summarise(cells_pre_filter = dplyr::n(), .groups = "drop") %>%
  left_join(all_filt %>% group_by(condition) %>%
    summarise(cells_post_filter = dplyr::n(), .groups = "drop"),
    by = "condition") %>%
  mutate(qc_threshold = 0.06)
# Add conditions with 0 post-filter cells
cell_counts$cells_post_filter[is.na(cell_counts$cells_post_filter)] <- 0

write.table(cell_counts, file.path(tab_dir, "cell_counts_coriell.txt"),
            sep = "\t", row.names = FALSE, quote = FALSE)
message("Saved cell_counts_coriell.txt")

message("Done: 11b_sce_violin_coriell.R")
```
