## Supplementary material for "Genome-wide mapping of Cas9-induced sister chromatid exchange across single and over 200 genomic targets": Cas9_RNP_scripts.zip: 12_genome_pileup.R

```
#!/usr/bin/env Rscript
# 12_genome_pileup.R
# Genome-wide pileup figures for all BJ gene-cut conditions
# (237 pileups already in 08_237_pileup.R; HAP1 in 03_hap1_pileup.R)

message("=== 12_genome_pileup.R ===")
source("scripts/load_env.R")
load("data/summaries_earlier.RData")
load("data/summaries_new10.RData")
load("data/xrcc1_rescue.RData")
load("data/lig3_rescue.RData")
load("data/lig4_rescue.RData")
load("data/xrcc4_rescue.RData")

hg19_cent <- get_centromere_coordinates("hg19", ensembl = TRUE)

# BJ-5ta inversions and blacklist for breakpoint filtering (panels 3-4)
bj_region_filt <- rtracklayer::import.bed("/u/project/yeastyin/chovanec/software/scistrandfig/inst/extdata/regions/BJ-5ta_inv_bl.bed")
hg19_blk <- rtracklayer::import.bed("/u/project/yeastyin/yeastyin/pipelines/sciL3StrandPipe/resources/blacklist/hg19-blacklist.v2.ensembl.bed")

rnp_regions <- GRanges(seqnames = c(19, 17),
                       ranges = IRanges(start = c(44047463, 33307531),
                                        end   = c(44079679, 33336762)))
mcols(rnp_regions)$name <- c("XRCC1", "LIG3")

rnp_lig4 <- GRanges(seqnames = 13, ranges = IRanges(start = 108859790, end = 108870697))
mcols(rnp_lig4)$name <- "LIG4"

rnp_xrcc4 <- GRanges(seqnames = 5, ranges = IRanges(start = 82367495, end = 82643756))
mcols(rnp_xrcc4)$name <- "XRCC4"

suffix_earlier <- "\\.human\\.filt\\.bam\\.RData_filtered|\\.hg19\\.filt\\.bam\\.RData_filtered"
suffix_new     <- "\\.hg19\\.filt\\.bam\\.RData_filtered|\\.hg19\\.filt\\.bam\\.RData"

# Helper: load breakpoints, clean names, return GRL
load_bp <- function(folder, samples, threads = 5, filt, suffix) {
  grl <- breakpoints_to_list(folder, samples = samples, num_threads = threads, filtered = filt)
  gr <- unlist(grl)
  names(gr) <- NULL
  mcols(gr)$sample <- gsub(suffix, "", gr$file)
  split(gr, gr$sample)
}

# Helper: 4 pileup calls + assemble 6-panel (2 regions) or 5-panel (1 region) figure
make_pileup_fig <- function(bp_all_grl, bp_filt_rescue_grl, regions) {
  agg_norm <- plot_aggregate_breaks(bp_all_grl, tilewidth = 1000000, cent = hg19_cent,
    hotspot_pval = NULL, return_individual_plots = TRUE, normalize_counts = TRUE,
    n = length(bp_all_grl))
  agg_raw <- plot_aggregate_breaks(bp_all_grl, tilewidth = 1000000, cent = hg19_cent,
    hotspot_pval = NULL, return_individual_plots = TRUE, normalize_counts = FALSE,
    n = length(bp_all_grl))
  filt_norm <- plot_aggregate_breaks(bp_filt_rescue_grl, tilewidth = 1000000, cent = hg19_cent,
    hotspot_pval = NULL, return_individual_plots = TRUE, normalize_counts = TRUE,
    n = length(bp_filt_rescue_grl))
  filt_raw <- plot_aggregate_breaks(bp_filt_rescue_grl, tilewidth = 1000000, cent = hg19_cent,
    hotspot_pval = NULL, return_individual_plots = TRUE, normalize_counts = FALSE,
    n = length(bp_filt_rescue_grl), regions = regions)

  n_reg <- length(regions)
  if (n_reg == 2) {
    p <- agg_norm[[1]] + agg_raw[[1]] + filt_norm[[1]] + filt_raw[[1]] +
      filt_raw[[4]] + filt_raw[[5]] +
      plot_layout(nrow = 6, heights = c(4, 4, 4, 4, 1, 1))
  } else {
    p <- agg_norm[[1]] + agg_raw[[1]] + filt_norm[[1]] + filt_raw[[1]] +
      filt_raw[[4]] +
      plot_layout(nrow = 5, heights = c(4, 4, 4, 4, 1))
  }
  p
}

# ===================================================================
# 1. p20250808 (XRCC1+LIG3+237, 228 cells) — rescue at both genes
# ===================================================================
message("--- p20250808 genome-wide pileup ---")

p08_folders <- c("/u/project/yeastyin/chovanec/projects/novaseq250815/bj_rnp_xrcc1_lig3_seq3_237/human/breakpointR_diploid/data_filtered/",
                 "/u/project/yeastyin/chovanec/projects/novaseq250919/bj_rnp_xrcc1_lig3_seq3_237/human/breakpointR_diploid/data_filtered/")

bp_p08_all_grl <- load_bp(p08_folders, summary_p20250808_filt$sample, 5, FALSE, suffix_earlier)

bp_p08_filt_grl <- load_bp(p08_folders, summary_p20250808_filt$sample, 5, TRUE, suffix_earlier)
bp_p08_filt_gr <- unlist(bp_p08_filt_grl)
names(bp_p08_filt_gr) <- NULL
mcols(bp_p08_filt_gr)$sample <- gsub(suffix_earlier, "", bp_p08_filt_gr$file)
bp_p08_filt_gr <- plyranges::filter_by_non_overlaps(bp_p08_filt_gr, c(bj_region_filt, hg19_blk))

bp_p08_filt_rescue_gr <- add_rescue_breaks(bp_p08_filt_gr, xrcc1_p20250808, "19:44047463-44079679")
bp_p08_filt_rescue_gr <- add_rescue_breaks(bp_p08_filt_rescue_gr, lig3_p20250808, "17:33307531-33336762")
bp_p08_filt_rescue_grl <- split(bp_p08_filt_rescue_gr, bp_p08_filt_rescue_gr$sample)

p_fig <- make_pileup_fig(bp_p08_all_grl, bp_p08_filt_rescue_grl, rnp_regions)
ggsave("pileup_genome_p20250808.pdf", p_fig, grDevices::cairo_pdf, fig_dir, height = 5, width = 12)
message("Saved pileup_genome_p20250808.pdf")
rm(bp_p08_all_grl, bp_p08_filt_grl, bp_p08_filt_gr, bp_p08_filt_rescue_gr, bp_p08_filt_rescue_grl, p_fig); gc()

# ===================================================================
# 2. p20250409 LIG3 (193 cells)
# ===================================================================
message("--- p20250409 LIG3 genome-wide pileup ---")

p0409_folder <- "/u/project/yeastyin/chovanec/projects/novaseq250806/BJ5ta_RNP_XRCC1_LIG3/hg19/breakpointR_diploid/data_filtered/"
p0409_lig3_samples <- novaseq250806_summary_bj_rnp_filt %>% filter(condition == "BJ_rnp_lig3") %>% pull(sample)

bp_0409lig3_all_grl <- load_bp(p0409_folder, p0409_lig3_samples, 5, FALSE, suffix_earlier)

bp_0409lig3_filt_grl <- load_bp(p0409_folder, p0409_lig3_samples, 5, TRUE, suffix_earlier)
bp_0409lig3_filt_gr <- unlist(bp_0409lig3_filt_grl)
names(bp_0409lig3_filt_gr) <- NULL
mcols(bp_0409lig3_filt_gr)$sample <- gsub(suffix_earlier, "", bp_0409lig3_filt_gr$file)
bp_0409lig3_filt_gr <- plyranges::filter_by_non_overlaps(bp_0409lig3_filt_gr, c(bj_region_filt, hg19_blk))

bp_0409lig3_filt_rescue_gr <- add_rescue_breaks(bp_0409lig3_filt_gr, lig3_p20250409, "17:33307531-33336762")
bp_0409lig3_filt_rescue_grl <- split(bp_0409lig3_filt_rescue_gr, bp_0409lig3_filt_rescue_gr$sample)

p_fig <- make_pileup_fig(bp_0409lig3_all_grl, bp_0409lig3_filt_rescue_grl, rnp_regions)
ggsave("pileup_genome_p20250409_lig3.pdf", p_fig, grDevices::cairo_pdf, fig_dir, height = 5, width = 12)
message("Saved pileup_genome_p20250409_lig3.pdf")
rm(bp_0409lig3_all_grl, bp_0409lig3_filt_grl, bp_0409lig3_filt_gr, bp_0409lig3_filt_rescue_gr, bp_0409lig3_filt_rescue_grl, p_fig); gc()

# ===================================================================
# 3. p20250409 XRCC1 (209 cells)
# ===================================================================
message("--- p20250409 XRCC1 genome-wide pileup ---")

p0409_xrcc1_samples <- novaseq250806_summary_bj_rnp_filt %>% filter(condition == "BJ_rnp_xrcc1") %>% pull(sample)

bp_0409xrcc1_all_grl <- load_bp(p0409_folder, p0409_xrcc1_samples, 5, FALSE, suffix_earlier)

bp_0409xrcc1_filt_grl <- load_bp(p0409_folder, p0409_xrcc1_samples, 5, TRUE, suffix_earlier)
bp_0409xrcc1_filt_gr <- unlist(bp_0409xrcc1_filt_grl)
names(bp_0409xrcc1_filt_gr) <- NULL
mcols(bp_0409xrcc1_filt_gr)$sample <- gsub(suffix_earlier, "", bp_0409xrcc1_filt_gr$file)
bp_0409xrcc1_filt_gr <- plyranges::filter_by_non_overlaps(bp_0409xrcc1_filt_gr, c(bj_region_filt, hg19_blk))

bp_0409xrcc1_filt_rescue_gr <- add_rescue_breaks(bp_0409xrcc1_filt_gr, xrcc1_p20250409, "19:44047463-44079679")
bp_0409xrcc1_filt_rescue_grl <- split(bp_0409xrcc1_filt_rescue_gr, bp_0409xrcc1_filt_rescue_gr$sample)

p_fig <- make_pileup_fig(bp_0409xrcc1_all_grl, bp_0409xrcc1_filt_rescue_grl, rnp_regions)
ggsave("pileup_genome_p20250409_xrcc1.pdf", p_fig, grDevices::cairo_pdf, fig_dir, height = 5, width = 12)
message("Saved pileup_genome_p20250409_xrcc1.pdf")
rm(bp_0409xrcc1_all_grl, bp_0409xrcc1_filt_grl, bp_0409xrcc1_filt_gr, bp_0409xrcc1_filt_rescue_gr, bp_0409xrcc1_filt_rescue_grl, p_fig); gc()

# ===================================================================
# 4. p20250924 LIG3 (471 cells)
# ===================================================================
message("--- p20250924 LIG3 genome-wide pileup ---")

p0924_folder <- "/u/project/yeastyin/chovanec/projects/novaseq251007/BJ_RNP/human/breakpointR_diploid/data_filtered/"
p0924_lig3_samples <- summary_p20250924_filt %>% filter(condition == "bj_rnp_lig3") %>% pull(sample)

bp_0924lig3_all_grl <- load_bp(p0924_folder, p0924_lig3_samples, 5, FALSE, suffix_earlier)

bp_0924lig3_filt_grl <- load_bp(p0924_folder, p0924_lig3_samples, 5, TRUE, suffix_earlier)
bp_0924lig3_filt_gr <- unlist(bp_0924lig3_filt_grl)
names(bp_0924lig3_filt_gr) <- NULL
mcols(bp_0924lig3_filt_gr)$sample <- gsub(suffix_earlier, "", bp_0924lig3_filt_gr$file)
bp_0924lig3_filt_gr <- plyranges::filter_by_non_overlaps(bp_0924lig3_filt_gr, c(bj_region_filt, hg19_blk))

bp_0924lig3_filt_rescue_gr <- add_rescue_breaks(bp_0924lig3_filt_gr, lig3_p20250924, "17:33307531-33336762")
bp_0924lig3_filt_rescue_grl <- split(bp_0924lig3_filt_rescue_gr, bp_0924lig3_filt_rescue_gr$sample)

p_fig <- make_pileup_fig(bp_0924lig3_all_grl, bp_0924lig3_filt_rescue_grl, rnp_regions)
ggsave("pileup_genome_p20250924_lig3.pdf", p_fig, grDevices::cairo_pdf, fig_dir, height = 5, width = 12)
message("Saved pileup_genome_p20250924_lig3.pdf")
rm(bp_0924lig3_all_grl, bp_0924lig3_filt_grl, bp_0924lig3_filt_gr, bp_0924lig3_filt_rescue_gr, bp_0924lig3_filt_rescue_grl, p_fig); gc()

# ===================================================================
# 5. p20250924 XRCC1 (234 cells)
# ===================================================================
message("--- p20250924 XRCC1 genome-wide pileup ---")

p0924_xrcc1_samples <- summary_p20250924_filt %>% filter(condition == "bj_rnp_xrcc1") %>% pull(sample)

bp_0924xrcc1_all_grl <- load_bp(p0924_folder, p0924_xrcc1_samples, 5, FALSE, suffix_earlier)

bp_0924xrcc1_filt_grl <- load_bp(p0924_folder, p0924_xrcc1_samples, 5, TRUE, suffix_earlier)
bp_0924xrcc1_filt_gr <- unlist(bp_0924xrcc1_filt_grl)
names(bp_0924xrcc1_filt_gr) <- NULL
mcols(bp_0924xrcc1_filt_gr)$sample <- gsub(suffix_earlier, "", bp_0924xrcc1_filt_gr$file)
bp_0924xrcc1_filt_gr <- plyranges::filter_by_non_overlaps(bp_0924xrcc1_filt_gr, c(bj_region_filt, hg19_blk))

bp_0924xrcc1_filt_rescue_gr <- add_rescue_breaks(bp_0924xrcc1_filt_gr, xrcc1_p20250924, "19:44047463-44079679")
bp_0924xrcc1_filt_rescue_grl <- split(bp_0924xrcc1_filt_rescue_gr, bp_0924xrcc1_filt_rescue_gr$sample)

p_fig <- make_pileup_fig(bp_0924xrcc1_all_grl, bp_0924xrcc1_filt_rescue_grl, rnp_regions)
ggsave("pileup_genome_p20250924_xrcc1.pdf", p_fig, grDevices::cairo_pdf, fig_dir, height = 5, width = 12)
message("Saved pileup_genome_p20250924_xrcc1.pdf")
rm(bp_0924xrcc1_all_grl, bp_0924xrcc1_filt_grl, bp_0924xrcc1_filt_gr, bp_0924xrcc1_filt_rescue_gr, bp_0924xrcc1_filt_rescue_grl, p_fig); gc()

# ===================================================================
# 6-13. New gene-cut conditions (8 conditions)
# ===================================================================
new_base <- "/u/project/yeastyin/yeastyin/projects/Cas9_RNP/data/bpR_output/"

new_gene_conditions <- list(
  list(name = "Xrcc1_24hrs",    cond = "BJ_Cas9_RNP_Xrcc1_24hrs",    rescue = xrcc1_new_24hrs,    coord = "19:44047463-44079679", regions = rnp_regions),
  list(name = "Xrcc1_24wsBrdU", cond = "BJ_Cas9_RNP_Xrcc1_24wsBrdU", rescue = xrcc1_new_24wsBrdU, coord = "19:44047463-44079679", regions = rnp_regions),
  list(name = "Lig3_24hrs",     cond = "BJ_Cas9_RNP_Lig3_24hrs",     rescue = lig3_new_24hrs,     coord = "17:33307531-33336762", regions = rnp_regions),
  list(name = "Lig3_24wsBrdU",  cond = "BJ_Cas9_RNP_Lig3_24wsBrdU",  rescue = lig3_new_24wsBrdU,  coord = "17:33307531-33336762", regions = rnp_regions),
  list(name = "Lig4_24hrs",     cond = "BJ_Cas9_RNP_Lig4_24hrs",     rescue = lig4_new_24hrs,     coord = "13:108859790-108870697", regions = rnp_lig4),
  list(name = "Lig4_24wsBrdU",  cond = "BJ_Cas9_RNP_Lig4_24wsBrdU",  rescue = lig4_new_24wsBrdU,  coord = "13:108859790-108870697", regions = rnp_lig4),
  list(name = "Xrcc4_24hrs",    cond = "BJ_Cas9_RNP_Xrcc4_24hrs",    rescue = xrcc4_new_24hrs,    coord = "5:82367495-82643756",  regions = rnp_xrcc4),
  list(name = "Xrcc4_24wsBrdU", cond = "BJ_Cas9_RNP_Xrcc4_24wsBrdU", rescue = xrcc4_new_24wsBrdU, coord = "5:82367495-82643756",  regions = rnp_xrcc4)
)

for (item in new_gene_conditions) {
  message("--- ", item$name, " genome-wide pileup ---")

  folder <- file.path(new_base, item$cond, "data_filtered_10M/")
  samples <- summaries_new_filt[[item$cond]]$sample

  bp_all_grl <- load_bp(folder, samples, 5, FALSE, suffix_new)

  bp_filt_grl <- load_bp(folder, samples, 5, TRUE, suffix_new)
  bp_filt_gr <- unlist(bp_filt_grl)
  names(bp_filt_gr) <- NULL
  mcols(bp_filt_gr)$sample <- gsub(suffix_new, "", bp_filt_gr$file)
  bp_filt_gr <- plyranges::filter_by_non_overlaps(bp_filt_gr, c(bj_region_filt, hg19_blk))

  bp_filt_rescue_gr <- add_rescue_breaks(bp_filt_gr, item$rescue, item$coord)
  bp_filt_rescue_grl <- split(bp_filt_rescue_gr, bp_filt_rescue_gr$sample)

  p_fig <- make_pileup_fig(bp_all_grl, bp_filt_rescue_grl, item$regions)
  fname <- paste0("pileup_genome_", item$name, ".pdf")
  ggsave(fname, p_fig, grDevices::cairo_pdf, fig_dir, height = 5, width = 12)
  message("Saved ", fname)

  rm(bp_all_grl, bp_filt_grl, bp_filt_gr, bp_filt_rescue_gr, bp_filt_rescue_grl, p_fig); gc()
}

message("Done: 12_genome_pileup.R")
```
