## Supplementary material for "Genome-wide mapping of Cas9-induced sister chromatid exchange across single and over 200 genomic targets": Cas9_RNP_scripts.zip: 13_heatmap.R

```
#!/usr/bin/env Rscript
# 13_heatmap.R
# Heatmaps for all conditions with rescue data:
#   XRCC1: p20250924, 24hrs, 24wsBrdU
#   LIG3:  p20250924, 24hrs, 24wsBrdU
#   LIG4:  24hrs, 24wsBrdU
#   XRCC4: 24hrs, 24wsBrdU
# Full chromosome heatmap via plot_deeptools_panel + zoom around cut site

message("=== 13_heatmap.R ===")
source("scripts/load_env.R")
# profileplyr class is defined by sciStrandR; do NOT library(profileplyr)
# as it masks sciStrandR::export_deepToolsMat and breaks plot_deeptools_heatmap

load("data/summaries_earlier.RData")
load("data/summaries_new10.RData")
load("data/xrcc1_rescue.RData")
load("data/lig3_rescue.RData")
load("data/lig4_rescue.RData")
load("data/xrcc4_rescue.RData")

deeptools_path <- "/u/project/yeastyin/chovanec/miniconda3/envs/deeptools/bin/plotHeatmap"
blacklist <- "/u/project/yeastyin/chovanec/genomes/hg19-blacklist.v2.ensembl.bed"

heatmap_dir <- "data/heatmaps/"
matrix_dir  <- "data/heatmaps_matrices/"

# ---- Helper: safe wrapper for plot_deeptools_panel ----
safe_heatmap <- function(label, ...) {
  tryCatch({
    plot_deeptools_panel(...)
    message("Full chromosome ", label, " heatmap done")
    TRUE
  }, error = function(e) {
    message("WARNING: ", label, " heatmap failed: ", conditionMessage(e))
    FALSE
  })
}

# ---- Helper: safe wrapper for extract_zoom_heatmap ----
safe_zoom <- function(...) {
  tryCatch(extract_zoom_heatmap(...),
    error = function(e) message("WARNING: zoom heatmap failed: ", conditionMessage(e)))
}

# ---- Helper: zoom extraction from saved matrix ----
extract_zoom_heatmap <- function(mat_path, zoom_gr, rescue_df, out_pdf) {
  # Fall back to matrix_all.rds if final_matrix.rds doesn't exist
  if (!file.exists(mat_path)) {
    alt_path <- sub("_final_matrix\\.rds$", "_matrix_all.rds", mat_path)
    if (file.exists(alt_path)) {
      message("  Using matrix_all.rds (final_matrix.rds not found)")
      mat_path <- alt_path
    } else {
      stop("Neither final_matrix.rds nor matrix_all.rds found")
    }
  }
  mat <- readRDS(mat_path)
  gr_list <- split(zoom_gr, as.factor(zoom_gr))

  # Column metadata (split by panel index suffix)
  coldata <- split.default(SummarizedExperiment::colData(mat),
    sapply(strsplit(names(SummarizedExperiment::colData(mat)), "_"), "[[", 2))

  # Bins falling within zoom window
  desired_bins <- which(
    coldata[[1]]$start_1 > start(gr_list[[1]]) &
    coldata[[1]]$end_1   < end(gr_list[[1]])
  )
  if (length(desired_bins) == 0) {
    message("No bins in zoom window for ", out_pdf, " — skipping")
    return(invisible(NULL))
  }

  mat_z <- mat[, min(desired_bins):max(desired_bins), 1]
  SummarizedExperiment::colData(mat_z) <- coldata[[1]][min(desired_bins):max(desired_bins), ]
  colnames(SummarizedExperiment::colData(mat_z)) <- gsub("_1", "", colnames(SummarizedExperiment::colData(mat_z)))

  # Update sampleData range
  cd_gr    <- makeGRangesFromDataFrame(SummarizedExperiment::colData(mat_z))
  cd_range <- range(cd_gr)
  rownames(sampleData(mat_z)) <- paste(seqnames(cd_range),
    paste(start(cd_range), end(cd_range), sep = "-"), sep = ":")

  # Truncate sample names to match rescue IDs
  rownames(SummarizedExperiment::assay(mat_z, withDimnames = FALSE)) <- sub("^([^\\.]*\\.[^\\.]*).*", "\\1",
    rownames(SummarizedExperiment::assay(mat_z)))
  rownames(mat_z) <- sub("^([^\\.]*\\.[^\\.]*).*", "\\1", rownames(mat_z))

  # Filter to rescue cells, sorted by strand state
  cells_to_plot <- rescue_df %>%
    filter(rescue_break == TRUE) %>%
    mutate(sample = sub("^([^\\.]*\\.[^\\.]*).*", "\\1", ID)) %>%
    arrange(tiled_states) %>%
    pull(sample)

  sample_idx <- match(unique(cells_to_plot), rownames(mat_z))
  sample_idx <- sample_idx[!is.na(sample_idx)]

  if (length(sample_idx) == 0) {
    message("No rescue cells found in matrix for ", out_pdf, " — skipping")
    return(invisible(NULL))
  }
  message("  Zoom: ", length(sample_idx), " rescued cells, ",
          length(desired_bins), " bins")

  mat_z <- mat_z[sample_idx, , ]

  # DeepTools reference point parameters
  mat_z@sampleData$body       <- 0
  mat_z@sampleData$bin.size   <- 200000
  mat_z@sampleData$ref.point  <- "RNP"
  mat_z@sampleData$upstream   <- width(gr_list[[1]]) / 2
  mat_z@sampleData$downstream <- width(gr_list[[1]]) / 2

  plot_deeptools_heatmap(mat_z,
    out_plot = out_pdf,
    deeptools_plot_heatmap_path = deeptools_path,
    heatmap_width = 4, heatmap_height = 4,
    sort_regions = "no",
    interpolation = "nearest",
    additional_flags = c("--refPointLabel RNP --linesAtTickMarks"),
    zmin = "-200", zmax = "200")

  message("  Saved: ", out_pdf)
}

# =====================================================================
# 1. XRCC1 (p20250924) — .rds data, filtered=TRUE works natively
# =====================================================================
message("\n=== XRCC1 p20250924 ===")

xrcc1_folder  <- "/u/project/yeastyin/chovanec/projects/novaseq251007/BJ_RNP/human/breakpointR_diploid/data_filtered/"
xrcc1_samples <- summary_p20250924_filt$sample[summary_p20250924_filt$condition == "bj_rnp_xrcc1"]
message("  ", length(xrcc1_samples), " samples")

safe_heatmap("XRCC1_p20250924",
  folder = xrcc1_folder,
  windows = coordinates2gr("19", "hg19"),
  samples = xrcc1_samples,
  out_pdf = heatmap_dir,
  cluster_out_plot = heatmap_dir,
  max_bin_size = 200000,
  matrices_save_folder = matrix_dir,
  num_threads = 1,
  deeptools_plot_heatmap_path = deeptools_path,
  regions_to_mask = blacklist,
  sort_by_break_position = TRUE,
  data_prefix = "XRCC1_p20250924",
  plot_prefix = "XRCC1_p20250924",
  cpm = TRUE,
  opposite_no_break_category = TRUE,
  overwrite = "none",
  fragments_overlaps = FALSE,
  interpolation = "nearest",
  wsl = FALSE
)

# XRCC1 zoom: ±30.1 Mb (matching original Rmd)
gr_zoom_xrcc1 <- plyranges::stretch(
  coordinates2gr("19:44047463-44079679", "hg19"), extend = 30098608)

safe_zoom(
  mat_path   = file.path(matrix_dir, "XRCC1_p20250924_19_1-59128983_final_matrix.rds"),
  zoom_gr    = gr_zoom_xrcc1,
  rescue_df  = xrcc1_p20250924,
  out_pdf    = file.path(fig_dir, "heatmap_zoom_xrcc1_p20250924.pdf")
)

# =====================================================================
# 2. LIG4 conditions — .rds data in data_filtered_10M, filtered=TRUE works natively
# =====================================================================
base_new <- "/u/project/yeastyin/yeastyin/projects/Cas9_RNP/data/bpR_output"

# ---- LIG4_24hrs ----
message("\n=== LIG4_24hrs ===")
lig4_24hrs_samples <- summaries_new_filt[["BJ_Cas9_RNP_Lig4_24hrs"]]$sample
message("  ", length(lig4_24hrs_samples), " samples")

safe_heatmap("LIG4_24hrs",
  folder = file.path(base_new, "BJ_Cas9_RNP_Lig4_24hrs", "data_filtered_10M"),
  windows = coordinates2gr("13", "hg19"),
  samples = lig4_24hrs_samples,
  out_pdf = heatmap_dir,
  cluster_out_plot = heatmap_dir,
  max_bin_size = 200000,
  matrices_save_folder = matrix_dir,
  num_threads = 1,
  deeptools_plot_heatmap_path = deeptools_path,
  regions_to_mask = blacklist,
  sort_by_break_position = TRUE,
  data_prefix = "LIG4_24hrs",
  plot_prefix = "LIG4_24hrs",
  cpm = TRUE,
  opposite_no_break_category = TRUE,
  overwrite = "none",
  fragments_overlaps = FALSE,
  interpolation = "nearest",
  wsl = FALSE
)

# ---- LIG4_24wsBrdU ----
message("\n=== LIG4_24wsBrdU ===")
lig4_brdu_samples <- summaries_new_filt[["BJ_Cas9_RNP_Lig4_24wsBrdU"]]$sample
message("  ", length(lig4_brdu_samples), " samples")

safe_heatmap("LIG4_24wsBrdU",
  folder = file.path(base_new, "BJ_Cas9_RNP_Lig4_24wsBrdU", "data_filtered_10M"),
  windows = coordinates2gr("13", "hg19"),
  samples = lig4_brdu_samples,
  out_pdf = heatmap_dir,
  cluster_out_plot = heatmap_dir,
  max_bin_size = 200000,
  matrices_save_folder = matrix_dir,
  num_threads = 1,
  deeptools_plot_heatmap_path = deeptools_path,
  regions_to_mask = blacklist,
  sort_by_break_position = TRUE,
  data_prefix = "LIG4_24wsBrdU",
  plot_prefix = "LIG4_24wsBrdU",
  cpm = TRUE,
  opposite_no_break_category = TRUE,
  overwrite = "none",
  fragments_overlaps = FALSE,
  interpolation = "nearest",
  wsl = FALSE
)

# ---- LIG4 zoom: ±60 Mb around 13:108859790-108870697 ----
gr_zoom_lig4 <- plyranges::stretch(
  coordinates2gr("13:108859790-108870697", "hg19"), extend = 60000000)

safe_zoom(
  mat_path   = file.path(matrix_dir, "LIG4_24hrs_13_1-115169878_final_matrix.rds"),
  zoom_gr    = gr_zoom_lig4,
  rescue_df  = lig4_new_24hrs,
  out_pdf    = file.path(fig_dir, "heatmap_zoom_lig4_24hrs.pdf")
)

safe_zoom(
  mat_path   = file.path(matrix_dir, "LIG4_24wsBrdU_13_1-115169878_final_matrix.rds"),
  zoom_gr    = gr_zoom_lig4,
  rescue_df  = lig4_new_24wsBrdU,
  out_pdf    = file.path(fig_dir, "heatmap_zoom_lig4_24wsBrdU.pdf")
)

# =====================================================================
# 3. XRCC1 new conditions (24hrs, 24wsBrdU) — chr19
# =====================================================================

# ---- XRCC1_24hrs ----
message("\n=== XRCC1_24hrs ===")
xrcc1_24hrs_samples <- summaries_new_filt[["BJ_Cas9_RNP_Xrcc1_24hrs"]]$sample
message("  ", length(xrcc1_24hrs_samples), " samples")

safe_heatmap("XRCC1_24hrs",
  folder = file.path(base_new, "BJ_Cas9_RNP_Xrcc1_24hrs", "data_filtered_10M"),
  windows = coordinates2gr("19", "hg19"),
  samples = xrcc1_24hrs_samples,
  out_pdf = heatmap_dir,
  cluster_out_plot = heatmap_dir,
  max_bin_size = 200000,
  matrices_save_folder = matrix_dir,
  num_threads = 1,
  deeptools_plot_heatmap_path = deeptools_path,
  regions_to_mask = blacklist,
  sort_by_break_position = TRUE,
  data_prefix = "XRCC1_24hrs",
  plot_prefix = "XRCC1_24hrs",
  cpm = TRUE,
  opposite_no_break_category = TRUE,
  overwrite = "none",
  fragments_overlaps = FALSE,
  interpolation = "nearest",
  wsl = FALSE
)

safe_zoom(
  mat_path   = file.path(matrix_dir, "XRCC1_24hrs_19_1-59128983_final_matrix.rds"),
  zoom_gr    = gr_zoom_xrcc1,
  rescue_df  = xrcc1_new_24hrs,
  out_pdf    = file.path(fig_dir, "heatmap_zoom_xrcc1_24hrs.pdf")
)

# ---- XRCC1_24wsBrdU ----
message("\n=== XRCC1_24wsBrdU ===")
xrcc1_brdu_samples <- summaries_new_filt[["BJ_Cas9_RNP_Xrcc1_24wsBrdU"]]$sample
message("  ", length(xrcc1_brdu_samples), " samples")

safe_heatmap("XRCC1_24wsBrdU",
  folder = file.path(base_new, "BJ_Cas9_RNP_Xrcc1_24wsBrdU", "data_filtered_10M"),
  windows = coordinates2gr("19", "hg19"),
  samples = xrcc1_brdu_samples,
  out_pdf = heatmap_dir,
  cluster_out_plot = heatmap_dir,
  max_bin_size = 200000,
  matrices_save_folder = matrix_dir,
  num_threads = 1,
  deeptools_plot_heatmap_path = deeptools_path,
  regions_to_mask = blacklist,
  sort_by_break_position = TRUE,
  data_prefix = "XRCC1_24wsBrdU",
  plot_prefix = "XRCC1_24wsBrdU",
  cpm = TRUE,
  opposite_no_break_category = TRUE,
  overwrite = "none",
  fragments_overlaps = FALSE,
  interpolation = "nearest",
  wsl = FALSE
)

safe_zoom(
  mat_path   = file.path(matrix_dir, "XRCC1_24wsBrdU_19_1-59128983_final_matrix.rds"),
  zoom_gr    = gr_zoom_xrcc1,
  rescue_df  = xrcc1_new_24wsBrdU,
  out_pdf    = file.path(fig_dir, "heatmap_zoom_xrcc1_24wsBrdU.pdf")
)

# =====================================================================
# 4. LIG3 conditions (p20250924, 24hrs, 24wsBrdU) — chr17
# =====================================================================
old_folder <- "/u/project/yeastyin/chovanec/projects/novaseq251007/BJ_RNP/human/breakpointR_diploid/data_filtered/"

# LIG3 zoom: ±60 Mb (matching original Rmd)
gr_zoom_lig3 <- plyranges::stretch(
  coordinates2gr("17:33307531-33336762", "hg19"), extend = 60000000)

# ---- LIG3_p20250924 ----
message("\n=== LIG3_p20250924 ===")
lig3_p20250924_samples <- summary_p20250924_filt$sample[summary_p20250924_filt$condition == "bj_rnp_lig3"]
message("  ", length(lig3_p20250924_samples), " samples")

safe_heatmap("LIG3_p20250924",
  folder = old_folder,
  windows = coordinates2gr("17", "hg19"),
  samples = lig3_p20250924_samples,
  out_pdf = heatmap_dir,
  cluster_out_plot = heatmap_dir,
  max_bin_size = 200000,
  matrices_save_folder = matrix_dir,
  num_threads = 1,
  deeptools_plot_heatmap_path = deeptools_path,
  regions_to_mask = blacklist,
  sort_by_break_position = TRUE,
  data_prefix = "LIG3_p20250924",
  plot_prefix = "LIG3_p20250924",
  cpm = TRUE,
  opposite_no_break_category = TRUE,
  overwrite = "none",
  fragments_overlaps = FALSE,
  interpolation = "nearest",
  wsl = FALSE
)

safe_zoom(
  mat_path   = file.path(matrix_dir, "LIG3_p20250924_17_1-81195210_final_matrix.rds"),
  zoom_gr    = gr_zoom_lig3,
  rescue_df  = lig3_p20250924,
  out_pdf    = file.path(fig_dir, "heatmap_zoom_lig3_p20250924.pdf")
)

# ---- LIG3_24hrs ----
message("\n=== LIG3_24hrs ===")
lig3_24hrs_samples <- summaries_new_filt[["BJ_Cas9_RNP_Lig3_24hrs"]]$sample
message("  ", length(lig3_24hrs_samples), " samples")

safe_heatmap("LIG3_24hrs",
  folder = file.path(base_new, "BJ_Cas9_RNP_Lig3_24hrs", "data_filtered_10M"),
  windows = coordinates2gr("17", "hg19"),
  samples = lig3_24hrs_samples,
  out_pdf = heatmap_dir,
  cluster_out_plot = heatmap_dir,
  max_bin_size = 200000,
  matrices_save_folder = matrix_dir,
  num_threads = 1,
  deeptools_plot_heatmap_path = deeptools_path,
  regions_to_mask = blacklist,
  sort_by_break_position = TRUE,
  data_prefix = "LIG3_24hrs",
  plot_prefix = "LIG3_24hrs",
  cpm = TRUE,
  opposite_no_break_category = TRUE,
  overwrite = "none",
  fragments_overlaps = FALSE,
  interpolation = "nearest",
  wsl = FALSE
)

safe_zoom(
  mat_path   = file.path(matrix_dir, "LIG3_24hrs_17_1-81195210_final_matrix.rds"),
  zoom_gr    = gr_zoom_lig3,
  rescue_df  = lig3_new_24hrs,
  out_pdf    = file.path(fig_dir, "heatmap_zoom_lig3_24hrs.pdf")
)

# ---- LIG3_24wsBrdU ----
message("\n=== LIG3_24wsBrdU ===")
lig3_brdu_samples <- summaries_new_filt[["BJ_Cas9_RNP_Lig3_24wsBrdU"]]$sample
message("  ", length(lig3_brdu_samples), " samples")

safe_heatmap("LIG3_24wsBrdU",
  folder = file.path(base_new, "BJ_Cas9_RNP_Lig3_24wsBrdU", "data_filtered_10M"),
  windows = coordinates2gr("17", "hg19"),
  samples = lig3_brdu_samples,
  out_pdf = heatmap_dir,
  cluster_out_plot = heatmap_dir,
  max_bin_size = 200000,
  matrices_save_folder = matrix_dir,
  num_threads = 1,
  deeptools_plot_heatmap_path = deeptools_path,
  regions_to_mask = blacklist,
  sort_by_break_position = TRUE,
  data_prefix = "LIG3_24wsBrdU",
  plot_prefix = "LIG3_24wsBrdU",
  cpm = TRUE,
  opposite_no_break_category = TRUE,
  overwrite = "none",
  fragments_overlaps = FALSE,
  interpolation = "nearest",
  wsl = FALSE
)

safe_zoom(
  mat_path   = file.path(matrix_dir, "LIG3_24wsBrdU_17_1-81195210_final_matrix.rds"),
  zoom_gr    = gr_zoom_lig3,
  rescue_df  = lig3_new_24wsBrdU,
  out_pdf    = file.path(fig_dir, "heatmap_zoom_lig3_24wsBrdU.pdf")
)

# =====================================================================
# 5. XRCC4 conditions (24hrs, 24wsBrdU) — chr5
# =====================================================================

# XRCC4 zoom: ±60 Mb around 5:82367495-82643756
gr_zoom_xrcc4 <- plyranges::stretch(
  coordinates2gr("5:82367495-82643756", "hg19"), extend = 60000000)

# ---- XRCC4_24hrs ----
message("\n=== XRCC4_24hrs ===")
xrcc4_24hrs_samples <- summaries_new_filt[["BJ_Cas9_RNP_Xrcc4_24hrs"]]$sample
message("  ", length(xrcc4_24hrs_samples), " samples")

safe_heatmap("XRCC4_24hrs",
  folder = file.path(base_new, "BJ_Cas9_RNP_Xrcc4_24hrs", "data_filtered_10M"),
  windows = coordinates2gr("5", "hg19"),
  samples = xrcc4_24hrs_samples,
  out_pdf = heatmap_dir,
  cluster_out_plot = heatmap_dir,
  max_bin_size = 200000,
  matrices_save_folder = matrix_dir,
  num_threads = 1,
  deeptools_plot_heatmap_path = deeptools_path,
  regions_to_mask = blacklist,
  sort_by_break_position = TRUE,
  data_prefix = "XRCC4_24hrs",
  plot_prefix = "XRCC4_24hrs",
  cpm = TRUE,
  opposite_no_break_category = TRUE,
  overwrite = "none",
  fragments_overlaps = FALSE,
  interpolation = "nearest",
  wsl = FALSE
)

safe_zoom(
  mat_path   = file.path(matrix_dir, "XRCC4_24hrs_5_1-180915260_final_matrix.rds"),
  zoom_gr    = gr_zoom_xrcc4,
  rescue_df  = xrcc4_new_24hrs,
  out_pdf    = file.path(fig_dir, "heatmap_zoom_xrcc4_24hrs.pdf")
)

# ---- XRCC4_24wsBrdU ----
message("\n=== XRCC4_24wsBrdU ===")
xrcc4_brdu_samples <- summaries_new_filt[["BJ_Cas9_RNP_Xrcc4_24wsBrdU"]]$sample
message("  ", length(xrcc4_brdu_samples), " samples")

safe_heatmap("XRCC4_24wsBrdU",
  folder = file.path(base_new, "BJ_Cas9_RNP_Xrcc4_24wsBrdU", "data_filtered_10M"),
  windows = coordinates2gr("5", "hg19"),
  samples = xrcc4_brdu_samples,
  out_pdf = heatmap_dir,
  cluster_out_plot = heatmap_dir,
  max_bin_size = 200000,
  matrices_save_folder = matrix_dir,
  num_threads = 1,
  deeptools_plot_heatmap_path = deeptools_path,
  regions_to_mask = blacklist,
  sort_by_break_position = TRUE,
  data_prefix = "XRCC4_24wsBrdU",
  plot_prefix = "XRCC4_24wsBrdU",
  cpm = TRUE,
  opposite_no_break_category = TRUE,
  overwrite = "none",
  fragments_overlaps = FALSE,
  interpolation = "nearest",
  wsl = FALSE
)

safe_zoom(
  mat_path   = file.path(matrix_dir, "XRCC4_24wsBrdU_5_1-180915260_final_matrix.rds"),
  zoom_gr    = gr_zoom_xrcc4,
  rescue_df  = xrcc4_new_24wsBrdU,
  out_pdf    = file.path(fig_dir, "heatmap_zoom_xrcc4_24wsBrdU.pdf")
)

message("\nDone: 13_heatmap.R")
```
