## Supplementary material for "Genome-wide mapping of Cas9-induced sister chromatid exchange across single and over 200 genomic targets": Cas9_RNP_scripts.zip: 13_on_target_breakpoint.R

```
#!/usr/bin/env Rscript
# 13_on_target_breakpoint.R
# On-target breakpoint figure: called breaks at target + rescue strand-state calls
# "On target" = breakpointR called a break near target (±3Mb) OR rescue strand-state
# Same conditions as rescue_rate_all_conditions.pdf but with combined call+rescue metric

message("=== 13_on_target_breakpoint.R ===")
source("scripts/load_env.R")
load("data/summaries_earlier.RData")
load("data/summaries_new10.RData")

# ---------------------------------------------------------------
# check_on_target: load one cell, check called break + rescue
# ---------------------------------------------------------------
check_on_target <- function(path, coord_gr, target_region,
                             type = "diploid",
                             bin_width = 10000000, dist_from_gr = 3000000) {
  if (grepl("\\.RData$", path, ignore.case = TRUE)) {
    env <- new.env()
    load(path, envir = env)
    f <- ls(env)[1]
    dehcr_filt <- get(f, envir = env)
  } else {
    f <- load_from_rds(path)
    dehcr_filt <- get(f)
  }
  fragments <- dehcr_filt$fragments
  breaks    <- dehcr_filt$breaks

  # 1. Called break at target (within stretched coord, ±3Mb)
  called_at_target <- length(plyranges::filter_by_overlaps(breaks, target_region)) > 0

  # 2. Rescue strand-state check (same logic as check_location_for_breakpoints)
  coord_grl  <- split(coord_gr)
  coord_arms <- purrr::map(coord_grl, \(x) trim(bins_of_interest(x,
    dist_from_gr = dist_from_gr, bin_width = bin_width)))

  rescue_break <- FALSE
  for (i in seq_along(coord_arms)) {
    L_R_states <- suppressWarnings(state_assignment_bins(coord_arms[[i]], fragments))
    L_R_states <- sort(L_R_states)
    new_states <- paste(L_R_states$states, collapse = "_")
    if (type == "haploid") {
      rescue_break <- rescue_break | (new_states %in% c("ww_cc", "cc_ww"))
    } else {
      rescue_break <- rescue_break | (new_states %in% c("ww_cc", "cc_ww",
        "ww_wc", "wc_ww", "cc_wc", "wc_cc"))
    }
  }

  data.frame(ID = dehcr_filt$ID,
             called_at_target = called_at_target,
             rescue_break = rescue_break,
             on_target = called_at_target | rescue_break)
}

# ---------------------------------------------------------------
# run_check_on_target: parallelized wrapper
# ---------------------------------------------------------------
run_check_on_target <- function(data_folder, coord_str, samples,
                                 type = "diploid", num_threads = 10,
                                 stretch_amount = 6000000,
                                 bin_width = 10000000, dist_from_gr = 3000000,
                                 filtered = TRUE) {
  coord_gr      <- coordinates2gr(coord_str, genome_assembly = "hg19")
  target_region <- plyranges::stretch(coord_gr, stretch_amount)
  data_files    <- get_files_to_load(data_folder, samples, filtered = filtered)
  message("  Processing ", length(data_files), " cells...")

  if (num_threads > 1) {
    pb <- txtProgressBar(max = length(data_files), style = 3)
    progress <- function(n) setTxtProgressBar(pb, n)
    cl <- parallel::makeCluster(num_threads)
    doSNOW::registerDoSNOW(cl)
    out_df <- foreach(f = data_files,
                      .packages = c("sciStrandR", "dplyr", "purrr", "plyranges"),
                      .export = c("check_on_target", "bins_of_interest",
                                  "state_assignment_bins"),
                      .options.snow = list(progress = progress),
                      .combine = rbind) %dopar% {
      check_on_target(f, coord_gr, target_region, type = type,
                       bin_width = bin_width, dist_from_gr = dist_from_gr)
    }
    parallel::stopCluster(cl)
    close(pb)
  } else {
    out_lst <- lapply(data_files, function(f) {
      check_on_target(f, coord_gr, target_region, type = type,
                       bin_width = bin_width, dist_from_gr = dist_from_gr)
    })
    out_df <- bind_rows(out_lst)
  }
  rownames(out_df) <- NULL
  out_df
}

# ---------------------------------------------------------------
# summarize_on_target: per-condition summary
# ---------------------------------------------------------------
summarize_on_target <- function(df) {
  data.frame(
    total_cells      = nrow(df),
    called_at_target = sum(df$called_at_target),
    rescue_only      = sum(df$rescue_break & !df$called_at_target),
    on_target        = sum(df$on_target)
  )
}

# ---------------------------------------------------------------
# Paths
# ---------------------------------------------------------------
chovanec <- "/u/project/yeastyin/chovanec/projects"
new_base <- "/u/project/yeastyin/yeastyin/projects/Cas9_RNP/data/bpR_output"

p20250808_dirs <- c(
  file.path(chovanec, "novaseq250815/bj_rnp_xrcc1_lig3_seq3_237/human/breakpointR_diploid/data_filtered"),
  file.path(chovanec, "novaseq250919/bj_rnp_xrcc1_lig3_seq3_237/human/breakpointR_diploid/data_filtered"))
p20250409_dir    <- file.path(chovanec, "novaseq250806/BJ5ta_RNP_XRCC1_LIG3/hg19/breakpointR_diploid/data_filtered")
p20250924_dir    <- file.path(chovanec, "novaseq251007/BJ_RNP/human/breakpointR_diploid/data_filtered")
hap1_haploid_dir <- file.path(chovanec, "novaseq250806/HAP1_RNP_XRCC1_LIG3/hg19/breakpointR_haploid/data_filtered")
hap1_diploid_dir <- file.path(chovanec, "novaseq250806/HAP1_RNP_XRCC1_LIG3/hg19/breakpointR_diploid/data_filtered")

xrcc1_coord_str <- "19:44047463-44079679"
lig3_coord_str  <- "17:33307531-33336762"
lig4_coord_str  <- "13:108859790-108870697"
xrcc4_coord_str <- "5:82367495-82643756"

# ===================================================================
# XRCC1 (7 conditions)
# ===================================================================
message("\n========== XRCC1 ==========")

message("--- p20250808 ---")
xrcc1_p08_ot <- run_check_on_target(p20250808_dirs, xrcc1_coord_str,
  summary_p20250808_filt$sample, type = "diploid")

message("--- p20250409 ---")
xrcc1_0409_ot <- run_check_on_target(p20250409_dir, xrcc1_coord_str,
  novaseq250806_summary_bj_rnp_filt$sample[novaseq250806_summary_bj_rnp_filt$condition == "BJ_rnp_xrcc1"],
  type = "diploid")

message("--- p20250924 ---")
xrcc1_0924_ot <- run_check_on_target(p20250924_dir, xrcc1_coord_str,
  summary_p20250924_filt %>% filter(condition == "bj_rnp_xrcc1") %>% pull(sample),
  type = "diploid")

message("--- HAP1 haploid ---")
xrcc1_hap1h_ot <- run_check_on_target(hap1_haploid_dir, xrcc1_coord_str,
  summary_hap1_haploid_filt$sample, type = "haploid")

message("--- HAP1 diploid ---")
xrcc1_hap1d_ot <- run_check_on_target(hap1_diploid_dir, xrcc1_coord_str,
  summary_hap1_diploid_filt$sample, type = "diploid")

message("--- Xrcc1_24hrs ---")
xrcc1_24hrs_ot <- run_check_on_target(
  file.path(new_base, "BJ_Cas9_RNP_Xrcc1_24hrs/data_filtered_10M"), xrcc1_coord_str,
  summaries_new_filt[["BJ_Cas9_RNP_Xrcc1_24hrs"]]$sample, type = "diploid")

message("--- Xrcc1_24wsBrdU ---")
xrcc1_brdu_ot <- run_check_on_target(
  file.path(new_base, "BJ_Cas9_RNP_Xrcc1_24wsBrdU/data_filtered_10M"), xrcc1_coord_str,
  summaries_new_filt[["BJ_Cas9_RNP_Xrcc1_24wsBrdU"]]$sample, type = "diploid")

# ===================================================================
# LIG3 (7 conditions)
# ===================================================================
message("\n========== LIG3 ==========")

message("--- p20250808 ---")
lig3_p08_ot <- run_check_on_target(p20250808_dirs, lig3_coord_str,
  summary_p20250808_filt$sample, type = "diploid")

message("--- p20250409 ---")
lig3_0409_ot <- run_check_on_target(p20250409_dir, lig3_coord_str,
  novaseq250806_summary_bj_rnp_filt$sample[novaseq250806_summary_bj_rnp_filt$condition == "BJ_rnp_lig3"],
  type = "diploid")

message("--- p20250924 ---")
lig3_0924_ot <- run_check_on_target(p20250924_dir, lig3_coord_str,
  summary_p20250924_filt %>% filter(condition == "bj_rnp_lig3") %>% pull(sample),
  type = "diploid")

message("--- HAP1 haploid ---")
lig3_hap1h_ot <- run_check_on_target(hap1_haploid_dir, lig3_coord_str,
  summary_hap1_haploid_filt$sample, type = "haploid")

message("--- HAP1 diploid ---")
lig3_hap1d_ot <- run_check_on_target(hap1_diploid_dir, lig3_coord_str,
  summary_hap1_diploid_filt$sample, type = "diploid")

message("--- Lig3_24hrs ---")
lig3_24hrs_ot <- run_check_on_target(
  file.path(new_base, "BJ_Cas9_RNP_Lig3_24hrs/data_filtered_10M"), lig3_coord_str,
  summaries_new_filt[["BJ_Cas9_RNP_Lig3_24hrs"]]$sample, type = "diploid")

message("--- Lig3_24wsBrdU ---")
lig3_brdu_ot <- run_check_on_target(
  file.path(new_base, "BJ_Cas9_RNP_Lig3_24wsBrdU/data_filtered_10M"), lig3_coord_str,
  summaries_new_filt[["BJ_Cas9_RNP_Lig3_24wsBrdU"]]$sample, type = "diploid")

# ===================================================================
# LIG4 (2 conditions)
# ===================================================================
message("\n========== LIG4 ==========")

message("--- Lig4_24hrs ---")
lig4_24hrs_ot <- run_check_on_target(
  file.path(new_base, "BJ_Cas9_RNP_Lig4_24hrs/data_filtered_10M"), lig4_coord_str,
  summaries_new_filt[["BJ_Cas9_RNP_Lig4_24hrs"]]$sample, type = "diploid")

message("--- Lig4_24wsBrdU ---")
lig4_brdu_ot <- run_check_on_target(
  file.path(new_base, "BJ_Cas9_RNP_Lig4_24wsBrdU/data_filtered_10M"), lig4_coord_str,
  summaries_new_filt[["BJ_Cas9_RNP_Lig4_24wsBrdU"]]$sample, type = "diploid")

# ===================================================================
# XRCC4 (2 conditions)
# ===================================================================
message("\n========== XRCC4 ==========")

message("--- Xrcc4_24hrs ---")
xrcc4_24hrs_ot <- run_check_on_target(
  file.path(new_base, "BJ_Cas9_RNP_Xrcc4_24hrs/data_filtered_10M"), xrcc4_coord_str,
  summaries_new_filt[["BJ_Cas9_RNP_Xrcc4_24hrs"]]$sample, type = "diploid")

message("--- Xrcc4_24wsBrdU ---")
xrcc4_brdu_ot <- run_check_on_target(
  file.path(new_base, "BJ_Cas9_RNP_Xrcc4_24wsBrdU/data_filtered_10M"), xrcc4_coord_str,
  summaries_new_filt[["BJ_Cas9_RNP_Xrcc4_24wsBrdU"]]$sample, type = "diploid")

# ===================================================================
# Combined summary table
# ===================================================================
message("\n=== Building summary table ===")

all_on_target <- rbind(
  cbind(gene = "XRCC1", condition = "p20250808_xrcc1_lig3_237", summarize_on_target(xrcc1_p08_ot)),
  cbind(gene = "XRCC1", condition = "p20250409_xrcc1",          summarize_on_target(xrcc1_0409_ot)),
  cbind(gene = "XRCC1", condition = "p20250924_xrcc1",          summarize_on_target(xrcc1_0924_ot)),
  cbind(gene = "XRCC1", condition = "HAP1_haploid",             summarize_on_target(xrcc1_hap1h_ot)),
  cbind(gene = "XRCC1", condition = "HAP1_diploid",             summarize_on_target(xrcc1_hap1d_ot)),
  cbind(gene = "XRCC1", condition = "Xrcc1_24hrs",              summarize_on_target(xrcc1_24hrs_ot)),
  cbind(gene = "XRCC1", condition = "Xrcc1_24wsBrdU",           summarize_on_target(xrcc1_brdu_ot)),
  cbind(gene = "LIG3",  condition = "p20250808_xrcc1_lig3_237", summarize_on_target(lig3_p08_ot)),
  cbind(gene = "LIG3",  condition = "p20250409_lig3",            summarize_on_target(lig3_0409_ot)),
  cbind(gene = "LIG3",  condition = "p20250924_lig3",            summarize_on_target(lig3_0924_ot)),
  cbind(gene = "LIG3",  condition = "HAP1_haploid",              summarize_on_target(lig3_hap1h_ot)),
  cbind(gene = "LIG3",  condition = "HAP1_diploid",              summarize_on_target(lig3_hap1d_ot)),
  cbind(gene = "LIG3",  condition = "Lig3_24hrs",                summarize_on_target(lig3_24hrs_ot)),
  cbind(gene = "LIG3",  condition = "Lig3_24wsBrdU",             summarize_on_target(lig3_brdu_ot)),
  cbind(gene = "LIG4",  condition = "Lig4_24hrs",                summarize_on_target(lig4_24hrs_ot)),
  cbind(gene = "LIG4",  condition = "Lig4_24wsBrdU",             summarize_on_target(lig4_brdu_ot)),
  cbind(gene = "XRCC4", condition = "Xrcc4_24hrs",               summarize_on_target(xrcc4_24hrs_ot)),
  cbind(gene = "XRCC4", condition = "Xrcc4_24wsBrdU",            summarize_on_target(xrcc4_brdu_ot))
)

all_on_target$on_target_rate <- round(all_on_target$on_target / all_on_target$total_cells, 4)

write.table(all_on_target, file.path(tab_dir, "on_target_breakpoint_summary.txt"),
            sep = "\t", row.names = FALSE, quote = FALSE)
message("Saved on_target_breakpoint_summary.txt")
print(all_on_target)

# ===================================================================
# Save per-cell results
# ===================================================================
save(xrcc1_p08_ot, xrcc1_0409_ot, xrcc1_0924_ot,
     xrcc1_hap1h_ot, xrcc1_hap1d_ot,
     xrcc1_24hrs_ot, xrcc1_brdu_ot,
     lig3_p08_ot, lig3_0409_ot, lig3_0924_ot,
     lig3_hap1h_ot, lig3_hap1d_ot,
     lig3_24hrs_ot, lig3_brdu_ot,
     lig4_24hrs_ot, lig4_brdu_ot,
     xrcc4_24hrs_ot, xrcc4_brdu_ot,
     all_on_target,
     file = "data/on_target_breakpoint.RData")
message("Saved data/on_target_breakpoint.RData")

# ===================================================================
# Plot (same layout as rescue_rate_all_conditions.pdf)
# ===================================================================
all_on_target$label <- paste0(all_on_target$gene, ": ", all_on_target$condition)
all_on_target$label <- factor(all_on_target$label, levels = rev(all_on_target$label))

p_ot <- ggplot(all_on_target, aes(x = label, y = on_target_rate, fill = gene)) +
  geom_col() +
  geom_text(aes(label = paste0(on_target, "/", total_cells)),
            hjust = -0.1, size = 2.5) +
  coord_flip() +
  facet_wrap(~gene, scales = "free_y", ncol = 1) +
  scale_y_continuous(expand = expansion(mult = c(0, 0.3)), labels = scales::percent) +
  theme_classic() +
  theme(strip.background = element_blank(),
        legend.position = "none") +
  xlab(NULL) + ylab("On-target rate (called + rescue)") +
  scale_fill_manual(values = c("XRCC1" = "#E41A1C", "LIG3" = "#377EB8",
                                "LIG4" = "#4DAF4A", "XRCC4" = "#984EA3"))

ggsave("on_target_break_point.pdf", p_ot, grDevices::cairo_pdf, fig_dir,
       height = 6, width = 8)
message("Saved on_target_break_point.pdf")

message("\nDone: 13_on_target_breakpoint.R")
```
