## Supplementary material for "Genome-wide mapping of Cas9-induced sister chromatid exchange across single and over 200 genomic targets": Cas9_RNP_scripts.zip: 13b_call_only_plot.R

```
#!/usr/bin/env Rscript
# 13b_call_only_plot.R
# Plot called-only on-target rate from existing on_target_breakpoint_summary.txt
# No re-computation needed

message("=== 13b_call_only_plot.R ===")
source("scripts/load_env.R")

all_ot <- read.table(file.path(tab_dir, "on_target_breakpoint_summary.txt"),
                     header = TRUE, sep = "\t")

# Verify: called_at_target + rescue_only == on_target
stopifnot(all(all_ot$called_at_target + all_ot$rescue_only == all_ot$on_target))
message("Verified: called_at_target + rescue_only == on_target for all rows")

# Add call-only rate
all_ot$call_rate <- round(all_ot$called_at_target / all_ot$total_cells, 4)

# ---- Called-only plot ----
all_ot$label <- paste0(all_ot$gene, ": ", all_ot$condition)
all_ot$label <- factor(all_ot$label, levels = rev(all_ot$label))

p_call <- ggplot(all_ot, aes(x = label, y = call_rate, fill = gene)) +
  geom_col() +
  geom_text(aes(label = paste0(called_at_target, "/", total_cells)),
            hjust = -0.1, size = 2.5) +
  coord_flip() +
  facet_wrap(~gene, scales = "free_y", ncol = 1) +
  scale_y_continuous(expand = expansion(mult = c(0, 0.3)), labels = scales::percent) +
  theme_classic() +
  theme(strip.background = element_blank(),
        legend.position = "none") +
  xlab(NULL) + ylab("Called break at target rate") +
  scale_fill_manual(values = c("XRCC1" = "#E41A1C", "LIG3" = "#377EB8",
                                "LIG4" = "#4DAF4A", "XRCC4" = "#984EA3"))

ggsave("call_on_target_breakpoint.pdf", p_call, grDevices::cairo_pdf, fig_dir,
       height = 6, width = 8)
message("Saved call_on_target_breakpoint.pdf")

# ---- Print comparison table ----
message("\nComparison: call-only vs rescue-only vs on_target (call+rescue)")
compare <- data.frame(
  gene = all_ot$gene,
  condition = all_ot$condition,
  total = all_ot$total_cells,
  called = all_ot$called_at_target,
  rescue_only = all_ot$rescue_only,
  on_target = all_ot$on_target,
  call_rate = all_ot$call_rate,
  on_target_rate = all_ot$on_target_rate
)
print(compare)

message("\nDone: 13b_call_only_plot.R")
```
