## Supplementary material for "Genome-wide mapping of Cas9-induced sister chromatid exchange across single and over 200 genomic targets": Cas9_RNP_scripts.zip: 13c_cell_id_verification.R

```
#!/usr/bin/env Rscript
# 13c_cell_id_verification.R
# List cell IDs for called vs rescue for Lig3_24hrs and Xrcc1_24hrs

message("=== 13c_cell_id_verification.R ===")
source("scripts/load_env.R")
load("data/on_target_breakpoint.RData")

# ---- Lig3_24hrs ----
message("\n========== Lig3_24hrs ==========")
message("Called at target: ", sum(lig3_24hrs_ot$called_at_target))
message("Rescue break: ", sum(lig3_24hrs_ot$rescue_break))
message("On target (call | rescue): ", sum(lig3_24hrs_ot$on_target))

lig3_out <- data.frame(
  ID = lig3_24hrs_ot$ID,
  called_at_target = lig3_24hrs_ot$called_at_target,
  rescue_break = lig3_24hrs_ot$rescue_break,
  on_target = lig3_24hrs_ot$on_target,
  category = ifelse(lig3_24hrs_ot$called_at_target & lig3_24hrs_ot$rescue_break, "both",
             ifelse(lig3_24hrs_ot$called_at_target, "call_only",
             ifelse(lig3_24hrs_ot$rescue_break, "rescue_only", "neither")))
)
lig3_out <- lig3_out[order(-lig3_out$on_target, lig3_out$category, lig3_out$ID), ]

message("\nLig3_24hrs category counts:")
print(table(lig3_out$category))

message("\nLig3_24hrs on-target cells:")
print(lig3_out[lig3_out$on_target, ], row.names = FALSE)

# ---- Xrcc1_24hrs ----
message("\n========== Xrcc1_24hrs ==========")
message("Called at target: ", sum(xrcc1_24hrs_ot$called_at_target))
message("Rescue break: ", sum(xrcc1_24hrs_ot$rescue_break))
message("On target (call | rescue): ", sum(xrcc1_24hrs_ot$on_target))

xrcc1_out <- data.frame(
  ID = xrcc1_24hrs_ot$ID,
  called_at_target = xrcc1_24hrs_ot$called_at_target,
  rescue_break = xrcc1_24hrs_ot$rescue_break,
  on_target = xrcc1_24hrs_ot$on_target,
  category = ifelse(xrcc1_24hrs_ot$called_at_target & xrcc1_24hrs_ot$rescue_break, "both",
             ifelse(xrcc1_24hrs_ot$called_at_target, "call_only",
             ifelse(xrcc1_24hrs_ot$rescue_break, "rescue_only", "neither")))
)
xrcc1_out <- xrcc1_out[order(-xrcc1_out$on_target, xrcc1_out$category, xrcc1_out$ID), ]

message("\nXrcc1_24hrs category counts:")
print(table(xrcc1_out$category))

message("\nXrcc1_24hrs on-target cells:")
print(xrcc1_out[xrcc1_out$on_target, ], row.names = FALSE)

# ---- Save combined table ----
lig3_out$gene <- "LIG3"
lig3_out$condition <- "Lig3_24hrs"
xrcc1_out$gene <- "XRCC1"
xrcc1_out$condition <- "Xrcc1_24hrs"

combined <- rbind(
  lig3_out[, c("gene", "condition", "ID", "called_at_target", "rescue_break", "on_target", "category")],
  xrcc1_out[, c("gene", "condition", "ID", "called_at_target", "rescue_break", "on_target", "category")]
)

write.table(combined, "data/call_and_rescue_cell_ID_verification.txt",
            sep = "\t", row.names = FALSE, quote = FALSE)
message("\nSaved data/call_and_rescue_cell_ID_verification.txt")

message("\nDone: 13c_cell_id_verification.R")
```
