## Supplementary material for "Genome-wide mapping of Cas9-induced sister chromatid exchange across single and over 200 genomic targets": Cas9_RNP_scripts.zip: 14b_237_enrichment_collect.R

```
#!/usr/bin/env Rscript
# 14b_237_enrichment_collect.R
# Collect results from 14_237_enrichment_on_target.R (all 5 conditions)
# Build summary table and comparison plots
#
# Prerequisites:
#   1. 08_237_pileup.R must be run first — it produces pileup_237.RData
#      which contains bp_filt_gr per condition. bp_filt_gr is the set of
#      filtered breakpoints AFTER removing BJ-5ta inversions and hg19
#      blacklist regions (filter_by_non_overlaps). We use
#      unique(bp_filt_gr$sample) to define the valid cell set.
#   2. 14_237_enrichment_on_target.R must be run for all 5 conditions —
#      it produces per-condition RDS files with per-cell per-site results
#      for ALL QC-passed cells (_0.08_2_75 filter).
#
# =====================================================================
# Cell selection for contingency tables:
# =====================================================================
# The enrichment RDS files contain data for ALL cells passing the
# _0.08_2_75 QC filter (background.estimate > 0 & <= 0.08,
# med.reads.per.MB > 2, wc.perc + uknw.perc < 75).
#
# For contingency tables we subset to cells with >= 1 filtered
# breakpoint genome-wide that is NOT in a BJ-5ta inversion or hg19
# blacklist region. This cell set comes from:
#   pileup_237.RData -> <condition>$bp_filt_gr -> unique($sample)
#
# These are clean sample IDs (e.g. "yi478_AATTGA.ATCGCGTTTTACTTA")
# without the .human.filt.bam suffix that appears in the enrichment
# RDS cell IDs. We strip the suffix before matching.
#
# Why not just use post_filt > 0 from the QC CSV?
#   post_filt is the breakpointR internal filtered break count BEFORE
#   BJ-5ta inversion and hg19 blacklist removal. A cell can have
#   post_filt > 0 but all its breaks fall in blacklisted regions,
#   leaving it with zero valid breaks. Using bp_filt_gr$sample
#   (post-blacklist) gives the correct cell set.
#
# Cell counts per condition (QC-passed -> bp_filt_gr):
#   wt237:   76 ->  74
#   d10a:   338 -> 330
#   h840a:  785 -> 757
#   24hrs:  228 -> 214
#   brdu:  2622 -> 2547 (expected, verify after brdu completes)
# =====================================================================
#
# Contingency table construction:
# =====================================================================
# DO NOT use the cont_* matrices stored in the RDS directly — early
# runs had a bug where Absent = n_cells^2 * n_regions - Present
# instead of the correct n_cells * n_regions - Present.
# The Present column is always correct. Recompute Absent as:
#   total  <- n_cells * total_regions   (total cell-site pairs)
#   Absent <- total - Present
# =====================================================================

message("=== 14b_237_enrichment_collect.R ===")
source("scripts/load_env.R")

fig_out <- "data/figures_10conditions/enrichment_re_analysis/"
tab_out <- "data/tables_10conditions/enrichment_re_analysis/"

# ---------------------------------------------------------------
# Load valid cell sets from pileup_237.RData
# ---------------------------------------------------------------
# Produced by 08_237_pileup.R. Contains bp_filt_gr per condition:
# filtered breakpoints after BJ-5ta inversion + hg19 blacklist removal.
# unique(bp_filt_gr$sample) = cells with >= 1 valid filtered breakpoint.
load("data/pileup_237.RData")

valid_cells <- list(
  p20250924_WT237      = unique(p20250924_237_wt$bp_filt_gr$sample),
  p20250924_d10a       = unique(p20250924_237_d10a$bp_filt_gr$sample),
  p20250924_h840a      = unique(p20250924_237_h840a$bp_filt_gr$sample),
  "237cuts_24hrs"       = unique(new_237_24hrs$bp_filt_gr$sample),
  "237cuts_24ws_BrdU"   = unique(new_237_brdu$bp_filt_gr$sample)
)

cond_names <- names(valid_cells)

# ---------------------------------------------------------------
# Strip .human.filt.bam / .hg19.filt.bam suffix from RDS cell IDs
# to match the clean IDs in bp_filt_gr$sample
# ---------------------------------------------------------------
strip_bam_suffix <- function(ids) {
  sub("[.](human|hg19)[.]filt[.]bam$", "", ids)
}

# ---------------------------------------------------------------
# Build contingency table from sample_counts_df, subset to valid cells
# ---------------------------------------------------------------
make_cont <- function(sc, roi_col, rand_col) {
  n_cells <- nrow(sc)
  total_regions <- sc$total_regions[1]   # 237
  total <- n_cells * total_regions
  roi_present  <- sum(sc[[roi_col]])
  rand_present <- sum(sc[[rand_col]])
  matrix(
    c(roi_present,  total - roi_present,
      rand_present, total - rand_present),
    nrow = 2, byrow = TRUE,
    dimnames = list(Region = c("Cut_sites", "Random_sites"),
                    Breakpoint = c("Present", "Absent")))
}

# ---------------------------------------------------------------
# Load enrichment RDS and subset to valid cells
# ---------------------------------------------------------------
all_results <- list()
for (cn in cond_names) {
  rds_path <- file.path(tab_out, paste0("c237_", cn, "_on_target.rds"))
  if (!file.exists(rds_path)) stop("Missing: ", rds_path, " -- run 14_237_enrichment_on_target.R first")
  all_results[[cn]] <- readRDS(rds_path)
  message("Loaded ", cn)
}

# ---- Summary table ----
summary_rows <- list()
for (i in seq_along(cond_names)) {
  cn  <- cond_names[i]
  res <- all_results[[cn]]
  sc  <- res$sample_counts_df

  # Subset to valid cells (bp_filt_gr from pileup_237.RData)
  sc$sample_clean <- strip_bam_suffix(sc$sample)
  sc_sub <- sc[sc$sample_clean %in% valid_cells[[cn]], ]

  n_all <- nrow(sc)
  n_valid <- nrow(sc_sub)
  message("  ", cn, ": ", n_all, " QC-passed -> ", n_valid, " with valid breaks")

  cont_call      <- make_cont(sc_sub, "roi_call",      "rand_call")
  cont_rescue    <- make_cont(sc_sub, "roi_rescue",    "rand_rescue")
  cont_on_target <- make_cont(sc_sub, "roi_on_target", "rand_on_target")

  ct_call   <- chisq.test(cont_call)
  ct_rescue <- chisq.test(cont_rescue)
  ct_ot     <- chisq.test(cont_on_target)

  summary_rows[[i]] <- data.frame(
    condition = cn,
    n_cells_qc = n_all,
    n_cells_valid = n_valid,
    call_CutSites_present     = cont_call["Cut_sites", "Present"],
    call_Random_present       = cont_call["Random_sites", "Present"],
    call_pvalue               = ct_call$p.value,
    rescue_CutSites_present   = cont_rescue["Cut_sites", "Present"],
    rescue_Random_present     = cont_rescue["Random_sites", "Present"],
    rescue_pvalue             = ct_rescue$p.value,
    on_target_CutSites_present = cont_on_target["Cut_sites", "Present"],
    on_target_Random_present   = cont_on_target["Random_sites", "Present"],
    on_target_pvalue           = ct_ot$p.value
  )
}
enrichment_summary <- dplyr::bind_rows(summary_rows)

write.table(enrichment_summary,
  file.path(tab_out, "enrichment_237_on_target_summary.txt"),
  sep = "\t", row.names = FALSE, quote = FALSE)
message("\nEnrichment summary:")
print(enrichment_summary)

# ---- P-value comparison plot ----
plot_df <- enrichment_summary %>%
  tidyr::pivot_longer(
    cols = c(call_pvalue, rescue_pvalue, on_target_pvalue),
    names_to = "method", values_to = "pvalue") %>%
  mutate(
    method = factor(method,
      levels = c("call_pvalue", "rescue_pvalue", "on_target_pvalue"),
      labels = c("Call only", "Rescue only", "Call + Rescue")),
    condition = factor(condition, levels = rev(cond_names)),
    neg_log10_p = -log10(pvalue))

p_pval <- ggplot(plot_df, aes(x = condition, y = neg_log10_p, fill = method)) +
  geom_col(position = position_dodge(width = 0.8), width = 0.7) +
  geom_hline(yintercept = -log10(0.05), linetype = "dashed", color = "red") +
  annotate("text", x = 0.5, y = -log10(0.05) + 0.3, label = "p=0.05",
           color = "red", hjust = 0, size = 3) +
  coord_flip() +
  theme_classic() +
  theme(legend.position = "bottom") +
  xlab(NULL) + ylab(expression(-log[10](p-value))) +
  ggtitle("237-guide enrichment: call vs rescue vs on-target") +
  scale_fill_manual(values = c("Call only" = "#4DAF4A",
                                "Rescue only" = "#377EB8",
                                "Call + Rescue" = "#E41A1C"))

ggsave("enrichment_237_on_target_pvalues.pdf", p_pval,
  grDevices::cairo_pdf, fig_out, height = 5, width = 8)
message("Saved enrichment_237_on_target_pvalues.pdf")

# ---- Enrichment ratio plot ----
ratio_df <- enrichment_summary %>%
  mutate(
    call_ratio      = call_CutSites_present / pmax(call_Random_present, 1),
    rescue_ratio    = rescue_CutSites_present / pmax(rescue_Random_present, 1),
    on_target_ratio = on_target_CutSites_present / pmax(on_target_Random_present, 1)
  ) %>%
  tidyr::pivot_longer(
    cols = c(call_ratio, rescue_ratio, on_target_ratio),
    names_to = "method", values_to = "ratio") %>%
  mutate(
    method = factor(method,
      levels = c("call_ratio", "rescue_ratio", "on_target_ratio"),
      labels = c("Call only", "Rescue only", "Call + Rescue")),
    condition = factor(condition, levels = rev(cond_names)))

p_ratio <- ggplot(ratio_df, aes(x = condition, y = ratio, fill = method)) +
  geom_col(position = position_dodge(width = 0.8), width = 0.7) +
  geom_hline(yintercept = 1, linetype = "dashed", color = "grey50") +
  coord_flip() +
  theme_classic() +
  theme(legend.position = "bottom") +
  xlab(NULL) + ylab("Enrichment ratio (cut sites / random sites)") +
  ggtitle("237-guide enrichment ratio") +
  scale_fill_manual(values = c("Call only" = "#4DAF4A",
                                "Rescue only" = "#377EB8",
                                "Call + Rescue" = "#E41A1C"))

ggsave("enrichment_237_on_target_ratio.pdf", p_ratio,
  grDevices::cairo_pdf, fig_out, height = 5, width = 8)
message("Saved enrichment_237_on_target_ratio.pdf")

# ===================================================================
# High-SCE (post_filt >= 8) results
# ===================================================================
# NOTE: High-SCE RDS files already contain only cells with
# post_filt >= cutoff. These cells by definition have post_filt >= 8,
# so they all have >= 1 filtered breakpoint and the blacklist filter
# only removes a handful (those whose breaks are all in blacklisted
# regions despite having >= 8 total). We still apply the bp_filt_gr
# filter for consistency.
message("\n===== High-SCE (post_filt >= 8) =====")
hs_results <- list()
hs_available <- TRUE
for (cn in cond_names) {
  rds_path <- file.path(tab_out, paste0("c237_", cn, "_on_target_high_sce_8.rds"))
  if (!file.exists(rds_path)) {
    message("Missing high-SCE result for ", cn, " -- skipping high-SCE summary")
    hs_available <- FALSE
    break
  }
  hs_results[[cn]] <- readRDS(rds_path)
  message("Loaded high-SCE: ", cn, " (", hs_results[[cn]]$n_high_sce, " cells)")
}

if (hs_available) {
  hs_rows <- list()
  for (i in seq_along(cond_names)) {
    cn  <- cond_names[i]
    res <- hs_results[[cn]]
    sc  <- res$sample_counts_df

    sc$sample_clean <- strip_bam_suffix(sc$sample)
    sc_sub <- sc[sc$sample_clean %in% valid_cells[[cn]], ]

    cont_call      <- make_cont(sc_sub, "roi_call",      "rand_call")
    cont_rescue    <- make_cont(sc_sub, "roi_rescue",    "rand_rescue")
    cont_on_target <- make_cont(sc_sub, "roi_on_target", "rand_on_target")

    ct_call   <- chisq.test(cont_call)
    ct_rescue <- chisq.test(cont_rescue)
    ct_ot     <- chisq.test(cont_on_target)

    hs_rows[[i]] <- data.frame(
      condition = cn,
      n_cells_all = res$n_all,
      n_cells_high_sce = nrow(sc),
      n_cells_valid = nrow(sc_sub),
      call_CutSites_present     = cont_call["Cut_sites", "Present"],
      call_Random_present       = cont_call["Random_sites", "Present"],
      call_pvalue               = ct_call$p.value,
      rescue_CutSites_present   = cont_rescue["Cut_sites", "Present"],
      rescue_Random_present     = cont_rescue["Random_sites", "Present"],
      rescue_pvalue             = ct_rescue$p.value,
      on_target_CutSites_present = cont_on_target["Cut_sites", "Present"],
      on_target_Random_present   = cont_on_target["Random_sites", "Present"],
      on_target_pvalue           = ct_ot$p.value
    )
  }
  hs_summary <- dplyr::bind_rows(hs_rows)

  write.table(hs_summary,
    file.path(tab_out, "enrichment_237_on_target_high_sce_8_summary.txt"),
    sep = "\t", row.names = FALSE, quote = FALSE)
  message("\nHigh-SCE enrichment summary:")
  print(hs_summary)

  # P-value comparison plot (high-SCE)
  hs_plot_df <- hs_summary %>%
    tidyr::pivot_longer(
      cols = c(call_pvalue, rescue_pvalue, on_target_pvalue),
      names_to = "method", values_to = "pvalue") %>%
    mutate(
      method = factor(method,
        levels = c("call_pvalue", "rescue_pvalue", "on_target_pvalue"),
        labels = c("Call only", "Rescue only", "Call + Rescue")),
      condition = factor(condition, levels = rev(cond_names)),
      neg_log10_p = -log10(pvalue))

  p_hs <- ggplot(hs_plot_df, aes(x = condition, y = neg_log10_p, fill = method)) +
    geom_col(position = position_dodge(width = 0.8), width = 0.7) +
    geom_hline(yintercept = -log10(0.05), linetype = "dashed", color = "red") +
    annotate("text", x = 0.5, y = -log10(0.05) + 0.3, label = "p=0.05",
             color = "red", hjust = 0, size = 3) +
    coord_flip() +
    theme_classic() +
    theme(legend.position = "bottom") +
    xlab(NULL) + ylab(expression(-log[10](p-value))) +
    ggtitle("237-guide enrichment (high-SCE, post_filt >= 8)") +
    scale_fill_manual(values = c("Call only" = "#4DAF4A",
                                  "Rescue only" = "#377EB8",
                                  "Call + Rescue" = "#E41A1C"))

  ggsave("enrichment_237_on_target_high_sce_8_pvalues.pdf", p_hs,
    grDevices::cairo_pdf, fig_out, height = 5, width = 8)
  message("Saved enrichment_237_on_target_high_sce_8_pvalues.pdf")
}

message("\nDone: 14b_237_enrichment_collect.R")
```
