## Supplementary material for "Genome-wide mapping of Cas9-induced sister chromatid exchange across single and over 200 genomic targets": Cas9_RNP_scripts.zip: 14c_237_enrichment_on_target_high_sce.R

```
#!/usr/bin/env Rscript
# 14c_237_enrichment_on_target_high_sce.R
# Same as 14_237_enrichment_on_target.R but restricted to cells with post_filt >= 8
#
# Usage: Rscript scripts/14c_237_enrichment_on_target_high_sce.R <condition>
#   condition: wt237, d10a, h840a, 24hrs, brdu
#
# Launch all 5 in parallel:
#   for c in wt237 d10a h840a 24hrs brdu; do
#     Rscript scripts/14c_237_enrichment_on_target_high_sce.R $c &
#   done
#   wait

args <- commandArgs(trailingOnly = TRUE)
if (length(args) == 0) stop("Provide condition: wt237, d10a, h840a, 24hrs, brdu [cutoff]")
cond_arg <- args[1]
cutoff <- if (length(args) >= 2) as.integer(args[2]) else 8
message("=== 14c_237_enrichment_on_target_high_sce.R === condition: ", cond_arg, ", cutoff: ", cutoff)

source("scripts/load_env.R")
load("data/summaries_earlier.RData")
load("data/summaries_new10.RData")
load("data/pileup_237.RData")

tab_out <- "data/tables_10conditions/enrichment_re_analysis/"

# ---------------------------------------------------------------
# Functions (same as 14_237_enrichment_on_target.R)
# ---------------------------------------------------------------
check_location_on_target <- function(path, coord, cent,
                                     type = c("diploid", "haploid"),
                                     ensembl = TRUE, assembly = "hg19",
                                     bin_width = 10000000, dist_from_gr = 3000000,
                                     stretch_amount = 6000000) {
  if (grepl("\\.RData$", path, ignore.case = TRUE)) {
    env <- new.env()
    load(path, envir = env)
    f <- ls(env)[1]
    dehcr_filt <- get(f, envir = env)
  } else {
    f <- load_from_rds(path)
    dehcr_filt <- get(f)
  }
  fragments <- dehcr_filt$fragments
  breaks    <- dehcr_filt$breaks

  target_regions <- plyranges::stretch(coord, stretch_amount)
  called_vec <- GenomicRanges::countOverlaps(target_regions, breaks) > 0

  coord_grl  <- split(coord)
  coord_arms <- purrr::map(coord_grl, \(x) trim(bins_of_interest(x,
    dist_from_gr = dist_from_gr, bin_width = bin_width)))

  df_out <- list()
  for (i in seq_along(coord_arms)) {
    L_R_states <- suppressWarnings(state_assignment_bins(coord_arms[[i]], fragments))
    L_R_states <- sort(L_R_states)
    new_states <- paste(L_R_states$states, collapse = "_")
    if (type == "haploid") {
      rescue_break <- new_states %in% c("ww_cc", "cc_ww")
    } else if (type == "diploid") {
      rescue_break <- new_states %in% c("ww_cc", "cc_ww",
        "ww_wc", "wc_ww", "cc_wc", "wc_cc")
    }
    df_out[[i]] <- data.frame(
      ID = dehcr_filt$ID,
      called_at_target = called_vec[i],
      rescue_break = rescue_break,
      on_target = called_vec[i] | rescue_break,
      tiled_states = new_states,
      coord_idx = i)
  }
  out <- dplyr::bind_rows(df_out)
  assertthat::assert_that(nrow(out) == length(coord),
    msg = "Not all coordinates are being returned.")
  return(out)
}

run_check_on_target <- function(data_folder, coord, samples,
                                type = "diploid", num_threads = 10,
                                bin_width = 10000000, dist_from_gr = 3000000,
                                stretch_amount = 6000000, filtered = TRUE) {
  data_files <- get_files_to_load(data_folder, samples, filtered = filtered)
  message("  Processing ", length(data_files), " cells x ", length(coord), " sites...")
  cent <- get_centromere_coordinates("hg19", FALSE, ensembl = TRUE)

  if (num_threads > 1) {
    pb <- txtProgressBar(max = length(data_files), style = 3)
    progress <- function(n) setTxtProgressBar(pb, n)
    cl <- parallel::makeCluster(num_threads)
    doSNOW::registerDoSNOW(cl)
    out_df <- foreach(f = data_files,
                      .packages = c("sciStrandR", "dplyr", "purrr", "plyranges",
                                    "GenomicRanges", "assertthat"),
                      .export = c("check_location_on_target",
                                  "bins_of_interest", "state_assignment_bins"),
                      .options.snow = list(progress = progress),
                      .combine = rbind) %dopar% {
      check_location_on_target(f, coord, cent = cent, type = type,
        bin_width = bin_width, dist_from_gr = dist_from_gr,
        stretch_amount = stretch_amount)
    }
    parallel::stopCluster(cl)
  } else {
    out_lst <- list()
    for (f in data_files) {
      out_lst[[f]] <- check_location_on_target(f, coord, cent = cent, type = type,
        bin_width = bin_width, dist_from_gr = dist_from_gr,
        stretch_amount = stretch_amount)
    }
    out_df <- dplyr::bind_rows(out_lst)
  }
  rownames(out_df) <- NULL
  return(out_df)
}

# ===================================================================
# Chromosome arms
# ===================================================================
cent_hg19 <- get_centromere_coordinates("hg19", FALSE, ensembl = TRUE)
arms <- gaps(cent_hg19)
arms <- arms[strand(arms) == "*"]
arms_filt <- arms[1:42]
seqlevels(arms_filt, pruning.mode = "coarse") <- seqlevels(arms_filt)[1:23]

# ===================================================================
# Build post_filt lookup and filter samples
# ===================================================================
read_new_qc <- function(csv_path) {
  df <- read.csv(csv_path)
  df$sample <- sub("^([^\\.]*\\.[^\\.]*).*", "\\1", df$.id)
  df_filt <- df[df$background.estimate > 0 & df$background.estimate <= 0.08 &
                df$med.reads.per.MB > 2 & (df$wc.perc + df$uknw.perc) < 75, ]
  df_filt[, c("sample", "post_filt")]
}

old_qc <- summary_p20250924_filt %>%
  filter(condition %in% c("bj_rnp_seq3_237", "bj_rnp_seq3_237_d10a", "bj_rnp_seq3_237_h840a")) %>%
  select(sample, condition, post_filt)

qc_24hrs <- read_new_qc(file.path(new_base_path, "BJ_Cas9_RNP_237cuts_24hrs_16hrs", "breakPoint_qc_summary_10M.csv"))
qc_brdu  <- read_new_qc(file.path(new_base_path, "BJ_Cas9_RNP_237cuts_24ws_40hrs_BrdU", "breakPoint_qc_summary_10M.csv"))

cond_map <- list(
  wt237 = list(
    name = "p20250924_WT237",
    folder = "/u/project/yeastyin/chovanec/projects/novaseq251007/BJ_RNP/human/breakpointR_diploid/data_filtered/",
    all_samples = summary_p20250924_filt %>% filter(condition == "bj_rnp_seq3_237") %>% pull(sample),
    qc_df = old_qc %>% filter(condition == "bj_rnp_seq3_237")
  ),
  d10a = list(
    name = "p20250924_d10a",
    folder = "/u/project/yeastyin/chovanec/projects/novaseq251007/BJ_RNP/human/breakpointR_diploid/data_filtered/",
    all_samples = summary_p20250924_filt %>% filter(condition == "bj_rnp_seq3_237_d10a") %>% pull(sample),
    qc_df = old_qc %>% filter(condition == "bj_rnp_seq3_237_d10a")
  ),
  h840a = list(
    name = "p20250924_h840a",
    folder = "/u/project/yeastyin/chovanec/projects/novaseq251007/BJ_RNP/human/breakpointR_diploid/data_filtered/",
    all_samples = summary_p20250924_filt %>% filter(condition == "bj_rnp_seq3_237_h840a") %>% pull(sample),
    qc_df = old_qc %>% filter(condition == "bj_rnp_seq3_237_h840a")
  ),
  "24hrs" = list(
    name = "237cuts_24hrs",
    folder = "/u/project/yeastyin/yeastyin/projects/Cas9_RNP/data/bpR_output/BJ_Cas9_RNP_237cuts_24hrs_16hrs/data_filtered_10M/",
    all_samples = summaries_new_filt[["BJ_Cas9_RNP_237cuts_24hrs_16hrs"]]$sample,
    qc_df = qc_24hrs
  ),
  brdu = list(
    name = "237cuts_24ws_BrdU",
    folder = "/u/project/yeastyin/yeastyin/projects/Cas9_RNP/data/bpR_output/BJ_Cas9_RNP_237cuts_24ws_40hrs_BrdU/data_filtered_10M/",
    all_samples = summaries_new_filt[["BJ_Cas9_RNP_237cuts_24ws_40hrs_BrdU"]]$sample,
    qc_df = qc_brdu
  )
)

if (!cond_arg %in% names(cond_map)) {
  stop("Unknown condition: ", cond_arg, ". Use one of: ", paste(names(cond_map), collapse = ", "))
}
cond <- cond_map[[cond_arg]]

# Filter to high_sce cells (post_filt >= cutoff)
high_sce_samples <- cond$qc_df$sample[cond$qc_df$post_filt >= cutoff]
samples_to_use <- intersect(cond$all_samples, high_sce_samples)

message("\n========== ", cond$name, " (high_sce >= ", cutoff, ") ==========")
message("  All QC samples: ", length(cond$all_samples))
message("  High-SCE samples (post_filt >= ", cutoff, "): ", length(samples_to_use))

if (length(samples_to_use) == 0) {
  message("  No high-SCE samples for this condition. Skipping.")
  quit(save = "no", status = 0)
}

# ===================================================================
# Generate random regions (same seed 3 as main analysis)
# ===================================================================
set.seed(3)
random_regions_237 <- regioneR::randomizeRegions(A = yy_237_gr, genome = arms_filt,
  allow.overlaps = FALSE, per.chromosome = TRUE)

# ===================================================================
# Run: ROI sites
# ===================================================================
message("\n--- Checking ROI sites (237 guides) ---")
roi_results <- run_check_on_target(cond$folder, coord = yy_237_gr,
  samples = samples_to_use, type = "diploid", num_threads = 10)

# ===================================================================
# Run: Random sites
# ===================================================================
message("\n--- Checking random sites (237 matched regions) ---")
rand_results <- run_check_on_target(cond$folder, coord = random_regions_237,
  samples = samples_to_use, type = "diploid", num_threads = 10)

# ===================================================================
# Build per-cell summary and contingency tables
# ===================================================================
roi_by_cell  <- split(roi_results, roi_results$ID)
rand_by_cell <- split(rand_results, rand_results$ID)
total_regions <- length(yy_237_gr)

sample_counts <- list()
for (s in seq_along(roi_by_cell)) {
  cell_id   <- names(roi_by_cell)[s]
  roi_cell  <- roi_by_cell[[s]]
  rand_cell <- rand_by_cell[[cell_id]]

  sample_counts[[s]] <- data.frame(
    sample = cell_id,
    total_regions = total_regions,
    roi_call       = sum(roi_cell$called_at_target),
    rand_call      = sum(rand_cell$called_at_target),
    roi_rescue     = sum(roi_cell$rescue_break),
    rand_rescue    = sum(rand_cell$rescue_break),
    roi_on_target  = sum(roi_cell$on_target),
    rand_on_target = sum(rand_cell$on_target)
  )
}
sample_counts_df <- dplyr::bind_rows(sample_counts)

make_cont <- function(roi_col, rand_col) {
  n_cells <- nrow(sample_counts_df)
  total <- total_regions * n_cells
  matrix(
    c(sum(sample_counts_df[[roi_col]]),
      total - sum(sample_counts_df[[roi_col]]),
      sum(sample_counts_df[[rand_col]]),
      total - sum(sample_counts_df[[rand_col]])),
    nrow = 2, byrow = TRUE,
    dimnames = list(Region = c("Cut_sites", "Random_sites"),
                    Breakpoint = c("Present", "Absent")))
}

cont_call      <- make_cont("roi_call", "rand_call")
cont_rescue    <- make_cont("roi_rescue", "rand_rescue")
cont_on_target <- make_cont("roi_on_target", "rand_on_target")

# Print results
message("\n  Call-only contingency:")
print(cont_call)
ct_call <- chisq.test(cont_call)
message("  p-value (call): ", ct_call$p.value)

message("\n  Rescue-only contingency:")
print(cont_rescue)
ct_rescue <- chisq.test(cont_rescue)
message("  p-value (rescue): ", ct_rescue$p.value)

message("\n  On-target (call+rescue) contingency:")
print(cont_on_target)
ct_ot <- chisq.test(cont_on_target)
message("  p-value (on_target): ", ct_ot$p.value)

# ===================================================================
# Save everything
# ===================================================================
# NOTE: The key data are roi_results, rand_results, and sample_counts_df.
# The cont_* matrices are convenience copies for quick inspection but
# should be recomputed from sample_counts_df when used for analysis —
# early runs saved wrong Absent values (n_cells^2 * n_regions bug in
# make_cont). Present values are always correct. Recompute Absent as:
#   total <- n_cells * total_regions; Absent <- total - Present
saveRDS(list(
  condition = cond$name,
  cutoff = cutoff,
  n_all = length(cond$all_samples),
  n_high_sce = length(samples_to_use),
  roi_results = roi_results,
  rand_results = rand_results,
  sample_counts_df = sample_counts_df,
  cont_call = cont_call,
  cont_rescue = cont_rescue,
  cont_on_target = cont_on_target,
  random_regions = random_regions_237,
  seed = 3
), file.path(tab_out, paste0("c237_", cond$name, "_on_target_high_sce_", cutoff, ".rds")))
message("Saved ", tab_out, "c237_", cond$name, "_on_target_high_sce_", cutoff, ".rds")

write.table(sample_counts_df,
  file.path(tab_out, paste0("per_cell_counts_", cond$name, "_high_sce_", cutoff, ".txt")),
  sep = "\t", row.names = FALSE, quote = FALSE)

write.table(cont_call,
  file.path(tab_out, paste0("contingency_", cond$name, "_call_high_sce_", cutoff, ".txt")),
  sep = "\t", quote = FALSE)
write.table(cont_rescue,
  file.path(tab_out, paste0("contingency_", cond$name, "_rescue_high_sce_", cutoff, ".txt")),
  sep = "\t", quote = FALSE)
write.table(cont_on_target,
  file.path(tab_out, paste0("contingency_", cond$name, "_on_target_high_sce_", cutoff, ".txt")),
  sep = "\t", quote = FALSE)

message("\nDone: 14c_237_enrichment_on_target_high_sce.R [", cond$name, "]")
```
