## Supplementary material for "Genome-wide mapping of Cas9-induced sister chromatid exchange across single and over 200 genomic targets": Cas9_RNP_scripts.zip: 14d_verify_rescue_match.R

```
#!/usr/bin/env Rscript
# 14d_verify_rescue_match.R
# Verify that saved RDS can reproduce old rescue contingency tables
# by subsetting to cells with >=1 filtered breakpoint (the old cell set).
#
# =====================================================================
# NOTE ON CELL SELECTION (IMPORTANT):
# =====================================================================
# The re-analysis scripts (14_*.R) use ALL QC-passed cells from
# summaries_new_filt / summary_p20250924_filt.
#
# The OLD enrichment scripts (09_*.R) used only cells present in
# bp_filt_grl — i.e., cells with at least 1 genome-wide filtered
# breakpoint. This is a subset of QC-passed cells: some QC-passed
# cells have zero filtered breakpoints genome-wide.
#
# Cell counts by condition:
#   WT237:  76 QC-passed, 74 with >=1 filtered break (2 extra)
#   24hrs: 228 QC-passed, 214 with >=1 filtered break (14 extra)
#   d10a:  338 QC-passed, 330 with >=1 filtered break (8 extra)
#   h840a: 785 QC-passed, 757 with >=1 filtered break (28 extra)
#
# When we subset new results to the old cell set, rescue Present
# values match the old contingency tables EXACTLY. This confirms:
#   1. The RDS data is saved correctly
#   2. The rescue_break computation is identical to the old scripts
#   3. The random regions (seed=3) are identical
#
# Going forward: we use ALL QC-passed cells (the larger set) for
# the re-analysis, since there is no reason to exclude cells that
# pass QC but happen to have zero genome-wide breakpoints.
# =====================================================================
#
# Usage: Rscript scripts/14d_verify_rescue_match.R

strip_bam_suffix <- function(ids) {
  sub("[.](human|hg19)[.]filt[.]bam$", "", ids)
}

# NOTE: Do NOT use dat$cont_* from the RDS — early runs saved wrong Absent
# values. Recompute from roi_results/rand_results (always correct).
# Absent = n_cells * total_regions - Present.
recompute_from_rds <- function(rds_path, old_cont_path, cell_subset = NULL) {
  dat <- readRDS(rds_path)
  roi  <- dat$roi_results
  rand <- dat$rand_results

  roi$ID_clean  <- strip_bam_suffix(roi$ID)
  rand$ID_clean <- strip_bam_suffix(rand$ID)

  if (!is.null(cell_subset)) {
    roi  <- roi[roi$ID_clean %in% cell_subset, ]
    rand <- rand[rand$ID_clean %in% cell_subset, ]
  }

  total_regions <- length(unique(roi$coord_idx))
  n_cells <- length(unique(roi$ID))
  total <- total_regions * n_cells

  roi_by_cell  <- split(roi, roi$ID)
  rand_by_cell <- split(rand, rand$ID)

  sample_counts <- list()
  for (s in seq_along(roi_by_cell)) {
    cell_id <- names(roi_by_cell)[s]
    rc <- roi_by_cell[[s]]
    rn <- rand_by_cell[[cell_id]]
    sample_counts[[s]] <- data.frame(
      roi_rescue     = sum(rc$rescue_break),
      rand_rescue    = sum(rn$rescue_break),
      roi_call       = sum(rc$called_at_target),
      rand_call      = sum(rn$called_at_target),
      roi_on_target  = sum(rc$on_target),
      rand_on_target = sum(rn$on_target)
    )
  }
  sc <- dplyr::bind_rows(sample_counts)

  make_cont <- function(roi_col, rand_col) {
    matrix(
      c(sum(sc[[roi_col]]),   total - sum(sc[[roi_col]]),
        sum(sc[[rand_col]]),  total - sum(sc[[rand_col]])),
      nrow = 2, byrow = TRUE,
      dimnames = list(Region = c("Cut_sites", "Random_sites"),
                      Breakpoint = c("Present", "Absent")))
  }

  cont_rescue    <- make_cont("roi_rescue", "rand_rescue")
  cont_call      <- make_cont("roi_call", "rand_call")
  cont_on_target <- make_cont("roi_on_target", "rand_on_target")

  message("\n  N cells: ", n_cells, " | N regions: ", total_regions, " | Total: ", total)
  message("  Rescue contingency:")
  print(cont_rescue)

  if (file.exists(old_cont_path)) {
    old <- as.matrix(read.table(old_cont_path, sep = "\t", header = TRUE))
    message("  Old rescue contingency:")
    print(old)
    if (all(cont_rescue[, "Present"] == old[, "Present"])) {
      message("  >>> MATCH: rescue Present values match old results!")
    } else {
      message("  >>> MISMATCH")
      message("    New: ", paste(cont_rescue[, "Present"], collapse = ", "))
      message("    Old: ", paste(old[, "Present"], collapse = ", "))
    }
  }

  message("\n  Call-only contingency:")
  print(cont_call)
  message("  On-target (call+rescue) contingency:")
  print(cont_on_target)

  return(list(cont_rescue = cont_rescue, cont_call = cont_call,
              cont_on_target = cont_on_target, n_cells = n_cells))
}

# ---------------------------------------------------------------
# Conditions to verify
# ---------------------------------------------------------------
conds <- list(
  list(name = "wt237",
       rds = "data/c237_p20250924_WT237_on_target.rds",
       old_cont = "data/tables_10conditions/contingency_c237_wt_rescue.txt",
       old_rds = "data/c237_wt_cont_tab_resc.rds"),
  list(name = "24hrs",
       rds = "data/c237_237cuts_24hrs_on_target.rds",
       old_cont = "data/tables_10conditions/contingency_c237_24hrs_rescue.txt",
       old_rds = "data/c237_24hrs_cont_tab_resc.rds"),
  list(name = "d10a",
       rds = "data/c237_p20250924_d10a_on_target.rds",
       old_cont = "data/tables_10conditions/contingency_c237_d10a_rescue.txt",
       old_rds = "data/c237_d10a_cont_tab_resc.rds"),
  list(name = "h840a",
       rds = "data/c237_p20250924_h840a_on_target.rds",
       old_cont = "data/tables_10conditions/contingency_c237_h840a_rescue.txt",
       old_rds = "data/c237_h840a_cont_tab_resc.rds"),
  list(name = "brdu",
       rds = "data/c237_237cuts_24ws_BrdU_on_target.rds",
       old_cont = "data/tables_10conditions/contingency_c237_brdu_rescue.txt",
       old_rds = "data/c237_brdu_cont_tab_resc.rds")
)

for (cc in conds) {
  if (!file.exists(cc$rds)) {
    message("\n=== ", cc$name, " === RDS not found: ", cc$rds, " -- skipping")
    next
  }
  message("\n=============================================")
  message("=== ", cc$name, " ===")
  message("=============================================")

  message("\n--- All QC-passed cells ---")
  res_all <- recompute_from_rds(cc$rds, cc$old_cont)

  if (file.exists(cc$old_rds)) {
    old_dat <- readRDS(cc$old_rds)
    old_cells <- if ("sample_counts_df" %in% names(old_dat))
      unique(old_dat$sample_counts_df$sample) else NULL
    if (!is.null(old_cells)) {
      message("\n--- Subset to old cell set (", length(old_cells), " cells) ---")
      res_sub <- recompute_from_rds(cc$rds, cc$old_cont, cell_subset = old_cells)
    }
  }
}

message("\nDone: 14d_verify_rescue_match.R")
```
