## Supplementary material for "Genome-wide mapping of Cas9-induced sister chromatid exchange across single and over 200 genomic targets": Cas9_RNP_scripts.zip: pairwise_sisterness.cpp

```
// [[Rcpp::depends(RcppParallel)]]
#include <Rcpp.h>
#include <RcppParallel.h>
#include <cmath>
using namespace Rcpp;
using namespace RcppParallel;

struct SisternessWorker : public Worker {
  const RMatrix<double> score;
  const RMatrix<double> abs_score;
  RMatrix<double> out;
  double wc_thresh;
  int b;

  SisternessWorker(const NumericMatrix& score,
                   const NumericMatrix& abs_score,
                   NumericMatrix& out,
                   double wc_thresh)
    : score(score), abs_score(abs_score), out(out),
      wc_thresh(wc_thresh), b(score.ncol()) {}

  void operator()(std::size_t begin, std::size_t end) {
    int n = score.nrow();
    for (std::size_t i = begin; i < end; i++) {
      for (int j = i + 1; j < n; j++) {
        double total = 0.0;
        int count = 0;
        for (int k = 0; k < b; k++) {
          double ai = abs_score(i, k);
          double aj = abs_score(j, k);
          double mx = (ai > aj) ? ai : aj;
          if (mx > wc_thresh) {
            total += std::abs(score(i, k) + score(j, k));
            count++;
          }
        }
        double val = (count > 0) ? total / count : 0.0;
        out(i, j) = val;
        out(j, i) = val;
      }
    }
  }
};

// [[Rcpp::export]]
NumericMatrix pairwise_sisterness_cpp(NumericMatrix score_mat, double wc_thresh) {
  int n = score_mat.nrow();
  int b = score_mat.ncol();
  NumericMatrix out(n, n);

  // Pre-compute abs scores
  NumericMatrix abs_score(n, b);
  for (int i = 0; i < n; i++)
    for (int j = 0; j < b; j++)
      abs_score(i, j) = std::abs(score_mat(i, j));

  SisternessWorker worker(score_mat, abs_score, out, wc_thresh);
  parallelFor(0, n - 1, worker);

  return out;
}
```
