## Supplementary material for "Genome-wide mapping of Cas9-induced sister chromatid exchange across single and over 200 genomic targets": Cas9_RNP_scripts.zip: plot_sister_pair_haplotype.R

```
#!/usr/bin/env Rscript
# =============================================================================
# plot_sister_pair_haplotype.R
# =============================================================================
#
# PURPOSE:
#   Plot strand-seq read-count ideograms for a putative sister cell pair,
#   showing both the unsplit (all reads) and haplotype-resolved views.
#   237 CRISPR guide positions are marked as red ticks on each track.
#
# OUTPUT LAYOUT (5 panels, top to bottom):
#   1. Cell A — all reads (unsplit)
#   2. Cell A — Haplotype 1 only
#   3. Cell A — Haplotype 2 only
#   4. Cell B — all reads (unsplit)
#   5. Cell B — Haplotype 1 only
#   6. Cell B — Haplotype 2 only
#
# HAPLOTYPE SPLITTING:
#   Uses sciStrandR:::split_bam_by_haplotype_helper() to assign reads to
#   haplotypes based on BJ-5ta phased SNVs. Only reads overlapping a phased
#   heterozygous SNV can be assigned; the rest remain unassigned. This means
#   haplotype tracks have much lower coverage than the unsplit track.
#
#   Haplotype SNV files (tab-separated, columns: chr/chrom, pos, base):
#     hap1: BJ-5ta_allchr_allphasingoptimizations_hap1.txt.gz
#     hap2: BJ-5ta_allchr_allphasingoptimizations_hap2.txt.gz
#   These contain ~2M phased SNVs per haplotype for the BJ-5ta cell line.
#
# READ PROCESSING:
#   sciStrandR::bam_unique_reads() reads the BAM, deduplicates (offset=3,
#   ignore soft-clipping), and marks duplicates. We keep only unique reads
#   (duplicate == FALSE), then subset by haplotype read names.
#
# PLOTTING STYLE:
#   Matches breakpointR plotBreakpoints() top panel: 200kb bins, Watson reads
#   (+ strand) up in sandybrown, Crick reads (- strand) down in
#   paleturquoise4. Chromosomes arranged linearly with cumulative coordinates.
#
# PAIR CONFIGURATION:
#   Edit the `pairs` list below to plot different pairs/chromosomes.
#   The script caches haplotype-split fragments as RDS files so re-runs
#   are fast.
#
# PREREQUISITES:
#   - conda activate sci_bpR_claude
#   - sciStrandR package (from /u/project/yeastyin/chovanec/software/sciStrandR)
#   - pileup_237.RData in Cas9_RNP/analysis/data/ (for guide positions)
#
# USAGE:
#   cd /u/project/yeastyin/yeastyin/projects/Cas9_RNP/analysis
#   conda activate sci_bpR_claude
#   Rscript /u/project/yeastyin/yeastyin/projects/reciprocal_daughter_cell/analysis/strandseq_sister_cell/plot_sister_pair_haplotype.R
#
# =============================================================================

suppressPackageStartupMessages({
  library(sciStrandR)
  library(GenomicRanges)
  library(GenomeInfoDb)
  library(ggplot2)
  library(cowplot)
  library(vroom)
})

# =============================================================================
# Configuration
# =============================================================================
bam_base   <- "/u/project/yeastyin/yeastyin/projects/Cas9_RNP/data/BJ_Cas9_RNP_237cuts_24ws_40hrs_BrdU"
hap1_file  <- "/u/project/yeastyin/chovanec/projects/BJ-5ta/BJ-5ta_allchr_allphasingoptimizations_hap1.txt.gz"
hap2_file  <- "/u/project/yeastyin/chovanec/projects/BJ-5ta/BJ-5ta_allchr_allphasingoptimizations_hap2.txt.gz"
cache_dir  <- "/u/project/yeastyin/yeastyin/projects/Cas9_RNP/analysis/data/tables_10conditions/enrichment_re_analysis"
fig_dir    <- "/u/project/yeastyin/yeastyin/projects/Cas9_RNP/analysis/data/figures_10conditions/enrichment_re_analysis"
pileup_rdata <- "/u/project/yeastyin/yeastyin/projects/Cas9_RNP/analysis/data/pileup_237.RData"

# ---- Pairs to plot ----
# Each entry: pair number, sisterness score, two cell IDs, chromosomes to show.
# Cell IDs are the BAM file basenames without the .hg19.filt.bam suffix.
# These pairs come from BJ_Cas9_RNP_237cuts_24ws_40hrs_BrdU_top20_sister_pairs.csv.
pairs <- list(
  list(pair = 6, sisterness = 0.298,
       cell_a = "yi561_GTCCTA.GAGTCTTTAGAACTA",
       cell_b = "yi563_TTCTCC.ATGCTGCGGTAAAGG",
       chroms = c("2", "3", "4", "6", "7", "12", "15")),
  list(pair = 7, sisterness = 0.350,
       cell_a = "yi560_CGCTTG.ATGCTGCGCTGCATG",
       cell_b = "yi560_TTCTCC.ATGCTGCGGATTGTG",
       chroms = c("1", "2", "3", "8", "11", "16", "19", "X")),
  list(pair = 11, sisterness = 0.421,
       cell_a = "yi562_AGGATG.ATGCTGCGACGTCGT",
       cell_b = "yi563_AGGATG.GAGTCTTTCATTGCG",
       chroms = c("2", "6", "7", "X")),
  list(pair = 13, sisterness = 0.422,
       cell_a = "yi560_TATTCT.GAGTCTTTGACGGGG",
       cell_b = "yi561_TTCTCC.ATGCTGCGAAATGGA",
       chroms = c("1", "4", "10", "11", "12", "13", "16", "19"))
)

# =============================================================================
# Load haplotype SNVs (shared across all pairs)
# =============================================================================
message("Loading haplotype SNVs...")
hap1_df <- vroom(hap1_file, show_col_types = FALSE)
hap2_df <- vroom(hap2_file, show_col_types = FALSE)
hap1_df$haplotype <- "hap1"
hap2_df$haplotype <- "hap2"
colnames(hap1_df) <- c("chrom", "pos", "base", "haplotype")
colnames(hap2_df) <- c("chrom", "pos", "base", "haplotype")
message("  hap1: ", nrow(hap1_df), " SNVs, hap2: ", nrow(hap2_df), " SNVs")

# =============================================================================
# Helper: get haplotype-split + unsplit fragments for one cell
# =============================================================================
# Returns list(all, hap1, hap2) of GRanges with deduplicated unique reads.
# Results are cached as RDS to avoid re-processing.
get_cell_fragments <- function(cell_id) {
  rds_path <- file.path(cache_dir, paste0(cell_id, "_hap_frags.rds"))

  if (file.exists(rds_path)) {
    message("  Loading cached: ", basename(rds_path))
    return(readRDS(rds_path))
  }

  bam_path <- file.path(bam_base, paste0(cell_id, ".hg19.filt.bam"))
  message("  Splitting ", cell_id, " by haplotype...")

  # Step 1: assign reads to haplotypes via phased SNVs
  res <- sciStrandR:::split_bam_by_haplotype_helper(
    hap1_df, hap2_df, bam_path,
    paired_end = TRUE, min_mapq = 20,
    filter_alt_alignments = TRUE,
    write_mixed = FALSE, ensembl = TRUE,
    read_names_only = FALSE)

  hap1_rn <- res$read_names_df$read_names[res$read_names_df$haplotype == "hap1"]
  hap2_rn <- res$read_names_df$read_names[res$read_names_df$haplotype == "hap2"]

  # Step 2: read BAM, deduplicate, keep unique reads
  all_reads <- bam_unique_reads(bam_path, mapq = 20, offset = 3,
                                 ignore_soft_clipping = TRUE,
                                 mark_only = TRUE)
  unqs <- all_reads[all_reads$duplicate == FALSE]

  # Step 3: subset unique reads by haplotype
  frags_hap1 <- unqs[names(unqs) %in% hap1_rn]
  frags_hap2 <- unqs[names(unqs) %in% hap2_rn]

  message("    hap1=", length(frags_hap1), " hap2=", length(frags_hap2),
          " total_unique=", length(unqs))

  result <- list(all = unqs, hap1 = frags_hap1, hap2 = frags_hap2)
  saveRDS(result, rds_path)
  message("    Saved: ", basename(rds_path))
  result
}

# =============================================================================
# Helper: bin reads into 200kb windows and compute cumulative genome coords
# =============================================================================
make_read_df <- function(frags, chroms, sl, cum_offset) {
  frags <- keepSeqlevels(frags, chroms, pruning.mode = "coarse")
  seqlengths(frags) <- sl

  bins <- unlist(tileGenome(sl, tilewidth = 200000))

  # Watson = plus strand (orange, up), Crick = minus strand (green, down)
  W <- countOverlaps(bins, frags[strand(frags) == "+"])
  C <- countOverlaps(bins, frags[strand(frags) == "-"])

  df <- as.data.frame(bins)
  df$Watsonreads <- W
  df$Crickreads <- C
  df$midpoint <- cum_offset[as.character(df$seqnames)] +
    (df$start + df$end) / 2

  # Cap outliers at 99.9th percentile (same as breakpointR)
  W_cap <- quantile(df$Watsonreads, 0.999)
  C_cap <- quantile(df$Crickreads, 0.999)
  df$Watsonreads[df$Watsonreads >= W_cap] <- W_cap
  df$Crickreads[df$Crickreads >= C_cap] <- C_cap
  df$mCrickreads <- -df$Crickreads
  df
}

# =============================================================================
# Helper: plot one track (breakpointR top-panel style)
# =============================================================================
plot_track <- function(df, title_text, guides, chr_lines_df, chr_label_pos,
                       chroms, y_lim, show_x = FALSE) {
  my_theme <- theme(
    legend.position = "none",
    panel.background = element_blank(),
    panel.border = element_blank(),
    panel.grid.major = element_blank(),
    panel.grid.minor = element_blank(),
    plot.background = element_blank()
  )

  p <- ggplot(df) +
    geom_linerange(aes(ymin = 0, ymax = mCrickreads, x = midpoint),
                   color = "paleturquoise4", linewidth = 0.2) +
    geom_linerange(aes(ymin = 0, ymax = Watsonreads, x = midpoint),
                   color = "sandybrown", linewidth = 0.2) +
    geom_vline(data = chr_lines_df, aes(xintercept = y),
               color = "black", linewidth = 0.3) +
    geom_point(data = guides, aes(x = genome_pos, y = 0),
               color = "red", shape = 124, size = 3) +
    scale_x_continuous(breaks = chr_label_pos,
                       labels = chroms,
                       expand = c(0, 0)) +
    coord_cartesian(ylim = y_lim) +
    xlab(NULL) + ylab("Reads") +
    my_theme +
    ggtitle(title_text)

  if (!show_x) {
    p <- p + theme(axis.text.x = element_blank(),
                   axis.ticks.x = element_blank())
  }
  p
}

# =============================================================================
# Load 237 guide positions
# =============================================================================
load(pileup_rdata)  # loads yy_237_gr
guides_all <- as.data.frame(yy_237_gr)

# =============================================================================
# Generate plots for each pair
# =============================================================================
for (pp in pairs) {
  chroms <- pp$chroms
  message("\n=== Pair ", pp$pair, " ===")

  # Get fragments (cached after first run)
  frags_a <- get_cell_fragments(pp$cell_a)
  frags_b <- get_cell_fragments(pp$cell_b)

  # Chromosome coordinate system
  sl <- seqlengths(frags_a$all)
  sl <- sl[chroms]
  cum_offset <- c(0, cumsum(as.numeric(sl[-length(sl)])))
  names(cum_offset) <- chroms
  chr_lines <- cumsum(as.numeric(sl[-length(sl)]))
  chr_label_pos <- cum_offset + sl / 2
  chr_lines_df <- data.frame(y = chr_lines)

  # Guide positions in genome coordinates
  guides <- guides_all[guides_all$seqnames %in% chroms, ]
  guides$genome_pos <- cum_offset[as.character(guides$seqnames)] +
    (guides$start + guides$end) / 2

  # Bin reads for all 6 tracks
  df_a_all <- make_read_df(frags_a$all,  chroms, sl, cum_offset)
  df_a_h1  <- make_read_df(frags_a$hap1, chroms, sl, cum_offset)
  df_a_h2  <- make_read_df(frags_a$hap2, chroms, sl, cum_offset)
  df_b_all <- make_read_df(frags_b$all,  chroms, sl, cum_offset)
  df_b_h1  <- make_read_df(frags_b$hap1, chroms, sl, cum_offset)
  df_b_h2  <- make_read_df(frags_b$hap2, chroms, sl, cum_offset)

  # Consistent y-axis: use max across unsplit tracks for unsplit,
  # separate scale for haplotype tracks (much fewer reads)
  y_max_all <- max(c(df_a_all$Watsonreads, df_a_all$Crickreads,
                     df_b_all$Watsonreads, df_b_all$Crickreads))
  y_lim_all <- c(-y_max_all, y_max_all) * 1.05

  y_max_hap <- max(c(df_a_h1$Watsonreads, df_a_h1$Crickreads,
                     df_a_h2$Watsonreads, df_a_h2$Crickreads,
                     df_b_h1$Watsonreads, df_b_h1$Crickreads,
                     df_b_h2$Watsonreads, df_b_h2$Crickreads))
  y_lim_hap <- c(-y_max_hap, y_max_hap) * 1.05

  # Build 6 tracks
  p1 <- plot_track(df_a_all, paste0(pp$cell_a, " — all reads"),
                   guides, chr_lines_df, chr_label_pos, chroms, y_lim_all)
  p2 <- plot_track(df_a_h1, paste0(pp$cell_a, " — Haplotype 1"),
                   guides, chr_lines_df, chr_label_pos, chroms, y_lim_hap)
  p3 <- plot_track(df_a_h2, paste0(pp$cell_a, " — Haplotype 2"),
                   guides, chr_lines_df, chr_label_pos, chroms, y_lim_hap)
  p4 <- plot_track(df_b_all, paste0(pp$cell_b, " — all reads"),
                   guides, chr_lines_df, chr_label_pos, chroms, y_lim_all)
  p5 <- plot_track(df_b_h1, paste0(pp$cell_b, " — Haplotype 1"),
                   guides, chr_lines_df, chr_label_pos, chroms, y_lim_hap)
  p6 <- plot_track(df_b_h2, paste0(pp$cell_b, " — Haplotype 2"),
                   guides, chr_lines_df, chr_label_pos, chroms, y_lim_hap,
                   show_x = TRUE)

  title <- cowplot::ggdraw() +
    cowplot::draw_label(
      sprintf("Sister pair %d (sisterness = %.3f) — haplotype-resolved — red ticks = 237 guides",
              pp$pair, pp$sisterness),
      fontface = "bold", size = 12)

  p_final <- cowplot::plot_grid(
    title, p1, p2, p3, p4, p5, p6,
    ncol = 1, align = "v",
    rel_heights = c(0.05, 1, 0.7, 0.7, 1, 0.7, 0.7))

  chr_tag <- paste(chroms, collapse = "_")
  out_pdf <- file.path(fig_dir,
    sprintf("sister_pair%d_haplotype_chr%s.pdf", pp$pair, chr_tag))

  grDevices::cairo_pdf(out_pdf, width = 14, height = 14)
  print(p_final)
  dev.off()
  message("Saved: ", out_pdf)
}

message("\nDone: plot_sister_pair_haplotype.R")
```
