## Supplementary material for "Genome-wide mapping of Cas9-induced sister chromatid exchange across single and over 200 genomic targets": Cas9_RNP_scripts.zip: run_bj_xrcc1.R

```
library(GenomicRanges)
library(tidyverse)
library(patchwork)
library(ggrastr)
library(parallel)
library(breakpointR)
library(Rcpp)
library(RcppParallel)

plot_font_base <- 14
theme_set(theme_bw(base_size = plot_font_base) +
            theme(plot.title = element_text(hjust = .5)) +
            theme(strip.background = element_rect(fill = 'black'),
                  strip.text = element_text(color = 'white')))

# --- Constants ---
bg_threshold <- 0.12
wc_thresh <- 0.5
bin_width <- 1e6
chroms <- c(as.character(1:22), "X")

# --- Helper functions ---
load_breakpoints <- function(f) {
  env <- new.env()
  load(f, envir = env)
  env$breakpoints
}

extract_metrics <- function(f) {
  bp <- load_breakpoints(f)
  tibble(
    file = f,
    cell = bp$ID,
    n_fragments = length(bp$fragments),
    n_breaks = length(bp$breaks),
    background = bp$lib.metrics["background.estimate"],
    reads_per_mb = bp$lib.metrics["med.reads.per.MB"],
    perc_coverage = bp$lib.metrics["perc.coverage"]
  )
}

get_score_vector <- function(f, bins, min_reads = 5) {
  bp <- load_breakpoints(f)
  frags <- bp$fragments
  frags <- keepSeqlevels(frags, seqlevels(bins), pruning.mode = "coarse")
  W <- countOverlaps(bins, frags[strand(frags) == "-"])
  C <- countOverlaps(bins, frags[strand(frags) == "+"])
  total <- W + C
  ifelse(total >= min_reads, (W - C) / total, 0)
}

# --- Compile Rcpp ---
sourceCpp("pairwise_sisterness.cpp")

# --- Data directory ---
data_dir <- "/u/project/yeastyin/yeastyin/projects/reciprocal_daughter_cell/data/bpR_output/BJ_XRCC1_15H/data"

# --- Genome bins (hg19, 1Mb) ---
any_rdata <- list.files(data_dir, pattern = "\\.RData$", full.names = TRUE)[1]
bp_tmp <- load_breakpoints(any_rdata)
sl <- seqlengths(bp_tmp$fragments)
sl <- sl[names(sl) %in% chroms]
bins <- unlist(tileGenome(sl, tilewidth = bin_width))
bins <- keepSeqlevels(bins, chroms, pruning.mode = "coarse")
bin_labels <- paste0(as.character(seqnames(bins)), ":", round(start(bins) / 1e6), "Mb")
rm(bp_tmp)
cat("Genome bins:", length(bins), "x", bin_width / 1e6, "Mb\n")

# --- Run pipeline ---
files <- list.files(data_dir, pattern = "\\.RData$", full.names = TRUE)
cat("BJ_XRCC1_15H — total cells:", length(files), "\n")

metrics <- map_dfr(files, extract_metrics) |>
  mutate(pass_qc = background <= bg_threshold & background != 0)
passing <- metrics |> filter(pass_qc)
cat("BJ_XRCC1_15H — passing QC:", nrow(passing), "/", nrow(metrics), "\n")

score_list <- mclapply(passing$file, get_score_vector, bins = bins,
                       min_reads = 4, mc.cores = 12)
score_mat <- do.call(rbind, score_list)
rownames(score_mat) <- passing$cell
cat("BJ_XRCC1_15H — score matrix:", nrow(score_mat), "x", ncol(score_mat), "\n")

t0 <- proc.time()
sist <- pairwise_sisterness_cpp(score_mat, wc_thresh)
elapsed <- (proc.time() - t0)[3]
dimnames(sist) <- list(passing$cell, passing$cell)
cat("BJ_XRCC1_15H — sisterness computed in", round(elapsed, 1), "sec\n")

diag(sist) <- NA
upr <- which(upper.tri(sist), arr.ind = TRUE)
vals <- sist[upr]
n_top <- min(100, length(vals))
top_idx <- order(vals)[seq_len(n_top)]

top_pairs <- tibble(
  cell_a = passing$cell[upr[top_idx, 1]],
  cell_b = passing$cell[upr[top_idx, 2]],
  sisterness = vals[top_idx]
)
cat("BJ_XRCC1_15H — top sisterness range:", round(min(top_pairs$sisterness), 4),
    "to", round(max(top_pairs$sisterness), 4), "\n")

# Save results
write_csv(top_pairs, "bj_xrcc1_15h_top100_sister_pairs.csv")
cat("Saved bj_xrcc1_15h_top100_sister_pairs.csv\n")

# PDF
cell_to_file <- setNames(passing$file, passing$cell)
pdf("bj_xrcc1_15h_top100_sister_pairs.pdf", width = 14, height = 8)
for (k in seq_len(nrow(top_pairs))) {
  bp_a <- load_breakpoints(cell_to_file[top_pairs$cell_a[k]])
  bp_b <- load_breakpoints(cell_to_file[top_pairs$cell_b[k]])
  pa <- plotBreakpoints(bp_a)[[1]]
  pb <- plotBreakpoints(bp_b)[[1]]
  ttl <- sprintf("Pair %d (sisterness = %.3f)", k, top_pairs$sisterness[k])
  print(pa / pb + plot_annotation(title = ttl))
}
dev.off()
cat("Saved bj_xrcc1_15h_top100_sister_pairs.pdf\n")
cat("Done!\n")
```
